# The faculty-to-faculty mentorship experience: a survey on challenges and recommendations for improvements

**DOI:** 10.1101/2022.10.17.512624

**Authors:** Sarvenaz Sarabipour, Natalie M Niemi, Steven J Burgess, Christopher T Smith, Alexandre W Bisson Filho, Ahmed Ibrahim, Kelly Clark

## Abstract

Faculty at research institutions play a central role in advancing knowledge and careers, as well as promoting the well-being of students and colleagues in research environments. Faculty members must balance a host of activities - such as performing research, teaching, sourcing funds, administrative and service duties - with their roles as educators and mentors. Mentorship from experienced peers has been touted as critical for enabling these myriad roles to allow faculty development, career progression, and satisfaction. However, there is little information available on who supports faculty and best ways to structure a faculty mentorship program for early- and mid-career academics. Furthermore, the extent to which mentorship and mentoring programs have been offered to faculty members has not been widely characterized. It is also unclear what challenges faculty receiving mentorship may face and which aspects could be further improved. In the interest of advocating for increased and enhanced faculty mentoring and mentoring programs, we surveyed faculty (i.e., group leaders) around the world to gather data on whether and how they receive mentoring from peers, senior researchers, informal mentoring programs, or formal mentoring programs at their institutions. We received responses from 457 early- and mid-career faculty and found that a substantial portion of respondents either reported having no mentor or a lack of a formal mentoring scheme. Qualitative responses on the quality of mentorship revealed that the most common complaints regarding mentorship included lack of mentor availability, unsatisfactory commitment to mentorship, and non-specific or non-actionable advice. Our findings further identified key mentorship elements desired by faculty mentees. Based on these suggestions, we identify a need for training for faculty mentors as well as strategies for individual mentors, departments, and institutions for funding and design of more intentional and supportive mentorship programs for early- and mid-career faculty.

## Introduction

Mentorship plays an important role in academic success by aiding researchers’ well-being and career development. For academic faculty, mentorship can lead to tangible benefits including higher career satisfaction (DeCastro et al., 2014), increased sense of self-efficacy (Feldman et al., 2010; Garman et al., 2001), an expanded professional network (Morzinski and Fisher, 2002), greater likelihood in obtaining funding (Palepu et al., 1998), an increased number of publications (Risner et al., 2020; Sambunjak et al., 2009), more time spent on research (Levinson et al., 1991), a shorter period to tenure (Morrison et al., 2014), and improved retention in academia (Cox, 1997). Additionally, mentorship can assist early career faculty to adjust to new and demanding expectations in roles for which they may have little training such as managing budgets, writing grants, balancing research, mentorship and teaching loads and navigating departmental politics. As few postdoctoral researchers receive training in all of these skills, mentors can assist in easing the transition into the faculty role through their advising, experience, and wisdom gained in these areas. Despite these benefits, access to mentors and quality of mentorship may not be equitable, optimal, or tailored to mentee needs, leading to variable mentorship experiences (Sambunjak et al., 2006; Straus et al., 2013). Such variability can limit potential benefits to mentors and mentees, and increasing access to effective mentors could improve the well-being and overall performance of faculty. A report by the United States National Academies of Science Engineering and Medicine (NASEM) defined mentorship as “a professional, working alliance in which individuals work together over time to support the personal and professional growth, development, and success of the relational partners through the provision of career and psychosocial support” (National Academies of Sciences, 2019). While providing a baseline definition, the NASEM report did not examine the quality of mentorship received by faculty.

Although mentees are responsible for seeking and managing mentorship interactions, mentors play an important role in the life and careers of researchers across career stages. Mentoring has historically been viewed as a dyad between an experienced and a less experienced individual (Kram, 1983), but can exist in a variety of forms, such as networks (DeCastro et al., 2013), peer-groups (Fleming et al., 2015; Moss et al., 2008; Thomas et al., 2015), group mentoring (Pololi and Knight, 2005; Pololi and Evans, 2015) or distance mentoring (Lewellen-Williams et al., 2006). Importantly, mentoring can also occur on a formal (i.e., within the context of an official mentorship scheme) or informal basis (i.e., an independently sourced advisor not part of an official mentorship scheme) (Law et al., 2014). In order to understand who supports faculty, in which specific ways faculty mentorship interactions fail, and what can be improved in faculty-to-faculty mentoring experiences in research environments, it is necessary to understand what factors contribute to successful and to less desirable mentorship outcomes.

Researchers in the science of mentorship have identified determinants of positive mentoring relationships (Pfund et al., 2016; Sambunjak et al., 2010; Straus et al., 2013), producing some recommendations on how to implement them at the institutional level (Law et al., 2014; Nick et al., 2012). This leads to the realization that mentoring is a skill that can be taught and learned (Pfund et al., 2006), aided by the development of curricula (Pfund et al., 2013), as well as tools for practice (Handelsman et al., 2005) and assessment (Anderson et al., 2012; Berk et al., 2005). However, many faculty may not have mentors, be unaware of the advantages of having a mentor, or lack access to pool of able or willing mentors - issues that may be more pronounced for women or marginalized communities (Beech et al., 2013; Carapinha et al., 2016). This lack of mentors is thought to have a negative impact on the success and retention of faculty members in academia (Tabak and Collins, 2011). To address this, a number of institutions have established and openly documented formal faculty programs to increase access to mentoring (Beech et al., 2013; Buch et al., 2011; Fountain and Newcomer, 2016; Johnson et al., 2010; Kashiwagi et al., 2013; Pololi et al., 2002; Sorkness et al., 2017; Thorndyke et al., 2006; Wingard et al., 2004).

Understanding mentoring needs for diverse individuals, along with designing and implementing effective interventions, requires an understanding of the current status in order to focus efforts. Systematic reviews and longitudinal studies have tracked the success of faculty mentoring programs in some cases, but most studies tend to be limited to individual disciplines, institutes, or regions, with the majority of faculty mentoring programs to date located within the United States (“Columbia University Guide to Best Practices in Faculty Mentoring: A Roadmap for Departments, Schools, Mentors and Mentees,” n.d.; Feldman, 2017; Law et al., 2014). This gap in knowledge on faculty mentoring experiences worldwide is problematic, as the benefits of mentoring are believed to be universal, and it is unclear to what extent mentoring is being effectively deployed to support faculty development. Research institutions need broader information and studies that compare the differences in needs of faculty mentees and action of mentors, departments and institutions.

In specific disciplines, such longitudinal studies are still sparse and disconnected. Academics commonly discuss mentorship initiatives for students and trainees, but it would be valuable to know who supports faculty, and particularly how junior faculty are mentored early in their career. To obtain an overview of mentoring experiences in research environments, we conducted a survey of early and mid-career faculty across disciplines to gauge mentee perceptions of mentor effectiveness, collecting responses from individual faculty across six continents. We implemented quantitative and qualitative questions and received responses on a broad range of mentoring characteristics. The results reveal that faculty mentoring experiences are not homogeneous and that there remains a need for implementing constructive mentorship initiatives and relationships that are tailored to individual faculty members.

## Results

### Faculty Demographics

To understand how faculty perceive their professional mentorship relationships, we surveyed group leaders worldwide to assess mentorship practices and their efficacy. The terminology differs across countries, for the purpose of this study faculty and group leader are used interchangeably and include lecturers, assistant or associate professor level faculty, with or without tenure. The survey was distributed from March 2019 to March 2020 and contained both scaled-response and open-ended questions. We received responses from 457 faculty from 48 countries (Figure 1A). By continent, 37% of responses originated from North America, 36% from Europe, 12% from Latin America, 8% from Asia, 5% from Oceania, and 2% from Africa. Almost one third of our responses were from faculty working in the United States (35%) followed by the United Kingdom (16%), Argentina (8%), Australia (5%), Spain (5%), and Germany (3%). Faculty from Canada, China, Chile, France, Malaysia, Belgium, the Netherlands, India, Sweden, Japan, Denmark, Croatia, Panama and Finland comprised 1-2% of our responses each, with other countries comprising the other 10% of total responses. The majority of respondents (64%) were in the 35-45 age group (Figure 3D). Ninety five percent of respondents operated their lab at an academic institution (Figure 1B). Fifty four percent of respondents performed basic research in life and biomedical sciences, 12% in physical and mathematical sciences, computer science or engineering, 10% in social and behavioral sciences and humanities, 9% in environmental sciences and field research, 8% in clinical and medical research and 7% in translational research (Figure 1C). The respondents to our survey were relatively evenly distributed across self-identified genders, with 45% identifying as male, 53% as female, and 2% identified as non-binary or preferred to not to disclose this information (Figure 1D). Sixty eight percent of respondents were assistant professors (or equivalent rank) while 22% were associate professors (or equivalent rank), 6% were full professors and 4% identified as independent research fellows (Figure 2A). The survey asked respondents at senior career stages (i.e., full professors) to participate with regards to mentorship they received at early and mid-career stages. Collectively, these responses constitute, to our knowledge, the largest and broadest dataset on faculty mentoring practices and their effectiveness worldwide.

**Figure 1.**
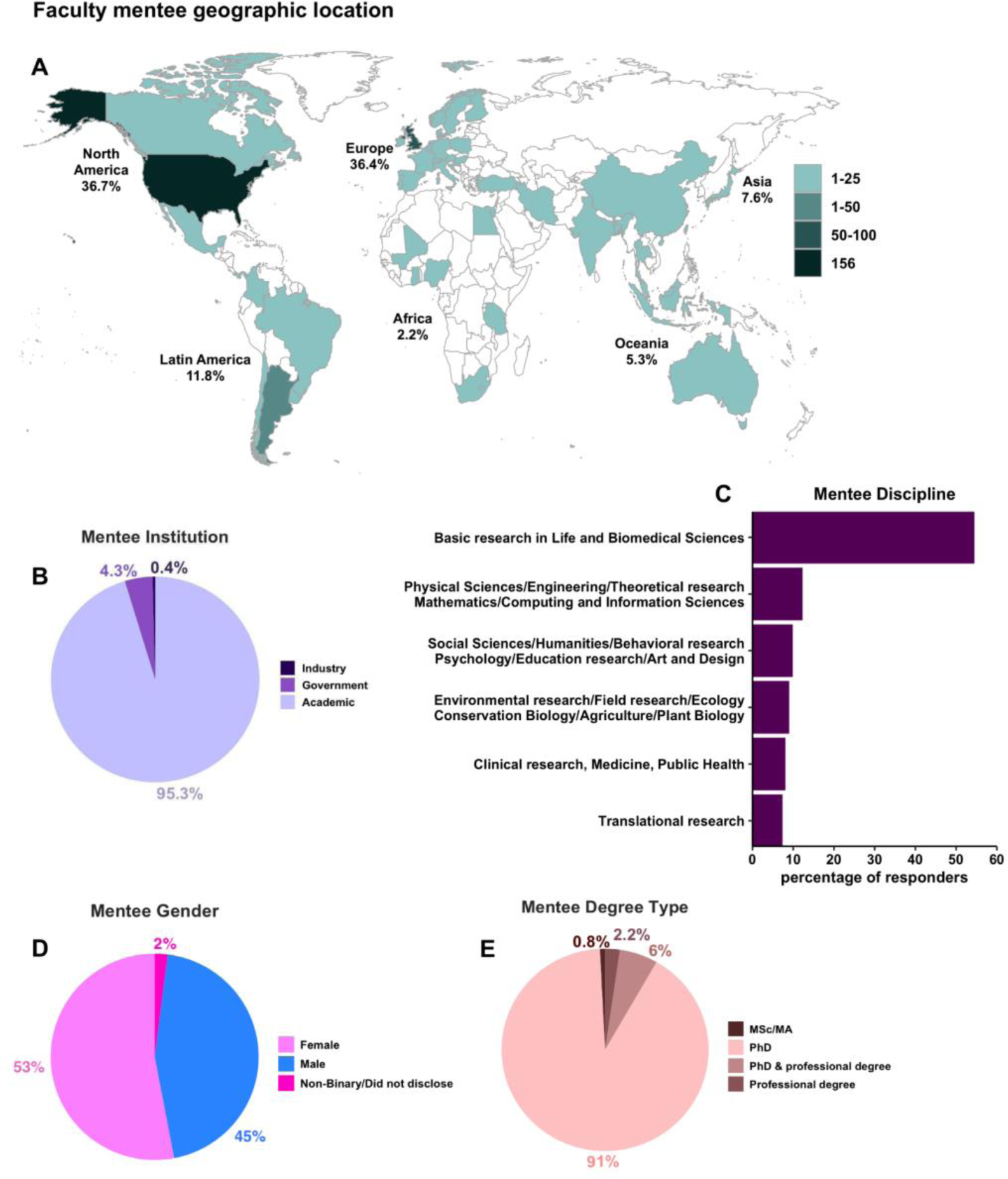
Demographics of mentees. Distribution of survey respondents by (A) country of research (where institutional affiliation was based) for the 457 early- and mid-career faculty (Table S1), (B) Type of mentee research institution (Table S2), (C) Mentee scientific discipline spanned basic research in life and biomedical sciences (54.3%), physical sciences, theoretical research, engineering, mathematics (12.1%), social sciences, humanities, psychology, education research, art and design (9.7%), environmental research, field research, ecology, agriculture, conservation biology, plant biology (8.8%), clinical research, medicine, public health (7.9%), and translational research (7.2%) (Table S3), (D) Mentee gender distribution (Table S4), (E) The type of advanced degree faculty mentee held: doctoral degree (PhD), professional degree (e.g., MD, DDS, RD, PT, PharmD, etc.), both PhD and professional degree (MD/PhD, MD/MPH, PharmD/MS, etc.) or Master’s degree (Science, Arts, Humanities) (Table S5), All responses were self-identified.

**Figure 2.**
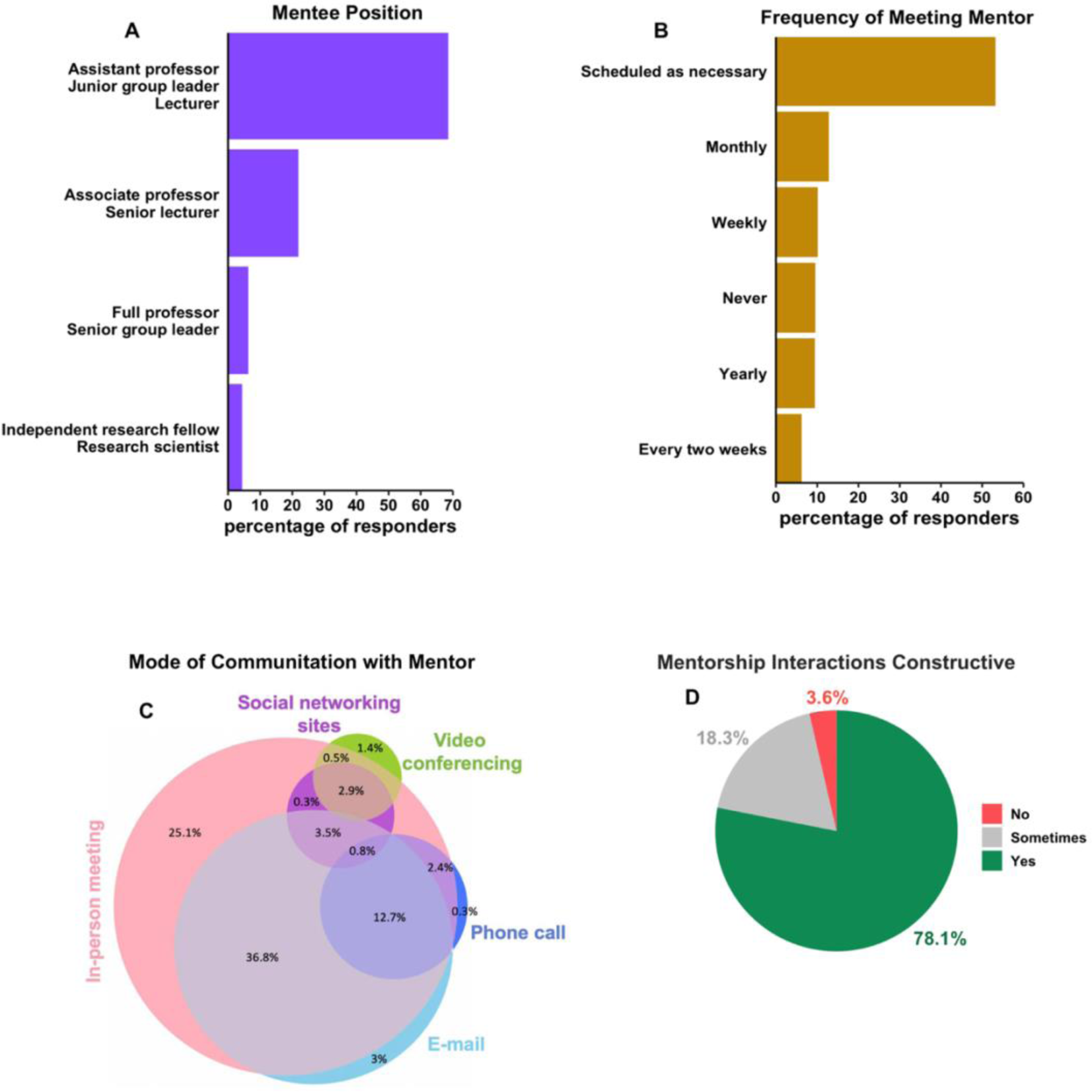
Characteristics of mentee-mentor interactions. (A) Mentee academic positions spanned assistant professors (68.3%) including tenure-track and non-tenure-track assistant professors, research professors, lecturers (a term commonly used in United Kingdom), junior group leaders (a term mostly used in Europe), associate professors (with or without tenure) (21.6%), independent research fellows and research scientists (4.1%) and full professors (6%) (Table S6). The survey asked respondents at senior career stages (i.e., full professors) to respond to the survey questions with regards to mentorship they received at early and mid-career stages, (B) Mentee-mentor meeting frequency was: scheduled as necessary (53%), monthly (12.6%), never (9.3%), weekly (9.9%), yearly (9.1%) and every two weeks (6%) (Table S7), (C) Faculty mentee format of communication with mentor. 30% of respondents only used one of the following communication modes: in-person (face-to-face) meeting, e-mail, phone call, video/web conferencing, chat/asynchronous interactions via social networking sites (such as Slack), while the rest of respondents (70%) used combinations of these formats (Table S9), (D) Quality of mentee-mentor interactions (Table S10), All responses were self-identified.

**Figure 3.**
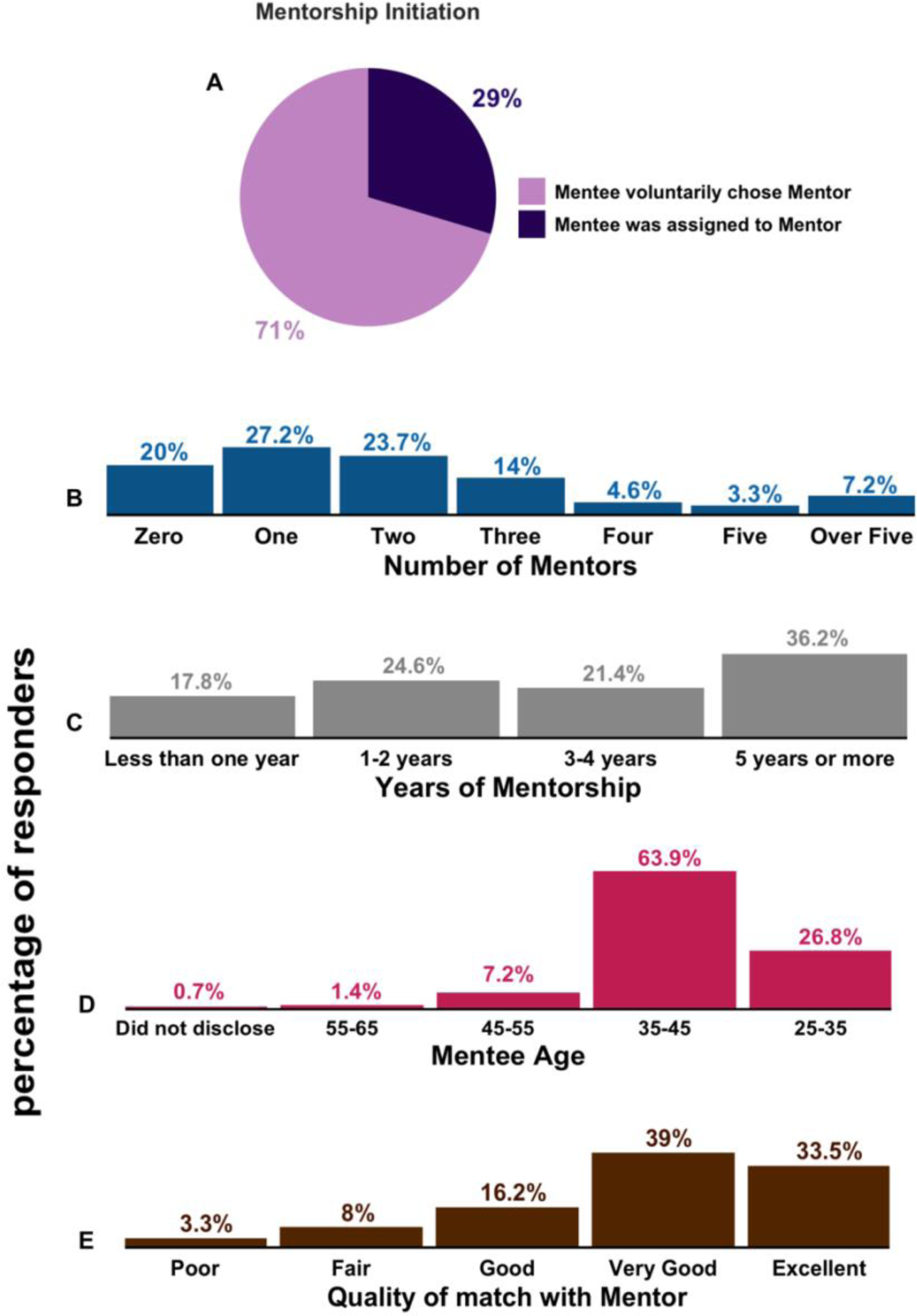
Mentee and mentorship characteristics. (A) Mentorship initiation mode (Table S8), (B) Number of Mentors (Table S12), (C) Years of working with the mentor(s) (Table S13), (D) Mentee age (Table S14), (E) Mentee assessment on the quality of match with mentor (Table S15).

### Characteristics of mentorship interactions

To understand how faculty mentorship is practiced across institutions and disciplines, we queried if mentorship was taking place and, if so, how mentoring relationships typically functioned among our respondents. Further, fifty three percent of respondents scheduled their meetings with mentors when necessary (Figure 2B). The majority of faculty mentees (70%) used a combination of formats to receive mentorship, suggesting that the sole use of face-to-face meetings is not the only option for sustaining mentoring relationships (Figure 2C). Overall, 78% of respondents found their interactions with their mentor constructive (Figure 2D), but women more commonly found their mentoring relationship less constructive than men (Figure 5C). The majority of mentees chose their faculty mentors voluntarily (71%) while a minority had a mentor assigned by their department (29%) (Figure 3A). While the majority of respondents indicated that they were receiving some mentorship, a considerable fraction (∼20%) of respondents reported not having a mentor, 27% had one mentor and 53% had two or more mentors (Figure 3B). Over 80% of respondents worked with their mentor for at least one year (Figure 3C). Consistently, 18% of respondents with mentors had received less than one year of mentorship, while 36% had received 5 or more years of mentorship (with the rest of respondents reporting an intermediate time frame of mentorship). Only 11% of mentees described the quality of match with their mentor as poor-to-fair, while 89% regarded their mentoring experience as good-to-excellent (Figure 3E).

### Impact of mentorship initiation mode

Analysis of responses by mentorship initiation format showed that faculty mentees who had an assigned mentor met with similar frequently with their mentor compared to faculty who voluntarily chose their mentor (Figure 4A, S4a, p=n.s.). Faculty mentees who had an assigned mentor found their mentorship interactions less constructive (Figure 4B, S4b, p<0.01), and were less satisfied with the mentorship match (Figure 4C, S4c, p<0.001), which remained consistent across genders (Figure S3b) compared to those who chose their mentor voluntarily.

**Figure 4.**
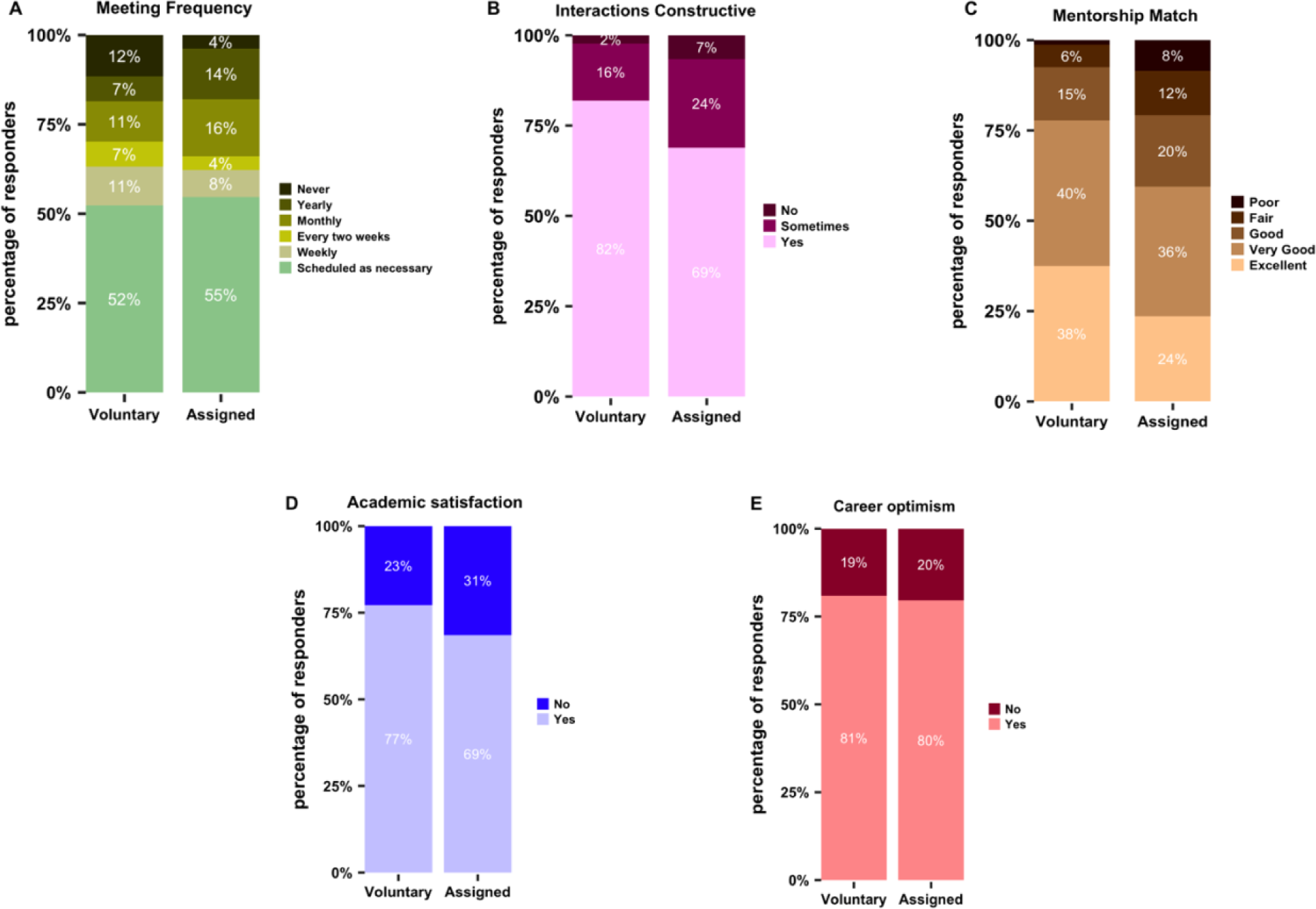
Mentorship features and quality by mentorship initiation mode. (A) Faculty mentee-mentor meeting frequency (Table S7b), (B) Quality of interactions with mentor (Table S10b), (C) Mentorship quality (Table S15a), (D) Mentee satisfaction on their current research program (Table S22a), (E) Optimism about their future research and position as an independent investigator by mentorship initiation mode (also see Figure S2c, d) (Table 23a). Data analysis for panels A, B, C excludes survey respondents with "0" number of mentors.

**Figure 5.**
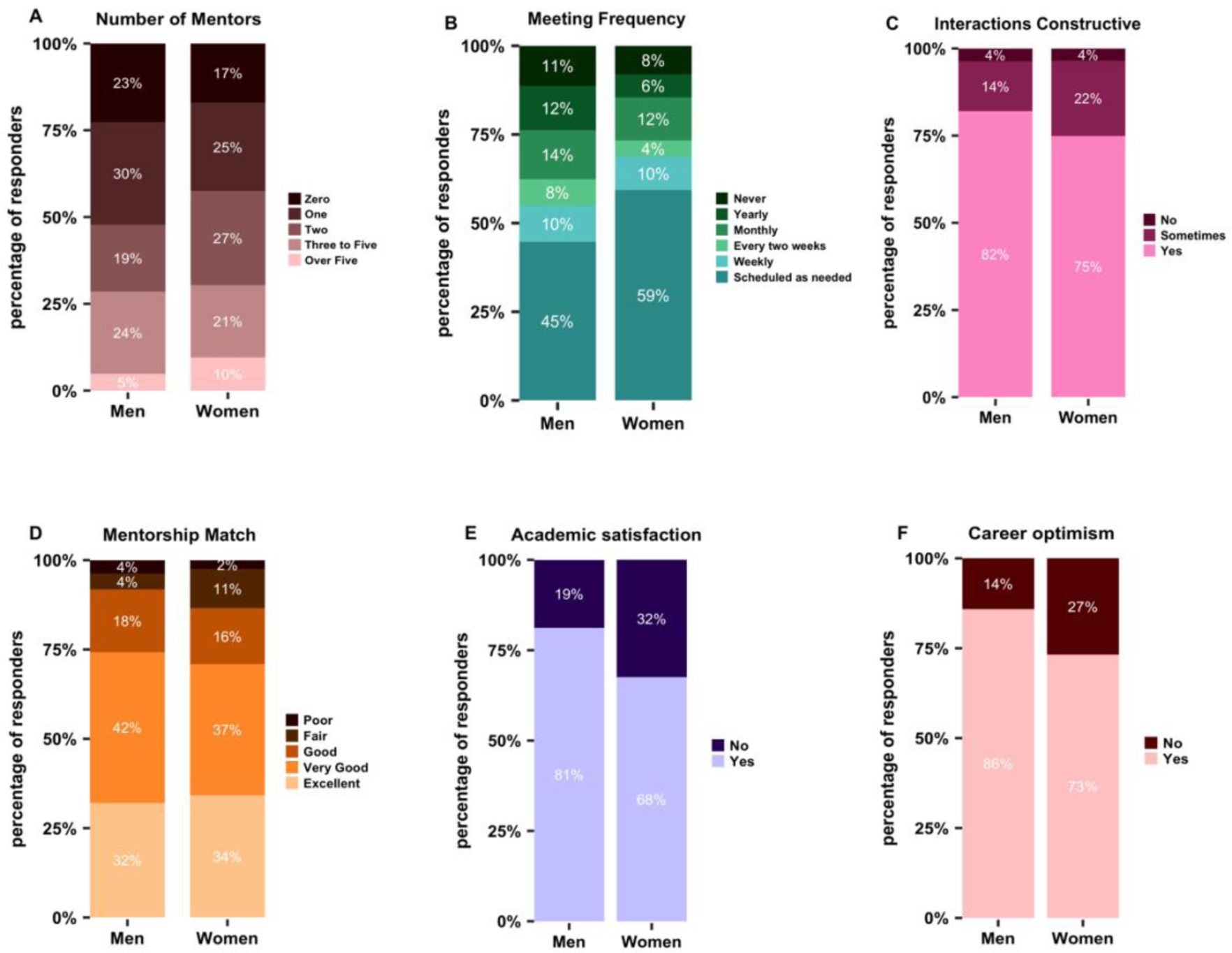
Mentorship features and quality by gender. (A) Number of mentors (Table S12), (B) Faculty mentee-mentor meeting frequency (Table S7), (C) Quality of interactions with mentor (Table S10), (D) Mentorship quality (Table S15), (E) Mentee satisfaction with current research program (Table S22), (F) Mentee optimism about their future research and position as an independent investigator by gender (Table S23) (Figure S2a, b). Data analysis for panels B, C, D, E, F excludes survey respondents with "0" number of mentors.

### Influence of gender on mentorship satisfaction

We assessed the survey results for possible trends on gender disparities in access to and satisfaction with mentors. 23% of men and 17% of women lacked mentors, with no significant difference between genders (Figure 5A, S4d, p=n.s.) and 58% of women having more than one mentor compared to 48% of men. 11% of Men reported few to no meetings with their mentor as compared to 8% of women, with no significant difference between genders (Figure 5B, S4e, p=0.0593, p=n.s.). Further, 26% of women and 18% of men reported their mentorship interactions to be “not constructive” or “sometimes constructive”, with no significant difference between genders (Figure 5C, S4f, p=n.s.). Men and women had similar neutral to negative and fair-to-poor mentorship match (Figure 5D, S4g, p=n.s.) and 24% of women than men and 17% of men did not maintain contact with former mentors (Figure 7D-E). The majority of mentees of both genders reported having a male mentor (71% of men and 60% of women) (Figure 7D-E) consistent with persistent faculty gender gap across disciplines (Casad et al., 2022; Li et al., 2021; Spoon et al., 2023).

**Figure 6.**
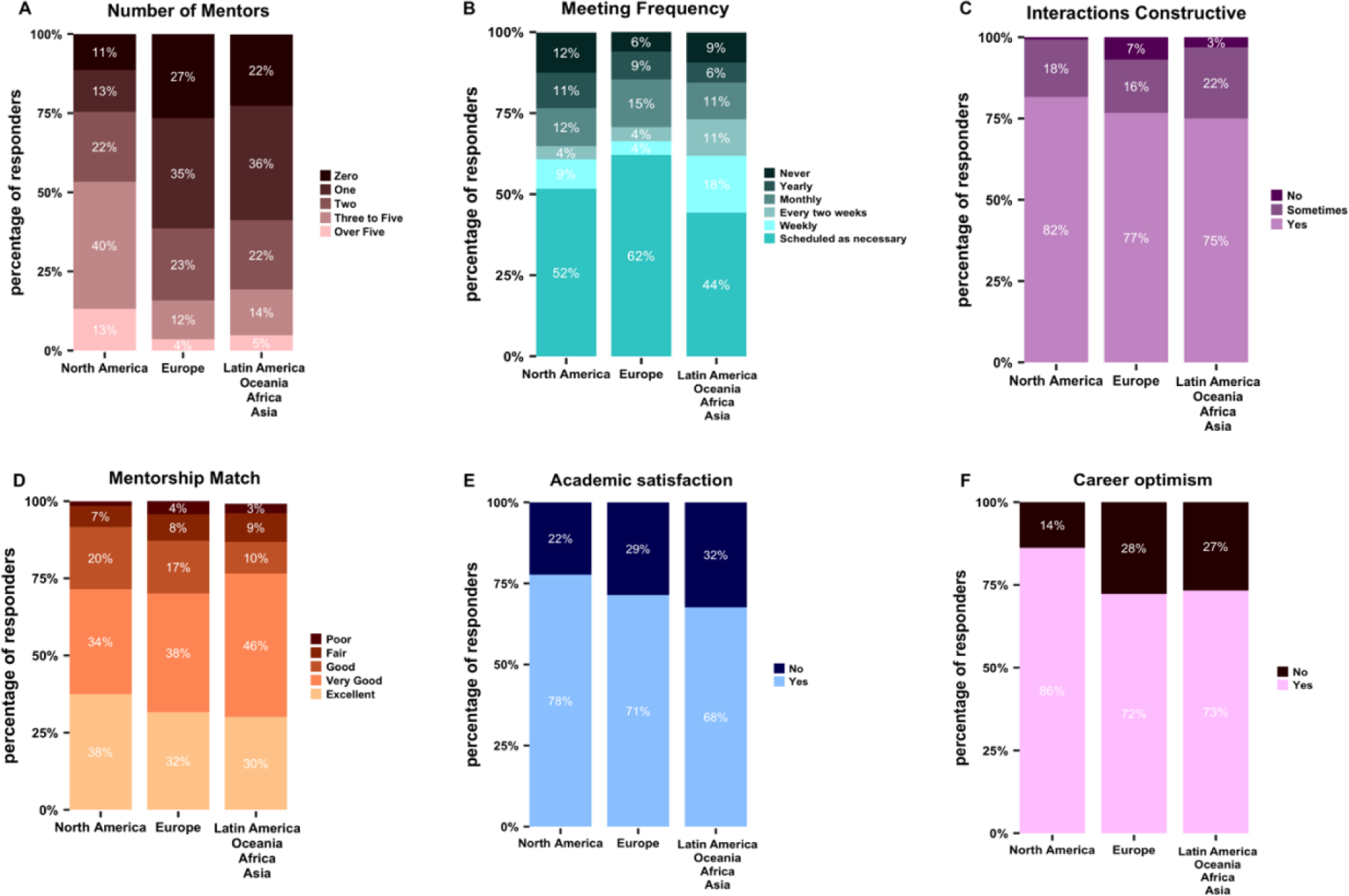
Comparison of mentorship features and quality by geographic region. (A) Number of mentors, North American respondents on average had 3 mentors while respondents from Europe and aggregate of all other continents had one (Table S12a), (B) Frequency of meeting with mentor(s) (Table S7a), (C) Quality of interactions with mentor(s) (Table S10a), (D) Quality of mentorship match (Table S15a), (E) Mentee satisfaction on current research program (Table S22b), and (F) Mentee optimism about their future research and position as an independent investigator (Table S23b) (also see Figure S2e, f). Data analysis for panels B, C, D, E, F excludes survey respondents with "0" number of mentors.

**Figure 7.**
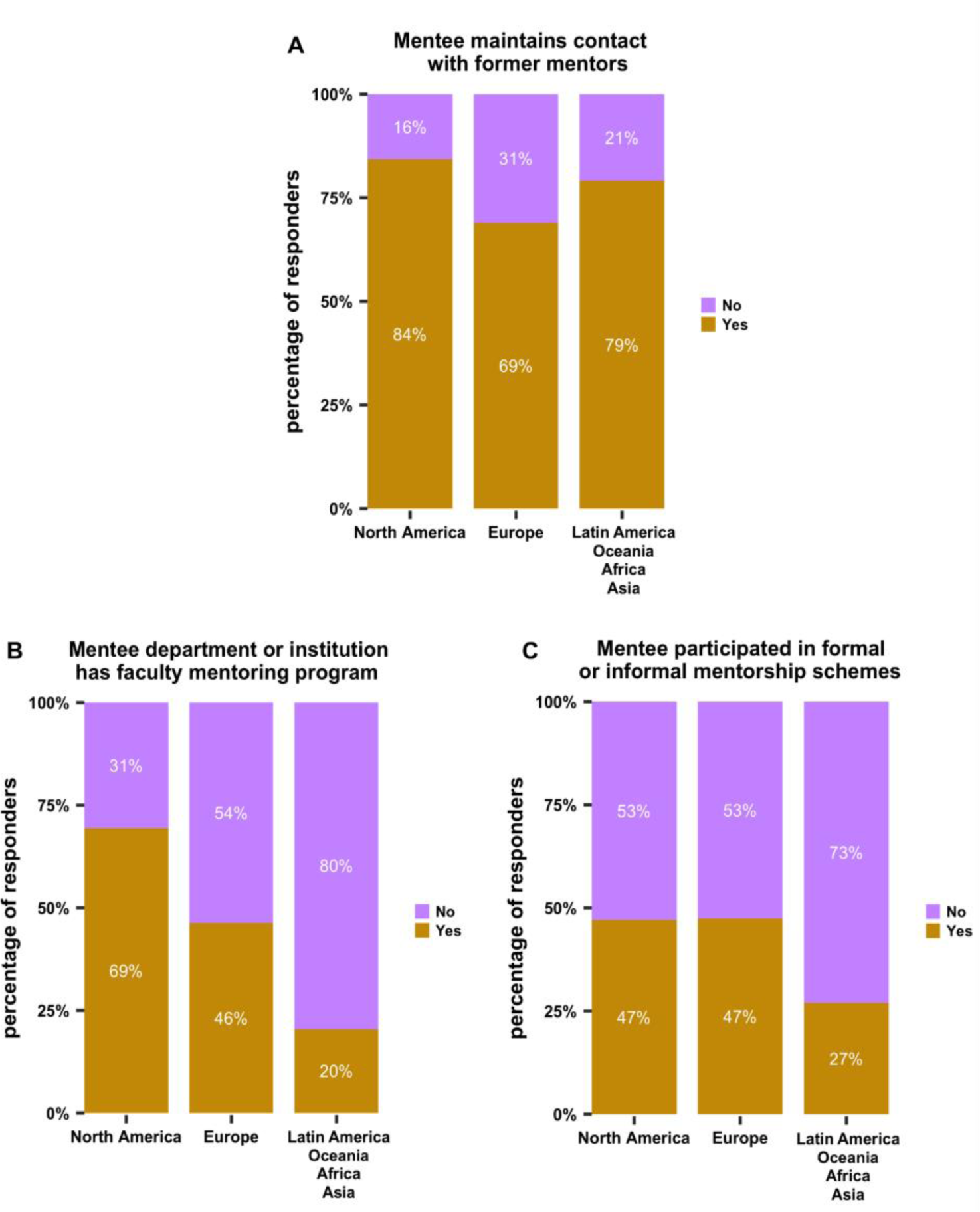
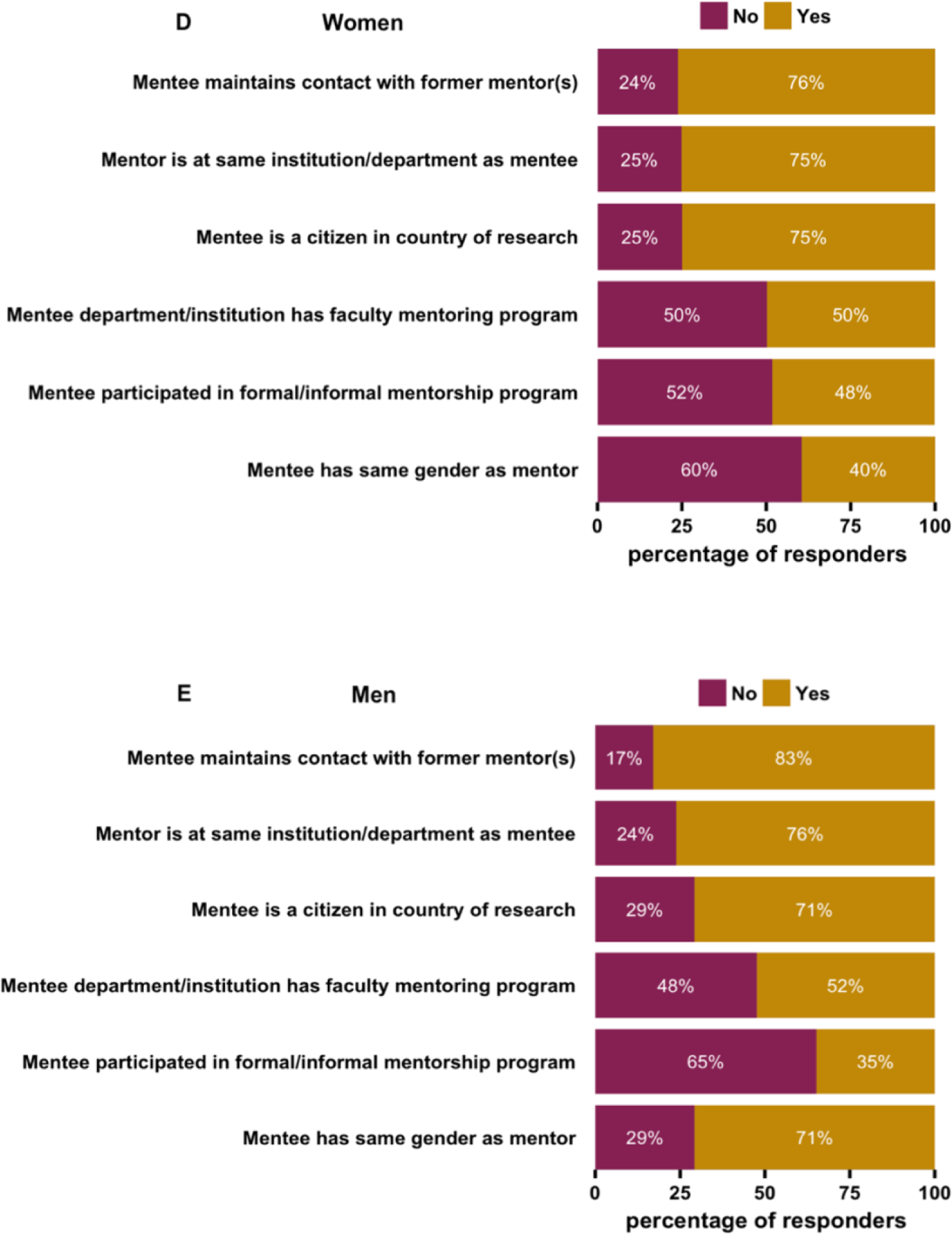
Mentee and mentorship characteristics by geographic region and gender. Whether faculty mentee (A) maintained contact with former mentor(s), (B) institution hired as an independent investigator provided a faculty mentoring program to mentor the faculty mentee on their own career, (C) participated in any formal (i.e., run by institutions) or informal programs (Tables S16a,19a,20a). (D-E) Analysis by gender shown for Mentor location (same department or institution or not), mentor accessibility, mentor gender (same as mentee or not), whether faculty mentee maintained contact with former mentor(s), whether faculty mentee participated in any formal (run by institutions) or informal programs, whether the institution respondents were hired at as an independent investigator provides a faculty mentoring program to mentor mentee on their own career, faculty mentee citizenship status in the country of research (Tables S16-23).

Half of respondents did not have a faculty mentoring program in their department or institution, and most reported that they had not participated in such a program (52% of women and 65% of men) (Figure 7D-E). Despite this, 75% of women and 76% of men reported having at least one mentor within their department or institution, while 25% reported having additional mentors at other institutions (Figure 7D-E).

We asked respondents to rate the characteristics of their mentor and the extent to which their mentor met their expectations. This included questions on how their mentor interacted with them (e.g., used active listening, was trustworthy, provided constructive feedback), whether their mentor advocated for them (e.g., acknowledged professional contributions, helped in acquiring important resources, and helped strategize and prioritize goals), whether the mentor ensured a hospitable working environment (e.g., decreased or eliminated workplace discrimination and harassment, valued and promoted diverse backgrounds, and acknowledged potential biases and prejudices). The most pressing issues that mentors could improve upon were advising on work-life balance (negatively perceived by 36% of women and 29% men), networking and introducing mentee to colleagues (negatively perceived by 33% of women and 18% of men), guidance on mentoring lab members (negatively perceived by 29% of women and 20% of men), sourcing grants or other resources (negatively perceived by 18% of women and 18% of men), and strategizing mentee career goals (negatively perceived by 17% of women and 13% of men) (Figure 8). However, these issues were perceived similarly across genders, with no significant differences across these comparisons (Figure S5).

**Figure 8.**
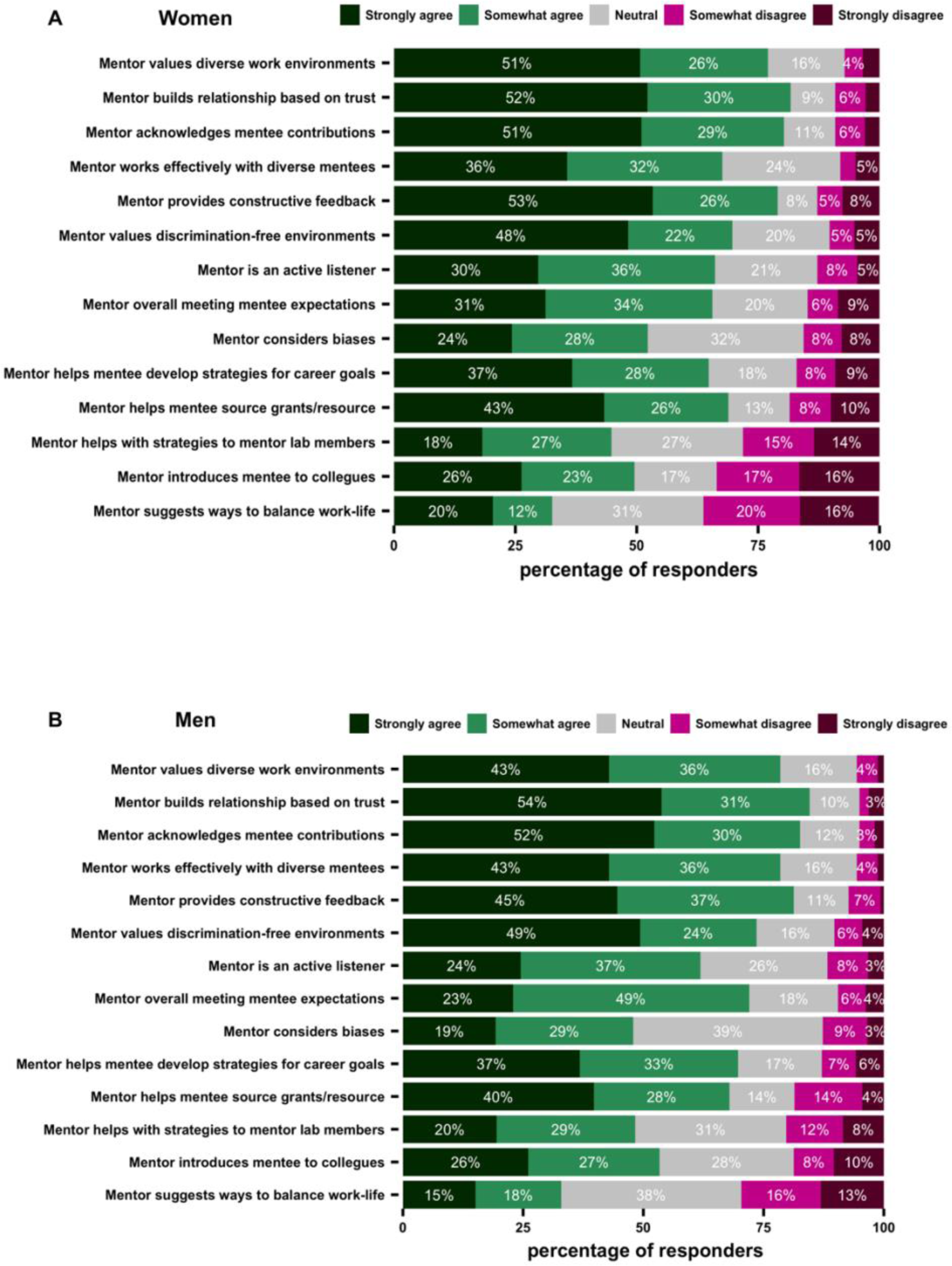
Characterization of mentor-mentee interactions by gender. Respondents evaluated their mentorship interactions on a Likert scale (Tables S24-S37).

### Faculty mentorship across continents

In analyzing responses on mentorship, trends suggest differences in number of mentors, mentee valuation of mentorship interactions based on mentee geographic location. We compared North America, Europe, and "all other continents combined" as this resulted in reasonably sized subdivisions of our data. Faculty in Europe and other continents had fewer mentors (in Europe 27% had zero mentors, 73% had one or more mentors) compared to faculty North America (11% had zero mentors, 89% had one or more mentors) (Figure 6A, S7a, p <0.001). Further, fewer European PIs (39%) had multiple mentors compared to North American PIs (75%). Respondents across continents met with similar frequency frequently (weekly, every two weeks, monthly or yearly) with their mentors (32%) compared to North American faculty (36%) (Figure 6B, S7b, p=n.s.). Respondents across continents found their mentorship interactions constructive to similar levels (Figure 6C, S7c, p=n.s.). Both European and North American faculty expressed similar and good-to-excellent mentorship matches (87% versus 92%) (Figure 6D, S7d, p=n.s.). Interestingly, there was no significant difference in mentorship quality reported for mentees with more mentors across geographic regions (Figure S3c, p=n.s.). These data reveal that North American faculty report significantly more mentors than faculty in other continents (Figure S3d), but this does not translate into significant differences in mentorship quality (Figure S3e). Thirty one percent of European group leaders did not maintain contact with former mentors compared to 16% of North American faculty with only significant differences between North America and European faculty mentees (Figure 7A, S6a, p <0.001). Significantly more departmental or institutional faculty mentoring programs were reported to have been provided to North American group leaders relative to faculty in Europe or other continents (Figure 7B, S6b, p <0.001 and p<0.0001). European and North American faculty were also significantly more likely to participate in peer mentorship programs than faculty in other continents combined (Figure 7C, S6c, p <0.01). Only 47% of European and North American faculty and 27% of faculty in other continents combined participated in a formal or informal mentorship program.

### Academic satisfaction and career optimism

The survey further inquired on faculty mentee satisfaction and optimism about their current and future research and position and found that women were less satisfied with their research progress (32% of women versus 19% of men) and less optimistic about the future of their career (27% of women versus 14% of men) (Figure 5E, 5F, S2a, b, S8a, b, p <0.05). Analysis of responses by mentorship initiation format showed mentees who had assigned mentors and mentees who had chosen their mentor voluntarily were similarly satisfied with their research progress (Figure 4D, 4E, S2c, d, S8c, d, p=n.s.). Analyzing responses across continents, Faculty in Europe and other continents expressed similar satisfaction with their research career, but significantly less satisfaction compared to North American faculty (Figure 6E, S2e, S8e, p <0.05). Similarly, North American respondents and expressed higher career optimism compared to group leaders from Europe or all other continents (Figure 6F, S2f, S8f, p<0.001). While presence or absence of mentors may be a significant contributing factor to respondents’ academic satisfaction and career optimism, differences in funding and career stability, biases, and other structural issues are additional likely contributing factors.

### Analysis of qualitative responses

We asked respondents, in the form of long response questions, if their interactions with mentors were constructive, and if not, what they were seeking in the mentorship relationship. Using these data, we summarized key features displayed by mentors that were noted as helpful (Figure 9A) or unhelpful (Figure 10A). Many of the issues raised by mentees appear to be caused by poor alignment of expectations, including mentees’ want for emotional support, professional development, career guidance and sponsorship. Through a compilation of responses from mentees on their mentoring relationships, we have identified common pitfalls in mentorship as noted by faculty mentees.

**Figure 9.**
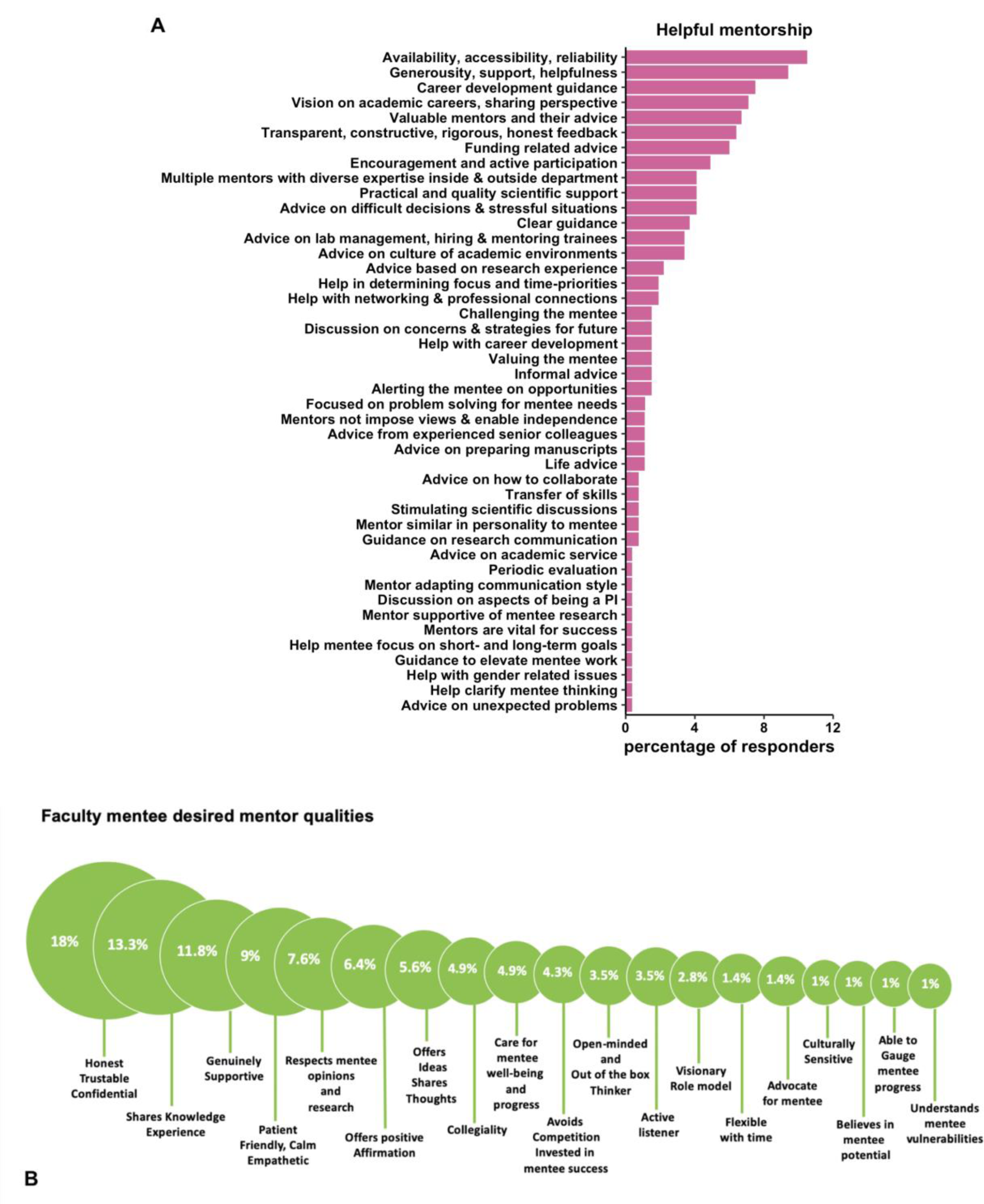
Percent of faculty mentee responses on various aspects of their mentorship aggregated as mentorship themes. (A) What mentees found helpful about their mentoring experience (Total of n=267 comments were received). (B) What qualities they found essential for mentors (Total of n=144 comments were received) (Table S46-S47).

**Figure 10.**
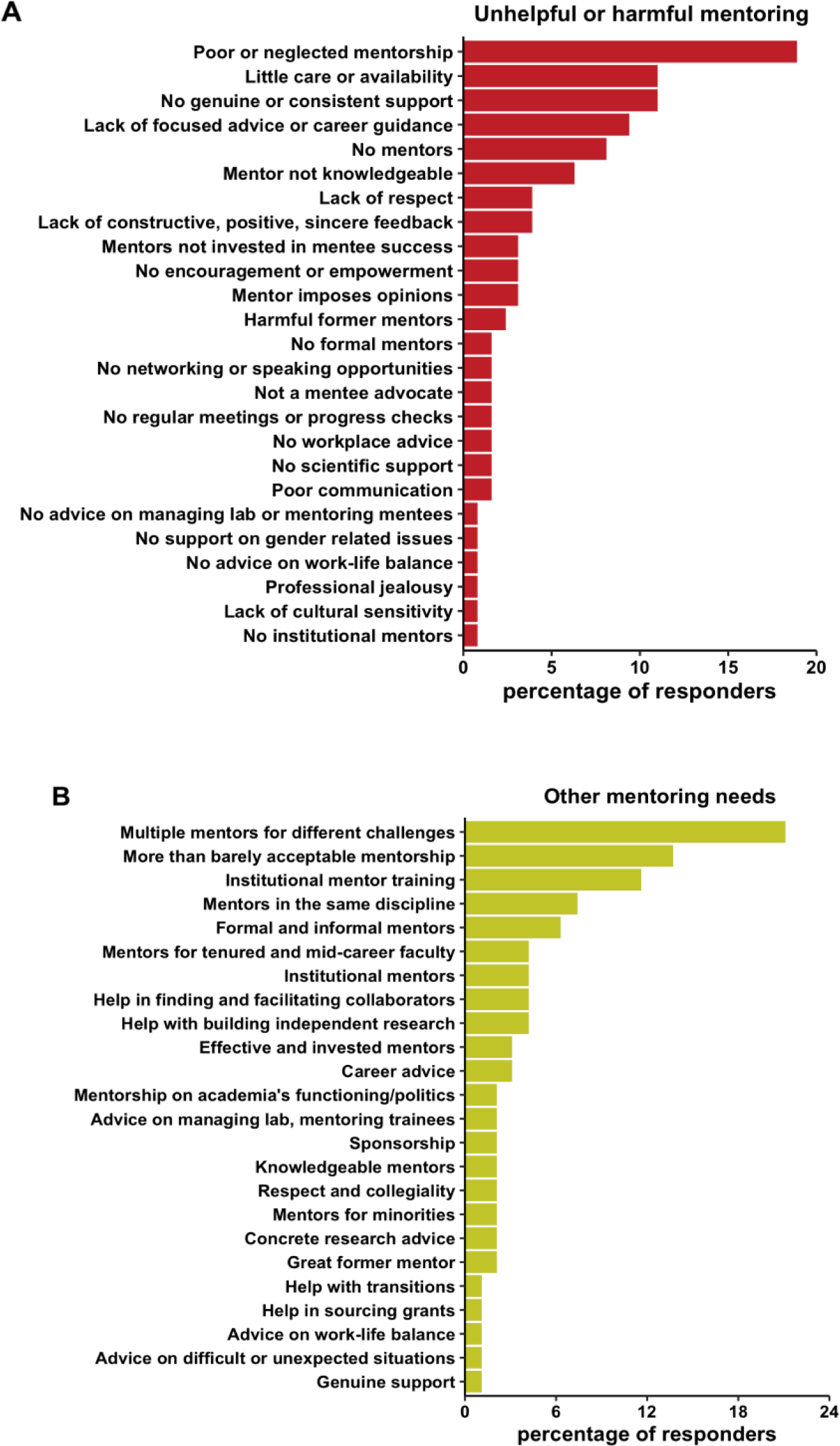
Percent mentee PI responses on various aspects of their mentorship aggregated as mentorship themes (A) Which aspects of mentoring were unhelpful or harmful (Total of n=127 comments were received) (B) What else faculty mentee felt was needed in mentorship (Total of n=95 comments were received) (S48-S49).

### Mentorship is of variable accessibility and quality to faculty

Survey responses indicated a wide range of access to mentorship for faculty. Concerningly, 20% of all respondents did not receive any substantive mentoring upon their transition to an independent investigator position (Figure 3B), but the underlying causes fueling this lack of mentorship varied. Some faculty reported organizational failures in promoting mentorship due to a lack of formal mentoring programs or an absence of mentoring culture. However, even access to institutional mentoring programs was not sufficient to promote proper mentoring if poorly designed or implemented. Some respondents noted that although they were assigned an individual mentor, that this mentor did not make an effort to understand what type of support the mentee needed, leaving the mentee on their own. Collectively, the data indicate that most faculty desire effective mentorship from more experienced faculty, and suggest that in many contexts, current mentorship plans and programs could be improved to, at a minimum, provide consistent guidelines to enable all early and mid-career faculty to access at least one experienced mentor.

Those receiving mentorship reported a large range in quality of interactions with their mentor(s). In agreement with quantitative survey responses (Figures 5-6), in some cases, this was driven by geographical or cultural differences, as respondents noted that in some countries it is uncommon for faculty to source mentors at their university, whereas in others it is normal or mandatory. This variability in quality was also driven by the mechanisms in which mentoring relationships are established. Some respondents indicated that their mentor was assigned but failed to provide substantial value to their careers, indicating a mismatch in the mentoring relationship, personality-wise or in terms of specific scientific interests, leading to some respondents noting that their mentor did not appreciate or value their research. A scientific mismatch led to mentees feeling that their mentors lacked important insights on research approaches, or were unable to appreciate and vet ideas regarding their research program. Other respondents noted that their mentor offered useful guidelines for research excellence and how to obtain research funds, but failed to ensure a working environment free of harassment and bias. Generational differences also impacted perspective and could contribute to perceived mismatches in mentorship; some respondents noted that they were unsure whether their senior mentors recognized the differences faced by different generations in academia. Low valuation of mentorship for faculty by senior faculty was also expressed as a common problem, which we elaborate on below.

### Mentorship is not consistently valued amongst faculty mentors

The most common complaints regarding mentoring relationships include poor or neglected mentorship, a lack of care or availability, and a lack of genuine or consistent support. Many mentees noted having mentors who rarely met with them, resulting in an inability to foster a genuine relationship, and a lack of advice on issues consistently and urgently faced by mentees. Others reported that mentors would only make time to meet if asked repeatedly, leaving the mentee feeling as though the relationship was not valued by the mentor. However, mentees recognized that these behaviors may at times be driven by the inherent pressures of academia; many felt that their mentors do care about them, but are under constant pressure to publish or perish, and thus lack the bandwidth to take on the role of an engaged mentor. This disengagement was reflected in feedback that mentors, while helpful, often did not go out of their way to provide guidance, or that when they offered direct assistance, they would fail to follow through with these offers. Still others noted that their faculty mentoring relationship was not inherently poor, but rather turned negative due to neglect or apathy over time.

Another common complaint from mentees was that their mentoring relationships lack perceived value. Many noted that their mentors did not provide focused advice or career guidance and gave little to no constructive feedback to their mentees. For instance, some respondents indicated that it is common knowledge that early career faculty would need high quality papers and grants, finding this level of advice unhelpful. Responders noted that some mentors did not provide effective and constructive advice on how to achieve future goals, nor did they help to develop an actionable career plan. Some respondents noted that mentors often asserted what they would do in a situation, which sometimes was not aligned with mentee interests or made the mentee uncomfortable. Other faculty found that they did not receive specific advice on how to be academically successful in either research or career development, even after soliciting such advice from their mentor.

Such failings in a mentoring relationship may stem from poor communication between mentor and mentee, which was noted by many survey respondents. Some stated that their contact with their mentor was through specific means (e.g., via email) which restricted their interactions. Others found mentors were difficult to talk to, had poor listening skills, or projected their own experience upon mentees. However, poor communication can also originate from the mentee, as multiple respondents noted that they were not completely forthcoming about their struggles, leading to misunderstandings about their needs.

A number of responses mentioned specific, harmful, and concerning behaviors of mentors. Respondents noted mentors that were negative and often ’forget’ to include or support mentee at key moments. Other respondents noted mentors who never read any material the mentee requested feedback on. Respondents noted mentors who cheered on mentees, and provided positive feedback, yet had a hierarchical attitude and belittled mentees by calling them inexperienced. Some noted poor or unrealistic advising, including recommendations that mentees abandon all projects not destined for top journals, which mentees found discouraging. Some respondents noted hostile untrustworthy mentors that undermined junior faculty. A number of mentees experienced bullying, or felt that their mentor exploited their funding and junior position for their own gains. For instance, mentors took advantage of the forced relationship for mentees to conduct experiments for their own work. Some mentors triggered conflicts between the mentee’s group and their own group, causing unnecessary conflicts within their department. Other junior faculty noted what they perceived to be professional jealousy, such that some mentors acted as if they saw mentee as a competitor for departmental resources. These behaviors are particularly concerning, as they document instances in which the mentorship of junior faculty is exploitative and would have a highly stressful and negative impact on their and their trainee’s careers.

### Recommendations for improving mentorship for faculty

Through a compilation of responses from mentees with both positive and negative mentoring relationships, we have assembled the following set of optimal characteristics desired by faculty mentees to be captured by faculty mentors, departmental leadership and institutions in faculty mentorship.

### Mentors can help improve and optimize mentorship for faculty mentees

#### Need for honesty, active listening, generosity, vision

One powerful outcome of our survey analysis is the emergence of a common set of qualities desired in a faculty mentor, which include honesty, trustworthiness, and an ability to have confidential discussions. The latter attribute was strongly valued (Figure 9B), and respondents mentioned the need for cordial and productive interactions with their mentors. Respondents valued mentors who made interactions with mentees relaxed, which allowed mentees the ability to speak frankly to a colleague and have transparent conversations about the highs and lows of the academic workplace. Mentees also value the opportunity to discuss topics without judgment from their mentor, and to receive unvarnished, truthful advice from an experienced perspective. Respondents desired their mentors’ honest opinions about their progress and future career, and highly valued sincerity. Some noted that mentors could be vague or afraid to direct the mentee in a specific way, but respondents appreciated mentors who were not afraid to tell the mentee when something could be improved.

#### Need for supportive and knowledgeable mentors across topics

Many respondents suggested that the best mentors show genuine support and concern for the careers and general well-being of their mentees (Figure 9A). Respondents appreciated when their mentor adapted to their unique personality and view of science, providing personalized genuineness to the relationship. Respondents desired mentors who offer encouragement, positive affirmation, and who hold a strong belief in mentee potential. Interestingly, while these qualities were consistently valued amongst respondents, these desires manifested in different objectives in mentoring relationships; while some respondents wanted mentors who value mentee qualities and contributions in publications (i.e., scientific contributions), other respondents noted that they value mentors on a personal level who made them feel safe and professionally valued. These differences in opinions captured in our qualitative questions highlight the various roles that mentors can have, both scientifically and personally. For example, some respondents noted being assigned a specific mentor (e.g., through a professional society) who was a fantastic mentor in professional situations, but could not offer support on other aspects of being a faculty. This suggests that mentees would be best served by identifying multiple mentors that can each provide mentorship in distinct roles including setting up a lab, the navigation of academia, and ways to manage work-life balance.

#### Need for advice setting up and running a lab

Mentees expressed an appreciation for mentors who supported their ideas and offered advice on career development. Respondents valued mentors who offered clear guidance and targeted advice when asked as well as general perspectives on common challenges. Effective mentors were reported as inquisitive, asking thought-provoking questions to both brainstorm ideas and provide focused advice. This advice was particularly useful when helping the mentee focus on their short- and long-term goals. Respondents appreciated informal advice that helped in clarifying the mentee’s thinking. Respondents noted mentors who were interested in helping mentees overcome faculty career transition difficulties such as helping mentees choose the best collaborations, hire suitable students, protect mentee time to focus on their research, and help mentees identify their own projects to build independent research. Mentees noted that effective mentors neither judge nor impose their own ideas or views, but rather allow the mentee to explore all possibilities before chiming in with their own opinion. These behaviors allow mentors to promote personal growth and encourage new ideas.

Respondents also wished for structured support in developing their research profile as faculty. Mentees further desired guidance on improving research communication, on preparing manuscripts and job applications, hiring and promotion advice, and advising on service efforts. Respondents mentioned that when they had mentors, they appreciated mentors’ advice on which conferences to attend, and identifying journals to which they could submit their work. Respondents desired mentorship and advice on seeking funding, advice on how various funding agencies function, how to write grants, and how to address grant revisions. Some respondents noted that their mentor had been particularly helpful in reviewing their grants. Respondents wished for a major effort from mentors to exploit their academic potential, to facilitate, establish and sustain collaborations, to produce better scientific projects.

Many respondents of our survey noted that there is no real training on how to manage a lab and little to no support available when things go wrong. Respondents wished for a mentor to provide advice and tips on how to recruit personnel, manage staff, and mentor their own trainees. Mentees also value advice on how to prioritize their time while minimizing energy spent on irrelevant matters (both academic and work relationships). Being offered these perspectives based on experiences otherwise unavailable to mentees proves highly valuable. Respondents noted that the transition to independence is difficult, and that having supportive mentors eases these difficulties substantially.

#### Need for mentors to assist with navigating academia across scales

Amongst the most commonly expressed needs of junior faculty are to have access to senior faculty to help them navigate academia as a system. This need typically will span multiple scales, with mentees needing assistance in understanding the policies and politics of their department, their institution, and their scientific field. Respondents noted that a mentor who could speak to all the necessary components to be successful in their department would be highly beneficial, suggesting junior faculty should seek out at least one senior member within their department from which they can obtain this information. Ideally, this departmental mentor could also speak in an unbiased fashion about departmental politics and relevant scenarios that may impact the professional life of the mentee.

Respondents also sought useful discussion on the tenure process, mentee career progression, and establishing themselves in their scientific fields. Mentees report valuing transparent, constructive, and rigorous feedback to ensure they are on track for a successful tenure case. Respondents wished that their mentor provided appropriate support and guidance to build up their career, including unbiased advice on career moves, career options or lack thereof, should these issues become relevant during the course of the mentee’s career. Mentees note that they value mentors who can anticipate areas that the mentee needs to think about that the mentee may not be aware of (i.e., who can flag "unknown unknowns"). Mentees desired strategic tips regarding mid-career planning spanning grant application strategies to deciding how to expand or focus the research interests of the mentee’s lab. Thus, mentees find value in a mentorship relationship well into their established careers, as the challenges faced by faculty are constantly changing throughout their careers.

#### Need for mentors’ sponsorship and advocacy

Respondents report that the best mentors also serve as their advocates in looking after mentee interests. This includes actively supporting their career development through discussions about research and career strategy while also recognizing mentee contributions to research. Amongst the most impactful actions that a mentor can have is to help mentees identify opportunities for their own work and to promote the mentee’s career. This could include introducing the mentee to senior research leaders in their discipline or promoting mentee successes to senior institutional leadership and to the wider community. Other respondents commented on their appreciation for mentors who tried to remove any barriers to mentee progress, whether at the administrative, scientific, or professional relationship level. Respondents hoped for mentors who understand their privilege or how to navigate in a changing academic culture.

#### Need for personalized mentorship for faculty

Beyond scientific and professional advice, respondents desired well-rounded mentors, noting the importance of having someone who celebrates and encourages success and growth above and beyond publishing papers and who can offer wisdom on issues such as work-life balance. Respondents noted that having mentors to advise on unique, stressful, or unexpected situations, to help the mentee navigate through challenges was valuable in the transition to independence. Mentees noted that having mentors reminds them that they have a support system when they feel lost on the academic journey. This can be particularly valuable for women, underrepresented minorities, or immigrants, who would highly benefit from mentorship on specific additional challenges that each of these groups face on both a scientific and personal level. Notably, these comments highlight the heterogeneity in faculty members and thus their mentorship needs, highlighting the need for diverse mentors to help mentees on their own unique route to success.

#### Need for mentor initiative, investment and commitment

Respondents noted that mentors need to be invested in and have care for mentee success, spending time and effort to improve the mentoring relationship. Mentees particularly appreciate mentors who are generous with their time (Figure 9B), noting that as a mentee it was reassuring to know that they could ask questions of their mentor without feeling that they were bothersome. Respondents noted mentors exceeding expectations with their help, at times even outside of typical work hours, and many acknowledged and appreciated their mentors taking time out of their busy schedule with no formal recognition or little benefit to them. Mentees particularly valued interacting with mentors with sufficient time to provide contextual advice, rather than standard aphorisms that apply to all. Many mentees desired a mentor who puts mentee career development and progress ahead of their personal interests and genuinely cared for the mentee’s progress and success.

#### Need for accessible and reliable mentors

Mentees highly preferred mentors who were accessible, available, and reliable. Many noted that their mentor met when necessary (as also evident in quantitative responses Figure 5B), and, as a result, was readily accessible to provide practical advice. An open-door policy was noted as critical, as some mentees noted that they felt as though they could only approach their mentor with important or pressing matters, or when they had specific questions to be answered. Some respondents noted that at times it was difficult to access their mentor but they had a positive relationship with their mentor, and when they did meet, the meetings were excellent. In these cases, the initiative to continue the mentoring relationship came almost exclusively from the mentee. Thus, the best mentors not only invest time to advise their mentees, but also take initiative in checking in on them.

Some noted the mentor being excellent and compassionate, dedicating significant time to the relationship, seeing the whole picture on how to get to the final goal (whether a grant or being published in appropriate journals), but still had a poor mentoring relationship with the mentee. This suggests that even ‘effective’ mentors can be mismatched with mentees and may lack key skills for cultivating a healthy mentorship relationship, further highlighting the need for mentor training programs.

Some respondents noted their mentor made good suggestions, but were generally hands off, which encouraged mentee independence. This raises an interesting dichotomy, as some mentees crave a hands-off relationship to promote independence, while others perceive this as a lack of interest. Thus, open communication is critical for a solid mentorship relationship to allow dynamic interactions that can ebb and flow as the relationship progresses and as mentorship needs change. These differences in expectations also highlight the need for personalized, tailored mentorship to promote the success of junior faculty.

### Departments, Institutions and funders can improve mentorship for faculty

#### Need for formal Institutional and departmental mentors

Our survey revealed that mentors both inside and outside of a mentee’s institution were important to provide guidance on the tenure process and mentee career progression. Respondents noted that formal mentorship schemes were important, as a lack of formal mentorship typically translated to few specific expectations from their mentors. In the absence of such schemes, ongoing support from a more senior colleague was deemed valuable by respondents, suggesting that informal mentorship is valued by junior faculty. Consistently, some survey respondents noted that no formal mentoring scheme exists in their institution because mentees felt they worked in a collegial environment with a low tenure bar where informal mentoring was readily available.

Respondents who only reported mentors at other institutions desired more frequent interactions with mentors, noting that at times it was difficult to get advice or feel supported. Some respondents also noted having an assigned mentor at their host institution, but that this mentor was not helpful or trustworthy. Thus, without institutional structure and active assistance from the departmental level, some faculty may not obtain necessary support to find a mentor. Respondents noted that working as a junior faculty member is difficult without a formal mentor, and, in the absence of effective mentorship, sought alternative sources for guidance. For instance, some respondents noted their postdoctoral advisor had acted as their unofficial mentor after transitioning to their faculty position, helping much more than any mentor at their current institution. Respondents also noted that having formally assigned mentors was desirable because they felt more freedom to contact their mentor, feeling less as though they were being bothersome because of the formality of the established relationship. Some also hoped that multiple formal interactions would lead to an informal relationship.

#### Need for informal mentorship

Responders noted having mentors who were not assigned to them but had mentored them of their own accord, naturally establishing informal mentoring relationships. Some responders noted that faculty mentorship should be informal and diverse, with many different faculty colleagues, including mentors in mentee’s specific discipline and on an as-needed basis. Some noted that their institution had a formal mentor program that was underwhelming, necessitating mentorship to be sourced elsewhere. Some respondents noted preferring a mentor of the same gender and outside their line management hierarchy. Others noted having a formal mentorship relationship that only made sure the mentee checked boxes for tenure. Respondents also noted that had they not sought mentors on their own, they would not have been as happy nor as effective in their job. Thus, informal mentors could serve as valuable additions to a mentorship team, each bringing different assets to mentee life and career.

#### Institutions need to offer mentorship and mentor training programs

Some respondents noted that their university did not have formal mentoring programs, effectively setting a low bar as to what to expect from a mentor. Mentoring programs for early career faculty on the tenure track have recently been introduced at a number of institutions in the United States (“Columbia University Guide to Best Practices in Faculty Mentoring: A Roadmap for Departments, Schools, Mentors and Mentees,” n.d.; “University of Michigan-Dearborn Faculty Mentoring Program,” n.d.; Feldman, 2017). Responders to our survey noted that they would have appreciated having a mentor on-site to talk to often in person. Responders noted that not many faculty are skilled in mentoring, with training and buy-in being key for both mentors and mentees. Some respondents who had informal mentors noted that their great mentors had received training on mentoring. Some also noted that their institution did not have a mentor program, so they felt fortunate to have a mentor, believing it should be an essential part of every academic institution. Some respondents noted that their institution was too small to have anyone who works in their specific field, suggesting that inter-institutional mentorship programs would be valuable. Some respondents noted a complete lack of mentors for their research and for receiving tenure, resulting in their desire to change the system to be more supportive of the current junior faculty. Responders also believed that it would be impossible for academics to be expected to be good mentors without training. Therefore, provided the number of skills required to be developed to become a good mentor, each faculty mentee likely needs more than one mentor assigned with a clear expectation. This would be far more easily achieved through formal mentoring schemes, which would be enabled by institutional support for mentorship training programs.

#### Need for multiple and diverse mentors or a mentorship team

Our survey suggests that highly effective mentorship comes from teams consisting of a diverse set of individuals and experiences as opposed to traditional mentoring dyads. Of respondents who found the mentorship they received barely acceptable, many noted that the advice they received had been valuable in only context-dependent situations, but was lacking in others. These respondents commented that their main mentors were valuable, but noted that there were some important professional aspects for which this one mentor was not well-suited, suggesting a mentorship team would be more helpful. This was true for other sets of mentees as well: some noted wished for a committee of mentors that were closer to their field as a strategy to mitigate the lack of knowledge or expertise from one specific mentor. Some answered the survey questions in regards to a variety of mentors (authors had asked respondents to respond with regards to their key mentor), noting that they received excellent mentoring from multiple mentors and that they had a great relationship with their primary mentor. Some responders also chose to focus on one of their closest institutional mentors in response to our survey, thinking that their shortcomings were more important to highlight. Still others noted that no one person was perfect in any specific thing and that different perspectives by multiple mentors are always illuminating, showing that there isn’t a single right way to attempt addressing various academic work and life challenges.

The benefits of mentoring teams were supported by comments from respondents’ desired perspective from a third party who can help the mentee evaluate their standing and track record. Sometimes one has to figure out what a person is experienced at to understand who would be the best person to ask for advice on certain issues. Respondents noted having chosen a group of mentors, each for specific skills. Each mentor did a great job at their task, which was effective as the mentee did not expect one mentor to advise on all topics. Other responders noted having an equal number of men and women mentors, locally, from previous institutions and also other institutions within the geographic area. Mentees approached mentors selectively, depending on the question or problem they were facing. Other responders noted having two senior males and a senior female mentor who provide complementary information about how to advance a mentee’s career. Some respondents noted having several mentors from multiple programs or departments.

#### Need for peer mentorship communities

Some respondents pointed out that as female academics, lack of mentorship and lack of academic support from senior researchers was a major contributor to the leaky pipeline and women’s opportunities for the transition from postdoc to independent investigator. Some reported that other than brief interactions with mentors (2-3 people, less than one hour per year for specific questions), that their only mentors are online resources (e.g., Twitter or online communities such as the New PI Slack and/or the mid-career PI Slack), noting that they received much more help and support to grow academically from peer mentorship on these forums. While peer support is helpful, it is not sufficient and not the same as institutional or formal mentorship from more senior faculty. This finding indicates the role of technological advances in shaping mentoring relationships, and further research is required to understand the relative benefits of social media to conventional in-person contacts.

#### Maintaining valuable interactions with former mentors

Some respondents noted having maintained mentorship interactions with former mentors (∼25% of respondents in quantitative questions did not, Figure 7D-E). Some noted that they had a supportive graduate advisor and, while their formal obligations to each other ended with the mentee’s graduation, mentor support and mentorship did not end. Some noted that despite their current mentorship situation being unhelpful, they still valued and received mentorship via email from their doctoral supervisor, some going on for decades. Reciprocally, some respondents noted having far more positive and pleasant interactions with their current faculty mentors despite unpleasant mentoring interactions during their training. Responders noted they valued mentors who prioritized mentee well-being over productivity. Responders also noted that they learned how to choose a mentor based on prior extremely negative mentor experiences. Some respondents mentioned mentors they had since graduate training that had guided the mentee through many aspects of their career development and job transitions.

#### Mid-career faculty also need mentors

The qualitative responses highlighted mid-career faculty as an overlooked community in mentoring relationships. Some tenured or established faculty noted that their mentor was their department chair, who was not a mentor in an official capacity, rather the only person they received substantial advice from since arrival. Some noted that in their institution, junior faculty had mentorship committees but that this was not the case for tenured faculty who joined the department. Some respondents noted that they mentor graduate students, postdoctoral researchers and tenure-track faculty, informally and as part of a mentoring program, but that they had never themselves been offered any formal mentoring from their institution. Other mid-career faculty noted that they did not have mentors themselves and now had unrealistic expectations placed on themselves by their students. So, in order to alleviate student stress and protect their work life balance and mental health, mid-career faculty assume that they have to provide their mentees with support all by themselves. Some mid-career faculty respondents noted receiving mentorship on-site and off-campus. Some noted benefiting from supportive professors externally who had been encouraging, but internally it seemed to be expected that faculty wouldn’t benefit from mentorship once they were promoted to a tenured or associate professor position, even though their funding opportunities are reduced compared with junior colleagues. Many regard mentorship for early (pre-tenure) faculty as much more available than mentorship for mid-career faculty. It is harder to find peer mentors within the institution to help guide the jump from associate to full professorship, and there are fewer online resources and tutorials on this transition. Thus, there is a need to facilitate continued mentorship for faculty members, even upon the transition to tenure.

## Discussion

Our goal in conducting this study was to document the attributes and showcase the impact of faculty mentorship. We believe that illuminating the current state of faculty mentorship will draw attention to areas that require improvements to elevate faculty mentoring worldwide. We analyzed a survey of 457 early and mid-career faculty from over 40 countries to characterize features of mentorship interactions. The findings of this survey reveal interesting insights about the current state of faculty mentorship. First, we found that up to a quarter of faculty across continents did not have mentors, and that many (16-31%) did not maintain contact with former (i.e., graduate or postdoctoral) mentors. This suggests that a substantial portion of early and mid-career faculty do not have adequate access to mentorship - even from past mentors - and the benefits that it would provide. Based on our survey results, the most likely contributor to differential access to mentorship is faculty mentee geographic location. Specifically, twice as many faculty located in Latin America, Oceania, Africa, Asia, or Europe reported having zero mentors as their colleagues in North America (Figure 6). Furthermore, we found that North American faculty on average have more mentors (an average of 3 mentors) compared to Europeans and all other continents aggregated (which have an average of 1 mentor). Finally, departments and institutions in North America have more faculty mentoring programs and faculty themselves seek more mentors (Figure 7B-C). This suggests cultural differences in seeking and providing mentorship, and likely points to important considerations in both the perceptions and effective implementation of mentorship strategies worldwide.

Beyond geographic differences, we found differences in how men and women faculty perceive mentorship. Female faculty indicated less satisfaction regarding the mentorship that they received (Figure 5) and lower optimism on future career prospects than their male counterparts (Figure 5). This is consistent with differences previously observed in women faculties’ perception of their work environment from their male counterparts (Levinson et al., 1989) and additional obstacles women face in moving up the tenure ladder, in part due to work-family balance (Ward and Sloane, 2003; Webber and Rogers, 2018) and the academic workplace climate (Spoon et al., 2023).

Despite some faculty receiving suboptimal mentorship, many respondents of our survey not only received mentorship, but also reported finding their interactions with their mentor(s) constructive (Figure 2D) and their mentorship match of high quality (Figure 3E). Survey respondents emphasized that having a diverse set of mentors that span a range of areas of expertise worked well to accommodate their needs. This often included multiple mentors - both formal and informal - to provide many potential sources for advice on different aspects of a scientist’s job, as consistent with previous studies (DeCastro et al., 2013). For instance, junior faculty would almost certainly benefit from a mentor within their department to assist in navigating institutional guidelines and politics, but would additionally benefit from a senior mentor within their field to identify opportunities for networking, exposure, and promoting their research program. Additionally, junior faculty would also likely benefit from a cohort of informal peer mentors that share resources and offer mutual learning to assist with questions on laboratory management, grant writing, and managing work-life balance. It would be highly improbable to find all of these qualities in just one person, highlighting the strengths of having a mentorship team to navigate being an academic faculty member. Indeed, our respondents noted that mentors are vital for success, and a number of respondents mentioned that without support from mentors they would have left academia.

While many faculty were satisfied with their faculty mentorship, our survey indicates that there is room for improvement across mentoring relationships. The most prominent issues in need of attention involve adequately addressing work-life balance, networking, training in mentorship and grant writing, and helping the mentee to strategize career goals. There is also room to improve accessibility to mentorship, as we found a large range of variability in the institutional commitment to mentees and mentors alike. Our survey found that as many as 40-80% of their institutions had not implemented formal mentorship programs, and that over 50% of faculty were not participating in any formal or informal mentorship schemes. Given that there is demand for mentors in the same specialty, and for at least some mentors outside of the institutional hierarchy, scientific societies could organize mentoring between universities. A number of prominent issues with mentorship were identified in our survey which could be further addressed by departmental and university leadership reevaluating their mentoring programs. While mentorship for faculty and faculty mentorship programs may be new concepts in many institutions worldwide, our survey respondents valued mentorship as one of the most important factors in long-term success and faculty retention (Figure 9A,10B). Thus, investing in the short-term efforts of ensuring proper mentorship of junior faculty will likely result in substantial long-term gains for departments and institutions.

There are specific actions that can be taken at the departmental and institutional levels to effectively promote faculty mentorship. First, it is important for departments and institutions to provide quality mentoring relationships, as described by our qualitative survey responses, as opposed to merely assigning faculty mentor(s) in the department. As there will be variations in mentee needs and mentor skills and knowledge, care and consideration should be taken to match junior faculty with senior faculty that can promote a positive mentoring experience. This should include consulting junior faculty on their needs and would benefit from soliciting their opinion (or, minimally, their approval) of proposed mentors. Departments should promote continuous and dynamic conversations on mentorship among colleagues and university administration, rather than promoting a “one size fits all” mentorship solution (Figure 11). Along with discussing the merits of faculty membership, departments and institutions should provide protected time and training for mentors and mentees (Greco et al., 2022; Jackevicius et al., 2014; Lumpkin, 2011). Protected time to ensure quality mentorship will reinforce its value, and departments could further incentivize quality mentorship by including mentorship metrics as permanent components of promotion, hiring, and funding decisions. Departmental and institutional leadership could also promote more effective mentoring interactions by including this evaluation metric in tenure and promotion rubrics (Anna Hatch et al., 2019; Hatch and Curry, 2020; Hicks et al., 2015; Moher et al., 2018).

**Figure 11.**
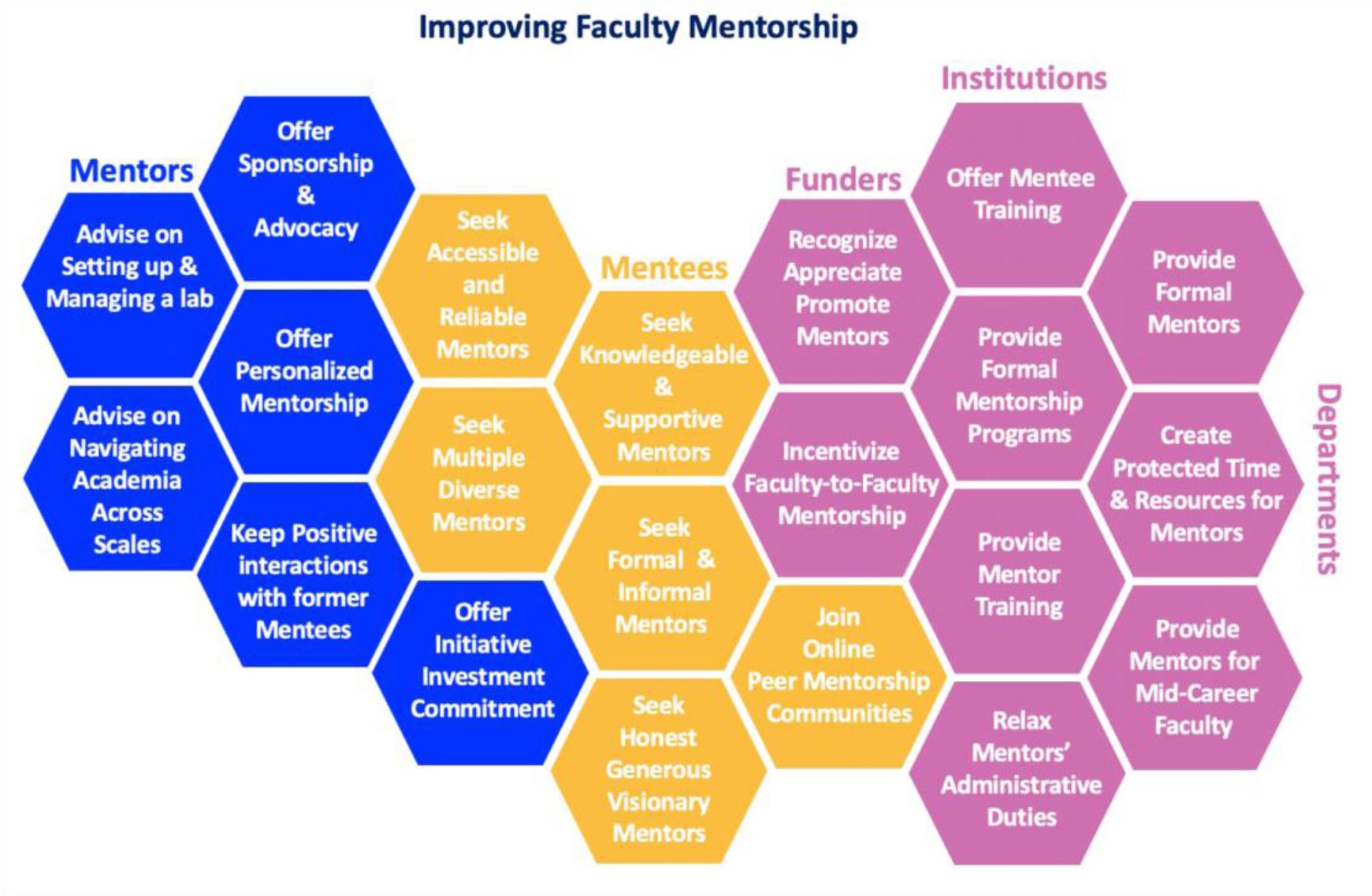
Summary of faculty mentorship recommendations for faculty mentees, mentors, funders, departments and institutions.

Alongside departmental and institutional mechanisms, improving faculty mentorship should also include listening to early and mid-career colleagues and appreciating their specific needs. Mentors can go out of their way to promote their new colleagues by suggesting talented students apply to their mentees’ labs, or to foster their network of colleagues. By listening to the mentee’s needs instead of enforcing the mentor’s own ideas for their development, they will instill a culture of effective mentorship that will hopefully be perpetuated when newer faculty ultimately become mentors creating a sustainably supportive academic environment.

### Potential issues faced by faculty mentors

Our survey focused on the evaluation of faculty mentorship from the viewpoints of junior faculty mentees, without consideration of the perceptions of their senior colleagues who take on the role of faculty mentors. The challenges reported in our survey may have been a key reason driving some faculty mentees to seek and create their own networks for mentorship (i.e., early-career and mid-career PI Slack groups). The root causes underlying suboptimal mentorship from senior faculty may vary. The results seem to indicate a need for mentorship training for senior faculty mentors, as our findings may reflect that faculty mentors don’t always understand mentee needs. It is unclear what faculty mentor competencies are and how these are determined, and whether senior faculty mentors are consistently aware of mentee needs and challenges in early- and mid-career stages. Additional contributors likely include the lack of protected time for faculty mentors, minimal institutional support to provide bandwidth for faculty to be mentors, and little relief from administrative and service duties to make time for mentorship. Furthermore, mentorship is typically neither recognized nor rewarded, and many faculty who spend less time on service activities (including mentorship) advance to full professorship more quickly than those who spend more time on service. Burnout could also be a contributing factor to lack of or suboptimal faculty-to-faculty mentorship (even for faculty who want to be mentors) as there may just not be enough time to mentor well. Once faculty are tenured, they are expected to increase teaching, research, service, and administrative duties, rendering the very faculty most qualified to be mentors the least available to perform this critical task. Thus, it is crucial that higher education and funding institutions openly allocate time and resources to promote mentorship so that implementing recommendations for best mentoring practices become feasible and not a burden to the overwhelmed faculty. Given the importance of mentorship to early- and mid-career faculty in increasing their performance, satisfaction, and retention, departments and institutions would strongly benefit from incentivizing mentorship programs and relationships for senior faculty members.

### Strengths and limitations

In this work, we reported the findings of a large and independent survey of faculty members who may represent as many as 457 academic departments worldwide. While some departments and institutions may run faculty mentorship surveys internally, their findings are often not made public, suggesting a need for data curation on the broad needs of faculty mentorship. As our survey was completed before the COVID-19 pandemic, mentoring perceptions may have changed in some aspects that are not reflected in our survey. The COVID-19 pandemic also may have reduced mentoring, at least for those who were funded to travel to conferences and informally seek external mentorship. Our responses are highly enriched in faculty in the life and biomedical sciences and medical research currently residing in North America or Europe; the survey was presented in English, thus influencing our reach. Nations with English-speaking researchers, on social media and English-speaking universities, have the highest number of researchers per million inhabitants, so it is common for similar research culture surveys to have responses concentrated in North America and Europe. Despite this commonality, this concentration of responses influences the inferences we can make about mentorship relationships worldwide, particularly in geographic regions not well represented in our survey responses. We did not collect data on the race or ethnicity of respondents and therefore cannot know how this may have influenced the findings of our survey. We also did not ask for specific information about the types of institutions that were surveyed, suggesting there may be undocumented differences between mentorship relationships at primarily research institutions (i.e., R1 institutions) versus primarily undergraduate institutions (i.e., R2 institutions). As the funding and tenure criteria are different at R1 and R2 institutions, this almost certainly affects mentorship needs of faculty, and future surveys could explore this aspect. Future surveys could also focus on faculty mentors, their mentorship training and their challenges and triumphs in the faculty-to-faculty mentoring interactions. Future research could also study how the mentorship challenges identified in this study may impact faculty mentee productivity, tenure and promotion outcomes and the mentorship of their mentees.

## Methods

### Survey Participant Recruitment

The text for the survey used in this work is included in the supplementary information. A Google form was used to conduct the survey and was distributed on various social media platforms including Twitter and academic Slack groups and via group leaders/faculty/principal investigators worldwide. The survey was distributed from March 2019 to March 2020 and contained both scaled-response and open-ended questions. Responses to the first survey question with "0" for number of mentors, were routed out of the data analysis for certain analysis as specified in the supplementary tables supporting the figures. The respondents to the survey were asked to self-report, and the information collected was not independently verified.

## Data Analysis

Microsoft Excel, Rstudio, ggplot package (Wickham, 2016) and the eulerr package (Larsson and Gustafsson, 2018) were used to graph the results shown in Figures 1-10 and Figures S1-S2. The qualitative survey comments were categorized by theme (keywords and context) describing each comment and the frequency of comments pertaining to a particular theme are summarized in Results and Discussion. Barplots were generated from distinct themes in qualitative responses (Figures 9 and 10). Figure 11 was created using Microsoft Powerpoint. For statistical analyses, Prism 9 was used to perform Ordinary one-way Analysis of variance (ANOVA) (Figure S3, S8), two-tailed Mann-Whitney U (Wilcoxon Rank Sum) test (Figure S4, S5) and the Turkey multiple comparisons test (Figure S6, S7, S8).

## Data Availability

The authors confirm that, for approved reasons, access restrictions apply to the data underlying the findings. Raw data underlying this study cannot be made publicly available in order to safeguard participant anonymity and that of their organizations. Ethical approval for the project was granted on the basis that only aggregated data is provided (as has been provided in the supplementary tables) (with appropriate anonymization) as part of this publication.

## Conflicts of Interest

The authors declare no competing interests.

## Statement of Ethics

The survey has been verified by the Johns Hopkins University Institutional Review Board (IRB) as Exempt, project number IRB-00012432.

## Author contributions

SS and SJB designed and distributed the survey. SS, SJB and NMN performed the data analysis. SS performed the data tabulation and visualization. SS, SJB, NMN wrote the original draft of the manuscript. SS, SJB, NMN, CTS, AWBF, AI, KC wrote, edited and reviewed the manuscript.

## Supporting information

Supplementary Information_Faculty-to-Faculty_Mentorship_Sarabipour et al 2023

## Acknowledgements

The authors thank all the faculty members worldwide who took the time to respond to the survey and offer their valuable input in support of this work. The author also thank Feilim Mac Gabhann, Richard Sever, Paul Macklin, Christian Frezza, Bradley Wyble, Roy Ziegelstein, Daniel Goodman, Iain Cheeseman, Megan Peters, Gunnar Blohm, Anne Urai, Stephen Royle, Marc Robinson-Rechavi and Sophien Kamoun for their valuable comments on an earlier version of this manuscript.

## Notes

### Competing Interest Statement

The authors have declared no competing interest.

### Summary of Updates

The new version of our paper includes additional statistical analysis, figures and tables on the quantitative survey data.

