## Supplementary Information_Faculty-to-Faculty_Mentorship_Sarabipour et al 2023 for "The faculty-to-faculty mentorship experience: a survey on challenges and recommendations for improvements"

#### Survey Questions

***SECTION 1: Mentoring Skills of your mentor (or mentors)***

1) How many Mentors do you currently have (i.e., senior faculty at your department/program or other senior or peer professors that provide mentorship)? (if the answer is zero please continue to the end). Mark only one oval.
0
1
2
3
4
5
>5

2) What do you like about your mentoring experiences?
Long Answer:---------------------

***SECTION 2: Mentor background:*** If you have multiple mentors please focus on the one mentor that covers your most important requirements.

3) Is your Mentor located in the same department/Institution as you? Mark only one option.
Yes
No

4) How long have you been working with your Mentor? Mark only one option.
Less than one year
1-2 years
3-4 years
5 years or more

5) How were you originally matched to this Mentor? Mark only one option.
Voluntarily (I chose the Mentor)
I was assigned to my Mentor.

6) Are you the same gender as your Mentor? Mark only one option.
Yes
No

7) Is your Mentor physically accessible? same campus, same building (so that you have unplanned meetings or encounters)? Mark only one option.
Yes
No

8) How frequently do you have formal meetings with your Mentor? Mark only one option.
weekly
every 2 weeks
monthly
Yearly
scheduled as necessary
Never

9) What format do you use to communicate with your Mentor? Tick all options that apply.
Face-to-Face meeting
Telephone
Email
Web/Video conferencing (e.g., Skype, Zoom)
Social networking sites
Other: Short Answer:---------------------

10) Please rate the quality of the match between you and your Mentor. Mark only one option.
Excellent
Very Good
Good
Fair
Poor

11) Do you find your interactions with your Mentor constructive? Mark only one option.
Yes
No
Sometimes

12) If you answered “NO” or "Sometimes", What type of mentoring relationship were you looking for? What were you hoping to achieve with your Mentor?
Long Answer:---------------------

13) Do you maintain contact with your former Mentors? Mark only one option.
Yes
No

14) Have you ever participated in any formal (run by institutions) or informal peer-mentorship Programs? Mark only one option.
Yes
No

15) Does the institution you are hired at as an independent investigator provide a faculty mentoring program to mentor you on your own career? Mark only one option.
Yes
No

***Your Mentor's Skills***
Please indicate whether you agree with the following statements about your Mentor. We understand that you can only speak from your personal experience. Please try to rate a skill whenever possible, reserving the 'not observed' category for cases where you have no basis for assessment. If a particular statement is not relevant to your mentoring relationship please skip.

16) Uses active listening (Meaning Identifying and accommodating different communication styles and employing strategies to improve communication with you). Mark only one option.

Strongly Disagree Neutral Strongly Agree

1 2 3 4 5

17) Established a relationship based on trust with you

Strongly Disagree Neutral Strongly Agree

1 2 3 4 5

18) Provides you with constructive feedback & positive affirmation

Strongly Disagree Neutral Strongly Agree

1 2 3 4 5

19) Ensures a working environment free from discrimination and harassment

Strongly Disagree Neutral Strongly Agree

1 2 3 4 5

20) Values a working environment/colleagues with a diverse background?

Strongly Disagree Neutral Strongly Agree

1 2 3 4 5

21) Takes into account the biases and prejudices s/he brings to your mentor/mentee relationship

Strongly Disagree Neutral Strongly Agree

1 2 3 4 5

22) Helps you acquire resources (e.g., grants, instrumentation, collaborations etc.)

Strongly Disagree Neutral Strongly Agree

1 2 3 4 5

23) Acknowledges your professional contributions

Strongly Disagree Neutral Strongly Agree

1 2 3 4 5

24) Helps you develop strategies to meet career goals

Strongly Disagree Neutral Strongly Agree

1 2 3 4 5

25) Suggests ways to you to balance work with your personal life

Strongly Disagree Neutral Strongly Agree

1 2 3 4 5

26) Introduce you to speakers and other Professors at meetings and seminars and scientific Society gatherings?

Strongly Disagree Neutral Strongly Agree

1 2 3 4 5

27) Works effectively with you whose personal background is different from his/her own (age, race, gender, class, region, culture, religion, family composition etc.)

Strongly Disagree Neutral Strongly Agree

1 2 3 4 5

28) Helps you develop strategies to better mentor your own mentees (undergraduate/graduate/postdoctoral trainees)?

Strongly Disagree Neutral Strongly Agree

1 2 3 4 5

29) Overall, to what extent do you feel that your current mentor is meeting your expectations?

Strongly Disagree Neutral Strongly Agree

1 2 3 4 5

30) Please explain your answer to the previous question: Long Answer--------------------

31) Do you feel happy & satisfied about your current research and position as an independent investigator? Mark only one option.
Yes
No

32) Please rate your feeling of happiness and satisfaction with your current research and position as an independent investigator?

very Disappointed Neutral very Optimistic

1 2 3 4 5

33) Do you feel optimistic about your future career? Mark only one option.
Yes
No

34) Please rate your optimism about your future career

very Disappointed Neutral very Optimistic

1 2 3 4 5

***Section 3: About You*** (the early to mid-career faculty Mentee): Professional Background

35) Which category below describes you best? Please choose only one oval.
Tenure-track Assistant professor

Tenured Associate professor
Full Professor (Tenured)
Junior Group Leader
Lecturer
Other: Short Answer:---------------------

36) What type of advanced degree do you hold? Mark only one option.
PhD
Professional degree (e.g., MD, DDS, RD, PT, PharmD, etc.)
Both PhD and professional degree (MD/PhD, MD/MPH, PharmD/MS, etc.)
Other: Short Answer:---------------------

37) Which category(s) best describes the focus of your research? Check all options that apply.
Basic(Fundamental) research in Life Sciences
Basic research in Physical Sciences
Engineering
Theoretical/Mathematical research
Translational research (specify type: T1, T2, etc.)
Clinical research
Behavioral research
Field/applied research
Social Science research
Environmental Sciences
Humanities
Medicine
Psychology
Other: Short Answer:---------------------

38) What type of research institution are you based at? Mark only one option.
Academic
Industry
Government
Other: Short Answer:---------------------

***SECTION 4: A bit about your own lab***
39) Does your current lab have a website to introduce your group? Mark only one option.
Yes
No

40) Has your lab posted a welcome letter on your lab website for prospective students? Mark only one option.
Yes
No

41) Have you written a lab manual for current lab members? Mark only one option.
Yes
No

***SECTION 5: Your (Mentee) Demographics***
42) Your Gender: Mark only one option.
Male
Female
Non-Binary
I would like to self-identify:-------------

43) What is your age?--------

44) Are you a citizen in the country you conduct your research at? Mark only one option.
Yes
No

45) Which country do you have your lab and perform research at currently?
Short Answer:-------------------

46) Other Comments about the survey or your experience that was not addressed in the Questions?
Long Answer:----------------------------------------

### Supplementary Figures

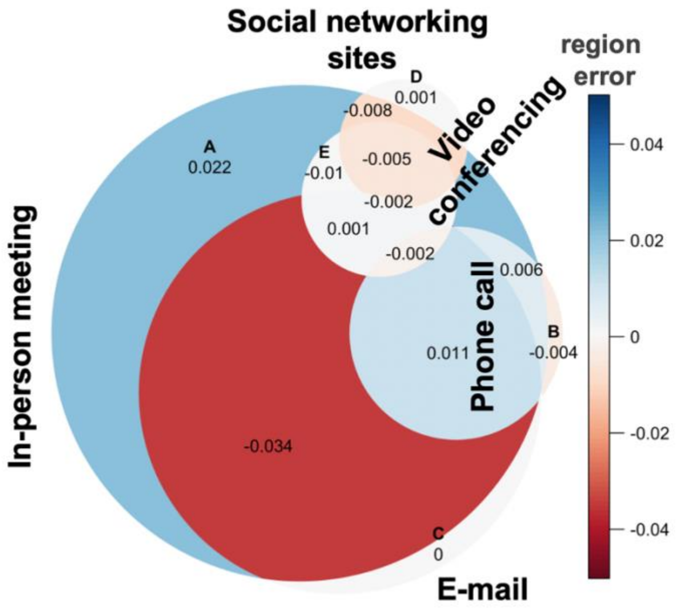

**Figure S1. Mentee demographics.** Error plot for *'euler'* objects, calculated using the eulerr package. The error plot visually evaluates the fit to the call euler() for the optimization performed for the Venn diagram in R studio*. Data analysis excludes survey respondents with "0" number of mentors. All responses were self-identified.

*****Wickham H. 2016 ggplot2: elegant graphics for data analysis.

*Larsson J, Gustafsson P. 2018. A Case Study in Fitting Area-Proportional Euler Diagrams with Ellipses Using eulerr. Proc Int Workshop Set Vis Reason. 2116: 84–91.
https://cran.r-project.org/package=eulerr
https://www.rdocumentation.org/packages/eulerr/versions/7.0.0

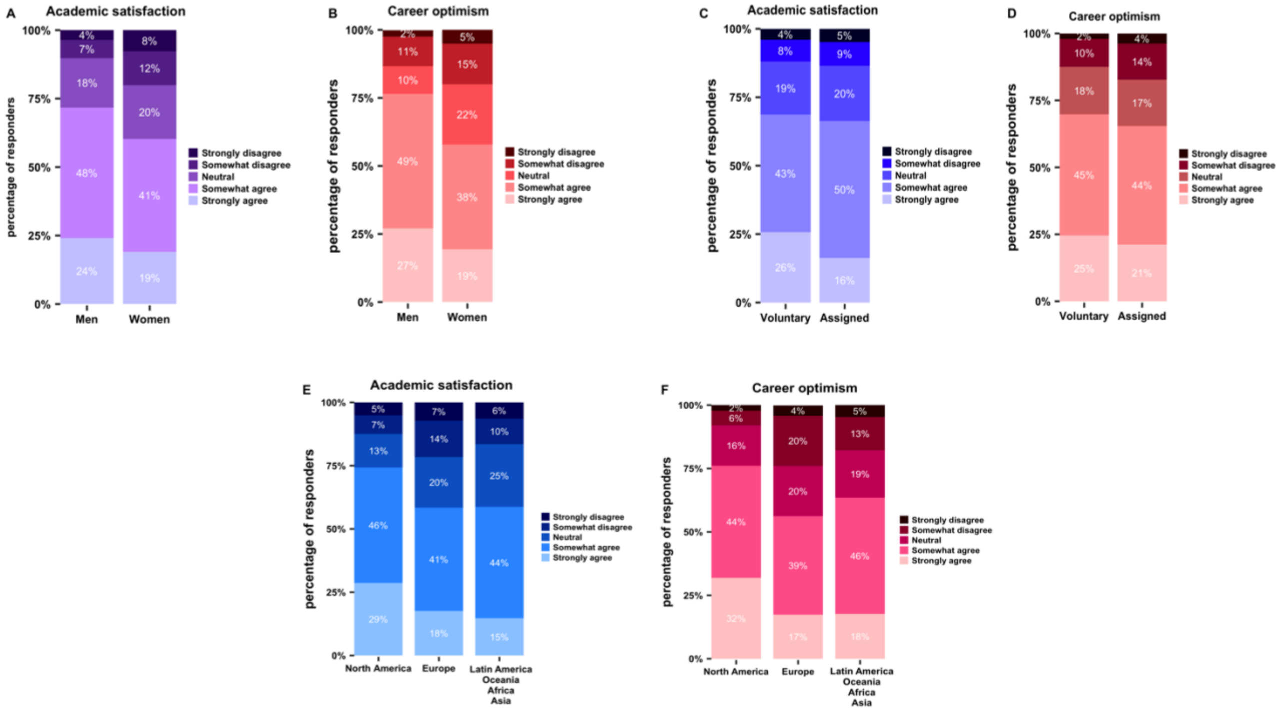

**Figure S2.** Respondents' academic satisfaction and career optimism about their current and future research and position as an independent investigator by (a-b) gender (Tables S38, S39), (c-d) mode of mentorship initiation mode (Table S38b, S39b), and (e-f) respondent geographic region (continent) (Table S38a, S39a). Responses are on Likert scale. Data analysis for all panels in this figure includes survey respondents with "0" number of mentors.

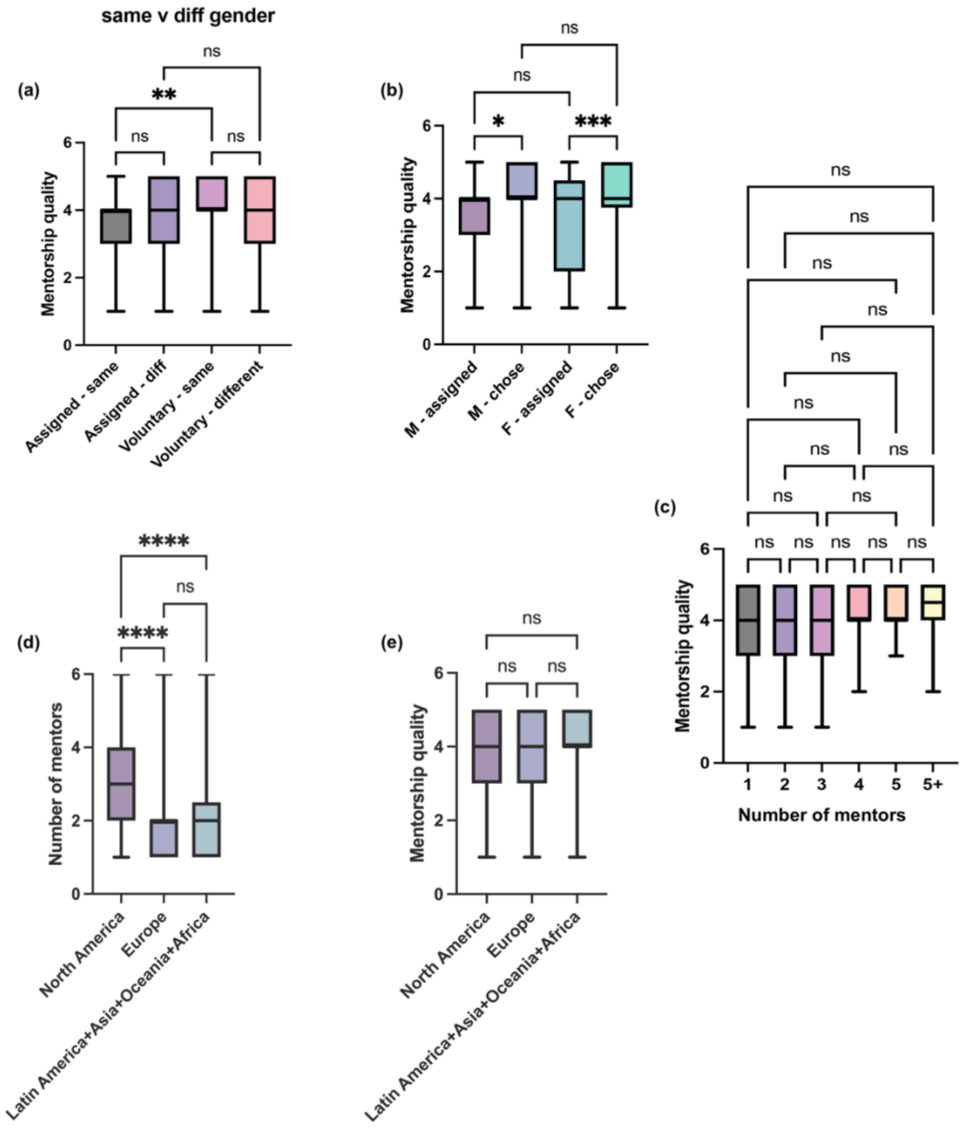

**Figure S3.** Statistical analysis of relationships between mentorship sourcing, gender, number of mentors, and reported quality of mentorship match. (a) Mentorship quality as a proxy of assigned versus voluntary choosing of mentor by a mentee of the same gender or different genders. ** $p<0.01$, n.s. = not significant, Ordinary one-way ANOVA (Table S40a), (b) Mentorship quality as reported by males (M) or females (F) with assigned or chosen mentors. * $p<0.05$, *** $p<0.001$, n.s. = not significant, Ordinary one-way ANOVA (Table S40b), (c) Mentorship quality as reported by number of mentors sourced. n.s. = not significant, Ordinary one-way ANOVA (Table S40c), (d) Number of mentors reported by mentees in North America, Europe, or other continents. **** $p<0.0001$, n.s. = not significant, Ordinary one-way ANOVA (Table S40d), (e) Mentorship quality reported by mentees in North America, Europe, or other continents. n.s. = not significant, Ordinary one-way ANOVA (Table S40e). Data analysis for panels (a), (b), (c), (e) in this figure excludes survey respondents with "0" number of mentors. For all box plots, the box extends from the 25$^{th}$ to 75$^{th}$ percentile of datapoints; whiskers stretch from minimum to maximum datapoints. The line in the middle of the box plot represents the median.

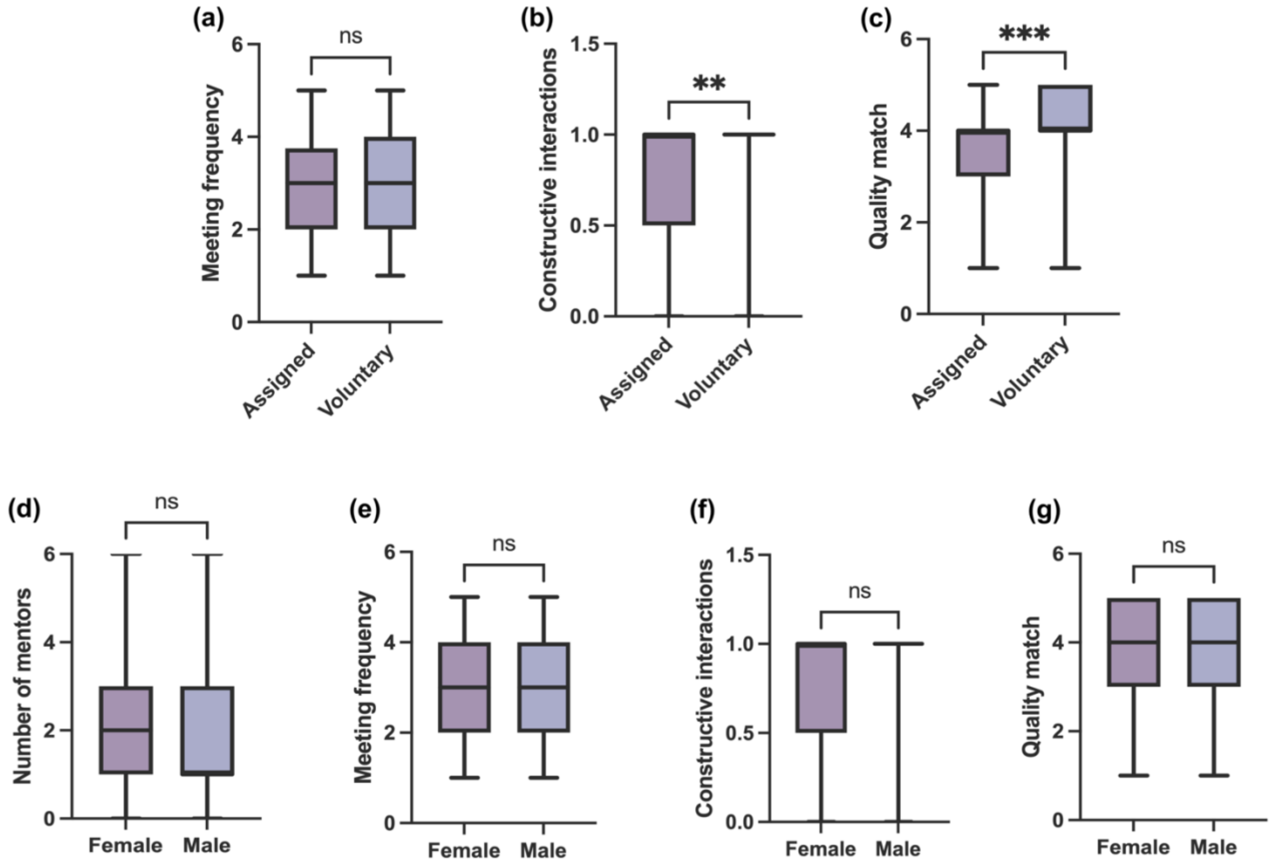

**Figure S4.** Statistical analysis of results shown in Figure 4 and Figure 5. (a) Meeting frequency (Table S41a), (b) constructive mentorship interactions (Table S41b), and (c) perceived quality of mentorship match (Table S41c) based on how mentees sourced their mentors (i.e., voluntarily chose or assigned). n.s. = not significant, ** $p<0.01$, *** $p<0.001$, Wilcoxson Rank Sum Test. (d) Number of mentors (Table S41d), (e) meeting frequency (Table S41e), (f) constructive mentorship interactions (Table S41f), and (g) perceived quality of mentorship match (Table S41g), for men and women mentees. n.s. = not significant, Wilcoxson Rank Sum Test. For all box plots, the box extends from the $25^{th}$ to $75^{th}$ percentile of datapoints; whiskers stretch from minimum to maximum datapoints. The line in the middle of the box plot represents the median.

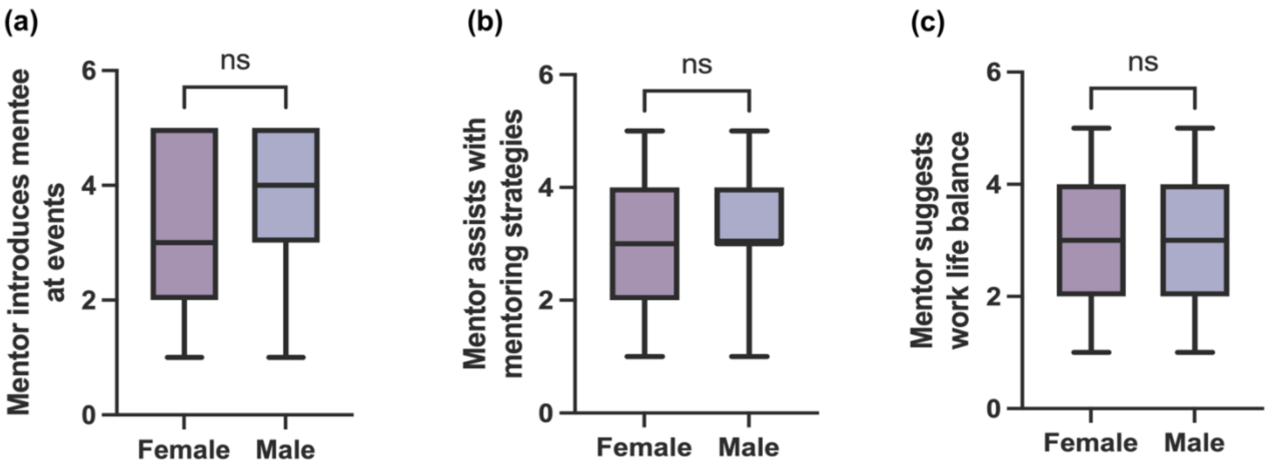

**Figure S5.** Statistical analysis of results shown in Figure 8. (a) Frequency of mentor introducing mentee to network at events (Table S42a), (b) mentor assisting with mentoring strategies for mentee (Table S42b), (c) and mentor suggesting work life balance (Table S42c) for men and women mentees. n.s. = not significant, Wilcoxson Rank Sum Test. For all box plots, the box extends from the 25$^{th}$ to 75$^{th}$ percentile of datapoints; whiskers stretch from minimum to maximum datapoints. The line in the middle of the box plot represents the median.

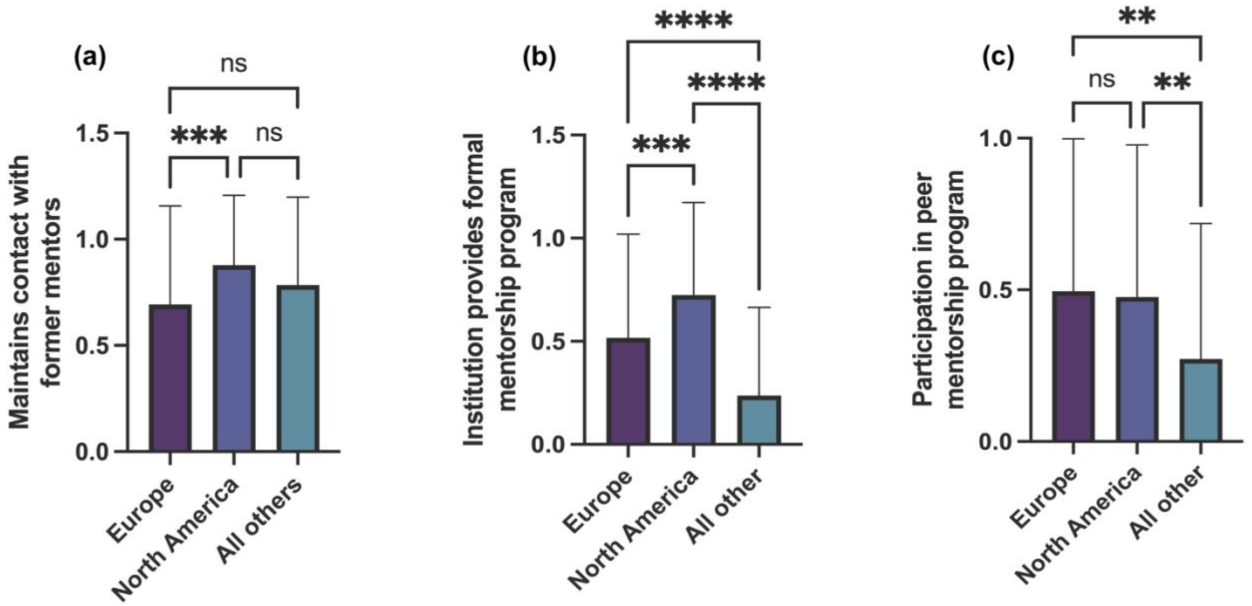

**Figure S6.** Statistical analysis of results shown in Figure 7 A-C. (a) Mentees who maintain contact with former mentors (Table S43a), (b) mentees who report institutional formal mentorship programs (Table S43ab), and (c) mentees who participate in peer mentorship programs (Table S43c) by geographic location. Locations were given as countries, which were sorted into three broad categories based on approximately equal distributions of continental responses, including Europe, North America, and all other continents combined. n.s. = not significant, ** p<0.01, *** p<0.001, **** p<0.0001, Ordinary one-way ANOVA. Error bars on graphs represent standard deviation of responses.

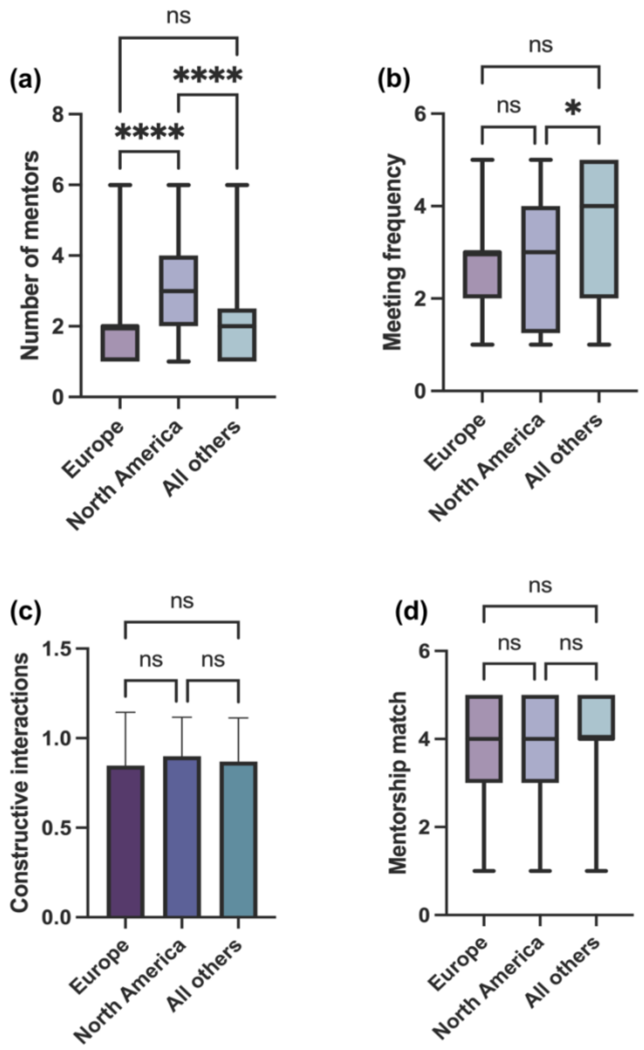

**Figure S7.** Statistical analysis of results shown in Figure 6 A-D. (a) Number of reported mentors (Table S44a), (b) meeting frequency with mentors (Table S44b), (c) constructive mentorship interactions (Table S44c), and (d) perceived mentorship match by mentee geographic location (Table S44d). Locations were given as countries, which were sorted into three broad categories based on approximately equal distributions of continental responses, including Europe, North America, and all other continents combined. n.s. = not significant, * p<0.05, **** p<0.0001, Ordinary one-way ANOVA. Error bars on bar graphs represent standard deviation of responses. For all box plots, the box extends from the 25$^{th}$ to 75$^{th}$ percentile of datapoints; whiskers stretch from minimum to maximum datapoints. The line in the middle of the box plot represents the median.

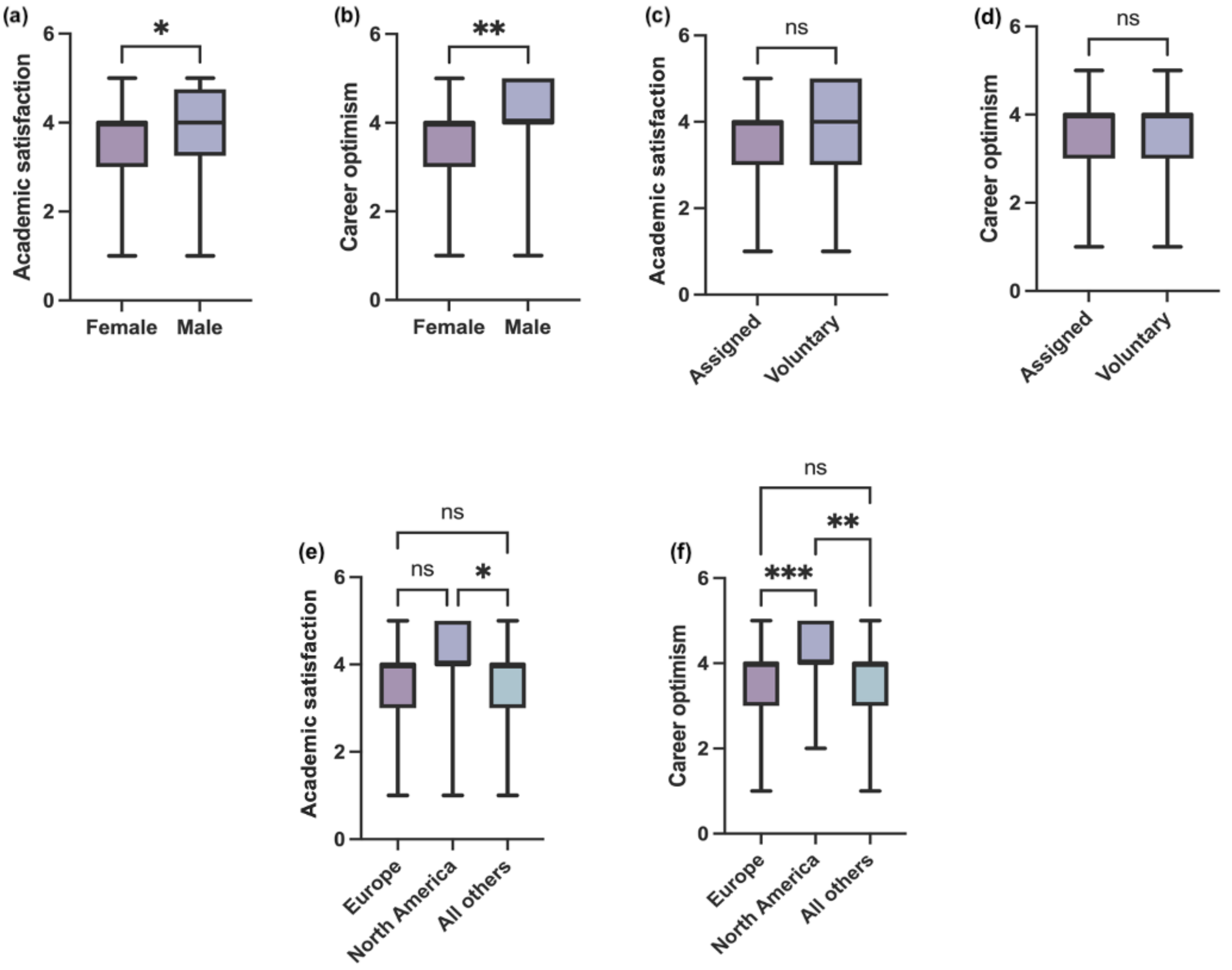

**Figure S8.** Statistical analysis of results shown in Figure S2. (a) Academic satisfaction (Table S45a) and (b) Career optimism (Table S45b) for female and male mentees. (c) Academic satisfaction (Table S45c) and (d) Career optimism (Table S45d) for mentees who were assigned a mentor or voluntarily chose a mentor. Data in (a-d) were analyzed using a Wilcoxon Rank Sum Test; n.s. = not significant, * $p<0.05$, ** $p<0.01$. (e) Academic satisfaction (Table S45e) and (f) Career optimism (Table S45f) based on geographic location. Locations were given as countries, which were sorted into three broad categories based on approximately equal distributions of continental responses, including Europe, North America, and all other continents combined. Data in (e.-f.) were analyzed using an Ordinary one-way ANOVA; n.s. = not significant, * $p<0.05$, ** $p<0.01$, *** $p<0.001$. For all box plots, the box extends from the $25^{th}$ to $75^{th}$ percentile of datapoints; whiskers stretch from minimum to maximum datapoints. The line in the middle of the box plot represents the median.

### Supplementary Tables

**Data Availability.** The authors confirm that, for approved reasons, access restrictions apply to the data underlying the findings. Raw data underlying this study cannot be made publicly available in order to safeguard participant anonymity and that of their organizations. Ethical approval for the project was granted on the basis that only aggregated data is provided (as has been provided in the supplementary tables) (with appropriate anonymization) as part of this publication. There are 49 supplementary Tables referenced in figure captions of the main article and in the supplementary information.

**Table S1. Faculty Demographics: country and continent of research (research group location)**
Distribution of the country of research of faculty/group leaders/principal investigators. Regions which had fewer than four respondents in our survey were aggregated as "Other countries" in the table. All percentages are calculated out of the total number of respondents to this particular survey question (450) not the total number of overall survey respondents (n=457). Data analysis for this table includes survey respondents with "0" number of mentors.

| Respondent Origin (country in which the faculty currently conduct research with their laboratory) | Number of respondents (faculty mentee) |
|---|---|
| **by Continent** | |
| **North America** | 36.7% (165 out of 450) |
| **Europe** | 36.4% (164 out of 450) |
| **Latin America** | 11.8% (53 out of 450) |
| **Asia** | 7.6% (34 out of 450) |
| **Oceania** | 5.3% (24 out of 450) |
| **Africa** | 2.2% (10 out of 450) |
| **by Country** | |
| **United States** | 34.7% (156 out of 450) |
| **United Kingdom** | 16.4% (74 out of 450) |
| **Argentina** | 8.2% (37 out of 450) |
| **Australia** | 5.3% (24 out of 450) |
| **Spain** | 4.7% (21 out of 450) |
| **Germany** | 2.9% (13 out of 450) |
| **Canada** | 2% (9 out of 450) |
| **China** | 2% (8 out of 450) |

| | |
|---|---|
| **Chile** | 2% (8 out of 450) |
| **France** | 2% (9 out of 450) |
| **Malaysia** | 1.3% (6 out of 450) |
| **Sweden** | 1.3% (6 out of 450) |
| **The Netherlands** | 1.1% (5 out of 450) |
| **India** | 1.1% (5 out of 450) |
| **Belgium** | 1.1% (5 out of 450) |
| **Japan** | 1% (4 out of 450) |
| **Denmark** | 1% (4 out of 450) |
| **Croatia** | 1% (4 out of 450) |
| **Panama** | 1% (4 out of 450) |
| **Finland** | 1% (4 out of 450) |
| **Other countries: Austria + Bangladesh + Brazil + Belize (Fiji, Papua New Guinea) + Colombia + Egypt + Ghana + Hungary + Iran + Ireland + Italy + Indonesia + Mali + Mexico + Nigeria + Poland + Portugal + Qatar + South Africa + Norway + Taiwan + Tanzania + Turkey + Thailand + Switzerland + Uruguay + Sierra Leone + Singapore** | 9.5% (43 out of 450) |
| **Did not respond to this survey Question** | 2.22% (7 out of 457) |

| **Table S2. Type of Research Institution faculty mentee was based at (hired at)**<br>Data analysis for this table includes survey respondents with "0" number of mentors. | |
|---|---|
| **Theme** | **Type of research Institution** |
| **Total (n=447)** | 95.3% (426 out of 447) Academic<br>0.45% (2 out of 447) Industry<br>4.3% (19 out of 447) Government |
| **Women Mentee (n=236)** | 94.5% (226 out of 239) Academic<br>0.4% (1 out of 239) Industry<br>5% (12 out of 239) Government |
| **Men Mentee (n=200)** | 95.5% (191 out of 200) Academic<br>0.5% (1 out of 200) Industry<br>4% (8 out of 200) Government |
| **Non-binary or did not disclose gender (n=8)** | 100% (8 out of 8) Academic<br>0% (0 out of 8) Industry<br>0% (0 out of 8) Government |

| | |
|---|---|
| **Did not respond to this survey Question** | 1.8% (8 out of 455) |

**Table S3. Faculty by their Field of Research**
Distribution of survey respondents by self-identified gender and self-identified scientific field.
All percentages are calculated out of the total number of respondents to this question.
Data analysis for this table includes survey respondents with "0" number of mentors.

| Theme | Number of respondents (n=445) |
|---|---|
| **Basic (Fundamental) research in Life and Biomedical Sciences** | 54.3% (242 out of 445) |
| **Basic research in Physical Sciences, Engineering research, Theoretical or Mathematical research, Computing and Information Sciences** | 12.1% (54 out of 445) |
| **Social Sciences, Humanities, Psychology, Behavioral research, Art and Design, Education research** | 9.7% (43 out of 445) |
| **Environmental Sciences, Conservation Biology, Field or applied research, Agriculture, Plant Biology** | 8.8% (39 out of 445) |
| **Clinical Research, Medicine, Public Health** | 7.9% (35 out of 445) |
| **Translational research (type: T1, T2, etc.)** | 7.2% (32 out of 445) |
| **Did not respond to this survey Question** | 2.6% (12 out of 457) |

**Table S4. Respondent Gender**
Respondent self-identified gender, Data analysis for this table includes survey respondents with "0" number of mentors.

| Theme | Number of respondents |
|---|---|
| **Women (n=240)** | 52.5% (240 out of 457) |
| **Men (n=207)** | 45.3% (207 out of 457) |
| **Non-binary/did not disclose gender (n=10)** | 2.2% (10 out of 457) |

| **Table S5. Faculty Mentee Academic Degree**<br>The type of advanced degree faculty held. All percentages are calculated out of the total number of respondents to this question n=446. Data analysis for this table includes survey respondents with "0" number of mentors. | |
|---|---|
| **Theme** | **Number of respondents** |
| **Doctoral Degree (PhD)** | 91.3% (407 out of 446) |
| **Professional degree (e.g., MD, DDS, RD, PT, PharmD, etc.)** | 2% (9 out of 446) |
| **Both PhD and professional degree (MD/PhD, MD/MPH, PharmD/MS, etc.)** | 6.3% (28 out of 446) |
| **Master's Degree (Science, Arts)** | 0.5% (2 out of 446) |
| **Did not respond to this survey Question** | 2.4% (11 out of 457) |

| **Table S6. Faculty Mentee Academic Position Type**<br>Faculty self-reported academic positions at their institution. All percentages are calculated out of the total number of respondents to this question. Assistant Professor (tenure-track and non-tenure track) and Associate Professorship (tenure-track and non-tenure track) positions are held in North America (United States of America and Canada). Lecturer, senior lecturer and research fellow positions are mainly held at institutions in the United Kingdom. Group leader faculty positions are held throughout Europe. Our respondents described the Lecturer faculty position in the UK as the equivalent of tenure-track Assistant Professor in the US but they are not on tenure-track. Our respondents described the Senior Lecturer position as equivalent to a tenured associate professor, on permanent contract but not a full professor yet. Data analysis for this table includes survey respondents with "0" number of mentors. | | | | |
|---|---|---|---|---|
| **Theme** | **Total Faculty number (n=446)** | **Female Faculty number (n=237)** | **Male Faculty number (n=201)** | **Non-Binary or Did not disclose gender Faculty number (n=8)** |
| **Assistant Professor (Tenure-track, non-Tenure-Track), Junior Group Leader (This term is mostly used across Europe))** | 63.2% (282 out of 446) | 63.3% (150 out of 237) | 63.7% (128 out of 201) | 50% (4 out of 8) |
| **Associate Professor (Tenured, Non-Tenured, Senior Group leader (this term is mostly used across Europe))** | 19.5% (87 out of 446) | 19.8% (47 out of 237) | 19.4% (39 out of 201) | 12.5% (1 out of 8) |
| **Lecturer (in the UK this is the equivalent of tenure-track Assistant Professor in the US), non-tenure** | 5.1% (23 out of 446) | 5.5% (13 out of 237) | 5% (10 out of 201) | 0% (0 out of 8) |

| | | | | |
|---|---|---|---|---|
| **track** | | | | |
| **Senior Lecturer (Equivalent to a tenured associate professor, on permanent contract but not a full professor yet), non-tenure track** | 2.1%<br>(9 out of 446) | 2.9%<br>(7 out of 237) | 1%<br>(2 out of 201) | 0%<br>(0 out of 8) |
| **Independently funded junior Research Fellow** | 2.2%<br>(10 out of 446) | 2.9%<br>(7 out of 237) | 1%<br>(2 out of 201) | 12.5%<br>(1 out of 8) |
| **Research Scientist/Staff Scientists/Research Professor (a research position higher than postdoctoral position but lower than tenure-track faculty position)** | 1.8%<br>(8 out of 446) | 0.84%<br>(2 out of 237) | 2.5%<br>(5 out of 201) | 12.5%<br>(1 out of 8) |
| **Full Professor (Tenured) (this is a position more senior than an Associate Professorship), senior group leader** | 6.05%<br>(27 out of 446) | 4.6%<br>(11 out of 237) | 7.5%<br>(15 out of 201) | 12.5%<br>(1 out of 8) |
| **Did not respond to this survey Question** | 2.4%<br>(11 out of 457) | 1.2%<br>(3 out of 240) | 3%<br>(6 out of 207) | 20%<br>(2 out of 10) |

| **Table S7. How frequently do you have formal meetings with your Mentor?**<br>**Analysis by gender,** Data analysis for this table excludes survey respondents with "0" number of mentors. | |
|---|---|
| **Theme** | **Frequency of Mentor-Mentee meetings** |
| **Total (n=364)** | 9.9% said Weekly (36 out of 364)<br>6% said Every 2 Weeks (22 out of 364)<br>12.6% said Monthly (46 out of 364)<br>53% said Scheduled as necessary (193 out of 364)<br>9.1% said Yearly (33 out of 364)<br>9.3% said Never (34 out of 364) |
| **Women Mentee (n=199)** | 9.5% said Weekly (19 out of 199)<br>4.5% said Every 2 Weeks (9 out of 199)<br>12% said Monthly (24 out of 199)<br>59.3% said Scheduled as necessary (118 out of 199)<br>6.5% said Yearly (13 out of 199)<br>8% said Never (16 out of 199) |
| **Men Mentee (n=159)** | 10.1% said Weekly (16 out of 159) |

| | |
|---|---|
| | 7.5% said Every 2 Weeks (12 out of 159)<br>13.8% said Monthly (22 out of 159)<br>44.7% said Scheduled as necessary (71 out of 159)<br>12.5% said Yearly (20 out of 159)<br>11.3% said Never (18 out of 159) |
| **Non-binary or did not disclose gender (n=6)** | 16.7% said Weekly (1 out of 6)<br>16.7% said Every 2 Weeks (1 out of 6)<br>0% said Monthly (0 out of 6)<br>66.6% said Scheduled as necessary (4 out of 6)<br>0% said Yearly (0 out of 6)<br>0% said Never (0 out of 6) |
| **Did not respond to this survey Question** | 21.7% (99 out of 457) |

**Table S7a. How frequently do you have formal meetings with your Mentor?**
**Analysis by continent,** Data analysis for this table excludes survey respondents with "0" number of mentors.

| Theme | Frequency of Mentor-Mentee meetings |
|---|---|
| **North America (n=145)** | 9% said Weekly (13 out of 145)<br>4.1% said Every 2 Weeks (6 out of 145)<br>11.7% said Monthly (17 out of 145)<br>51.7% said Scheduled as necessary (75 out of 145)<br>11% said Yearly (16 out of 145)<br>12.4% said Never (18 out of 145) |
| **Europe (n=116)** | 4.3% said Weekly (5 out of 116)<br>4.3% said Every 2 Weeks (5 out of 116)<br>14.7% said Monthly (17 out of 116)<br>62.1% said Scheduled as necessary (72 out of 116)<br>8.6% said Yearly (10 out of 116)<br>6% said Never (7 out of 116) |
| **Latin America+Asia+Oceania+Africa (n=97)** | 17.5% said Weekly (17 out of 97)<br>11.3% said Every 2 Weeks (11 out of 97)<br>11.3% said Monthly (11 out of 97)<br>44.3% said Scheduled as necessary (43 out of 97)<br>6.2% said Yearly (6 out of 97)<br>9.3% said Never (9 out of 97) |

**Table S7b. How frequently do you have formal meetings with your Mentor?**
**Analysis by mentorship initiation mode,** Data analysis for this table excludes survey respondents with "0" number of mentors.

| Theme | Frequency of Mentor-Mentee meetings |
|---|---|

| | |
|---|---|
| **Mentor was assigned to faculty mentee (n=106)** | 7.5% said Weekly (8 out of 106)<br>3.8% said Every 2 Weeks (4 out of 106)<br>16% said Monthly (17 out of 106)<br>54.7% said Scheduled as necessary (58 out of 106)<br>14.1% said Yearly (15 out of 106)<br>3.8% said Never (4 out of 106) |
| **Faculty mentee chose mentor voluntarily (n=258)** | 10.9% said Weekly (28 out of 258)<br>6.9% said Every 2 Weeks (18 out of 258)<br>11.2% said Monthly (29 out of 258)<br>52.3% said Scheduled as necessary (135 out of 258)<br>7% said Yearly (18 out of 258)<br>11.6% said Never (30 out of 258) |

**Table S7c. How frequently do you have formal meetings with your Mentor? Analysis by mentorship initiation mode and gender,** Data analysis for this table excludes survey respondents with "0" number of mentors.

| Theme | Quality of Mentor interactions with Mentee |
|---|---|
| **Women (n=139)<br>Chose mentor(s) voluntarily** | 10.1% said Weekly (14 out of 139)<br>5% said Every 2 Weeks (7 out of 139)<br>10.8% said Monthly (15 out of 139)<br>57.6% said Scheduled as necessary (80 out of 139)<br>6.5% said Yearly (9 out of 139)<br>10.1% said Never (14 out of 139) |
| **Women (n=60)<br>Were assigned to mentor(s)** | 8.3% said Weekly (5 out of 60)<br>3.3% said Every 2 Weeks (2 out of 60)<br>15% said Monthly (9 out of 60)<br>63.3% said Scheduled as necessary (38 out of 60)<br>6.7% said Yearly (4 out of 60)<br>3.3% said Never (2 out of 60) |
| **Men (n=114)<br>Chose mentor(s) voluntarily** | 11.4% said Weekly (13 out of 114)<br>8.8% said Every 2 Weeks (10 out of 114)<br>12.3% said Monthly (14 out of 114)<br>45.6% said Scheduled as necessary (52 out of 114)<br>7.9% said Yearly (9 out of 114)<br>14% said Never (16 out of 114) |
| **Men (n=45)<br>Were assigned to mentor(s)** | 6.7% said Weekly (3 out of 45)<br>2% said Every 2 Weeks (2 out of 45)<br>8% said Monthly (8 out of 45)<br>42.2% said Scheduled as necessary (19 out of 45)<br>24.4% said Yearly (11 out of 45)<br>4.4% said Never (2 out of 45) |
| **Non-binary or did not disclose gender<br>Chose mentor(s) voluntarily (n=5)** | 20% said Weekly (1 out of 5)<br>20% said Every 2 Weeks (1 out of 5)<br>0% said Monthly (0 out of 5)<br>60% said Scheduled as necessary (3 out of 5) |

| | |
|---|---|
| | 0% said Yearly (0 out of 5)<br>0% said Never (0 out of 5) |
| **Non-binary or did not disclose gender Were assigned to mentor(s) (n=1)** | 0% said Weekly (0 out of 1)<br>0% said Every 2 Weeks (0 out of 1)<br>0% said Monthly (0 out of 1)<br>100% said Scheduled as necessary (1 out of 1)<br>0% said Yearly (0 out of 1)<br>0% said Never (0 out of 1) |

| **Table S7d. How frequently do you have formal meetings with your Mentor? Analysis by mentorship initiation mode and geographic location,** Data analysis for this table excludes survey respondents with "0" number of mentors. | |
|---|---|
| **Theme** | **Quality of Mentor interactions with Mentee** |
| **North American Mentee Chose mentor(s) voluntarily (n=96)** | 10.4% said Weekly (10 out of 96)<br>3.1% said Every 2 Weeks (3 out of 96)<br>8.3% said Monthly (8 out of 96)<br>52.1% said Scheduled as necessary (50 out of 96)<br>8.3% said Yearly (8 out of 96)<br>17.7% said Never (17 out of 96) |
| **North American Mentee Were assigned to mentor(s) (n=49)** | 6.1% said Weekly (3 out of 49)<br>6.1% said Every 2 Weeks (3 out of 49)<br>18.4% said Monthly (9 out of 49)<br>51% said Scheduled as necessary (25 out of 49)<br>16.3% said Yearly (8 out of 49)<br>2% said Never (1 out of 49) |
| **European Mentee Chose mentor(s) voluntarily (n=79)** | 5.1% said Weekly (4 out of 79)<br>6.3% said Every 2 Weeks (5 out of 79)<br>13.9% said Monthly (11 out of 79)<br>59.5% said Scheduled as necessary (47 out of 79)<br>8.9% said Yearly (7 out of 79)<br>6.3% said Never (5 out of 79) |
| **European Mentee Were assigned to mentor(s) (n=37)** | 2.7% said Weekly (1 out of 37)<br>0% said Every 2 Weeks (0 out of 37)<br>16.2% said Monthly (6 out of 37)<br>67.6% said Scheduled as necessary (25 out of 37)<br>8.1% said Yearly (3 out of 37)<br>5.4% said Never (2 out of 37) |
| **Latin America+Asia+Africa+Oceania Chose mentor(s) voluntarily (n=77)** | 16.9% said Weekly (13 out of 77)<br>13% said Every 2 Weeks (10 out of 77)<br>9% said Monthly (9 out of 77)<br>45.5% said Scheduled as necessary (35 out of 77)<br>3.9% said Yearly (3 out of 77)<br>9.1% said Never (7 out of 77) |
| **Latin America+Asia+Africa+Oceania Were assigned to mentor(s)** | 21.1% said Weekly (4 out of 19)<br>5.3% said Every 2 Weeks (1 out of 19) |

| | |
|---|---|
| **(n=19)** | 10.5% said Monthly (2 out of 19)<br>42.1% said Scheduled as necessary (8 out of 19)<br>15.8% said Yearly (3 out of 19)<br>5.3% said Never (1 out of 19) |

| **Table S8. How were you originally matched to this Mentor?**<br>**Analysis by gender,** Data analysis for this table excludes survey respondents with "0" number of mentors. | |
|---|---|
| **Theme** | **Frequency of Mentor-Mentee meetings** |
| **Total (n=364)** | 70.9% Voluntarily (I chose the Mentor) (258 out of 364)<br>29.1% I was assigned to the Mentor (106 out of 364) |
| **Women Mentee (n=199)** | 69.8% Voluntarily (I chose the Mentor) (139 out of 199)<br>30.2% I was assigned to the Mentor (60 out of 199) |
| **Men Mentee (n=159)** | 71.7% Voluntarily (I chose the Mentor) (114 out of 159)<br>28.3% I was assigned to the Mentor (45 out of 159) |
| **Non-binary or did not disclose gender (n=6)** | 83.3% Voluntarily (I chose the Mentor) (5 out of 6)<br>16.7% I was assigned to the Mentor (1 out of 6) |

| **Table S9. What format do you use to communicate with your Mentor?**<br>Data analysis for this table excludes survey respondents with "0" number of mentors. | |
|---|---|
| **Theme** | **Format of Mentor-Mentee Communication** |
| **Total (n=363)** | **Use of a single approach Only:(n=108 or 30%)**<br>25.1% said Face-to-Face in-person (91 out of 363)<br>0.3% said Telephone (1 out of 363)<br>3.0% said E-mail (11 out of 363)<br>1.4% said Web/Video conferencing (e.g., Skype, Zoom) (5 out of 363)<br>0% said Social networking sites (0 out of 363)<br><br>**Use More than one approach:(n=255 or 70%)**<br>0.3% said Telephone+E-mail (1 out of 363)<br>36.8% said Face-to-Face+E-mail (135 out of 363)<br>2.4% said Face-to-Face+Telephone (9 out of 363)<br>0.3% said E-mail+Social networking sites (1 out of 363)<br>0.6% said E-mail+Web/Video conferencing (e.g., Skype, Zoom) (2 out of 363)<br>0.5% said Face-to-Face+Web/Video conferencing (e.g., Skype, Zoom) (2 out of 363)<br>2.9% said Face-to-Face+E-mail+Social networking sites (11 out of 363)<br>1.3% said Face-to-Face+E-mail+Text messaging (5 out of 363)<br>12.7% said Face-to-Face+Telephone+E-mail (46 out of 363)<br>0.5% said Telephone+E-mail+Web/Video conferencing (e.g., Skype, Zoom) (2 out of 363) |

| | |
|---|---|
| | 3.2% said Face-to-Face+E-mail+Web/Video conferencing (e.g., Skype, Zoom) (12 out of 363)<br>0.3% said Face-to-Face+E-mail+Web/Video conferencing (e.g., Skype, Zoom)+call/text messaging(1 out of 363)<br>0.3% said Face-to-Face+E-mail+Telephone+text messaging (1 out of 363)<br>1.6% said Face-to-Face+E-mail+Telephone+Web/Video conferencing (e.g., Skype, Zoom) (6 out of 363)<br>0.8% said Face-to-Face+E-mail+Telephone+Web/Video conferencing (e.g., Skype, Zoom)+Social networking sites (3 out of 363)<br>0.3% said Face-to-Face+Social networking sites (1 out of 363)<br>0.3% said Telephone+ Web/Video conferencing (e.g., Skype, Zoom) (1 out of 363)<br>0% said Telephone+Social networking sites+ Web/Video conferencing (e.g., Skype, Zoom) (0 out of 363)<br>0.5% said Face-to-Face+E-mail+Texting (2 out of 363)<br>0.3% said Face-to-Face+E-mail+Texting+Social networking sites (1 out of 363)<br>1% said Face-to-Face+E-mail+Web/Video conferencing (e.g., Skype, Zoom)+Social networking sites (4 out of 363)<br>0.6% said Face-to-Face+Telephone+E-mail+Social networking sites (2 out of 363)<br>0.3% said Face-to-Face+E-mail+Social networking sites (1 out of 363) |
| **Women Mentee (n=199)** | **Use of a single approach Only:(n=56 or 28.1%)**<br>22.6% said Face-to-Face in-person (45 out of 199)<br>0.5% said Telephone (1 out of 199)<br>3.5% said E-mail (7 out of 199)<br>1.5% said Web/Video conferencing (e.g., Skype, Zoom) (3 out of 199)<br>0% said Social networking sites (0 out of 199)<br><br>**Use More than one approach:(n=143 or 71.9%)**<br>0.5% said Telephone+E-mail (1 out of 199)<br>37.2% said Face-to-Face+E-mail (74 out of 199)<br>1.4% said Face-to-Face+Telephone (3 out of 199)<br>1.0% said E-mail+Social networking sites (2 out of 199)<br>0.5% said E-mail+Web/Video conferencing (e.g., Skype, Zoom) (1 out of 199)<br>0.5% said Face-to-Face+Web/Video conferencing (e.g., Skype, Zoom) (1 out of 199)<br>4.3% said Face-to-Face+E-mail+Social networking sites (9 out of 199)<br>2.4% said Face-to-Face+E-mail+Text messaging (5 out of 199)<br>12.6% said Face-to-Face+Telephone+E-mail (25 out of 199)<br>0.5% said Telephone+E-mail+Web/Video conferencing (e.g., Skype, Zoom) (1 out of 199)<br>3.5% said Face-to-Face+E-mail+Web/Video conferencing (e.g., Skype, Zoom) (7 out of 199)<br>0.5% said Face-to-Face+E-mail+Web/Video conferencing (e.g., Skype, Zoom)+call/text messaging(1 out of 199)<br>0.5% said Face-to-Face+E-mail+Telephone+text messaging (1 out of 199)<br>1.9% said Face-to-Face+E-mail+Telephone+Web/Video conferencing (e.g., Skype, Zoom) (4 out of 199)<br>1.4% said Face-to-Face+E-mail+Telephone+Web/Video conferencing (e.g., Skype, Zoom)+Social networking sites (3 out of 199) |
| **Men Mentee (n=158)** | **Use of a single approach Only:(n=50 or 31.6%)**<br>27.5% said Face-to-Face in-person (45 out of 158)<br>0% said Telephone (0 out of 158) |

| | |
|---|---|
| | 3.6% said E-mail (4 out of 158)<br>0.6% said Web/Video conferencing (e.g., Skype, Zoom) (1 out of 158)<br>0% said Social networking sites (0 out of 158)<br><br>**Use More than one approach:(n=108 or 68.3%)**<br><br>37.3% said E-mail+Face-to-Face (59 out of 158)<br>3.6% said Telephone+Face-to-Face (6 out of 158)<br>0.6% said E-mail+Web/Video conferencing (e.g., Skype, Zoom) (1 out of 158)<br>0.6% said Face-to-Face+Social networking sites (1 out of 158)<br>0.6% said Telephone+ Web/Video conferencing (e.g., Skype, Zoom) (1 out of 158)<br>0.6% said Telephone+E-mail+ Web/Video conferencing (e.g., Skype, Zoom) (1 out of 158)<br>0% said Telephone+Social networking sites+ Web/Video conferencing (e.g., Skype, Zoom) (0 out of 158)<br>0.6% said Face-to-Face+E-mail+Texting (2 out of 158)<br>0.6% said Face-to-Face+Web/Video conferencing (e.g., Skype, Zoom) (1 out of 158)<br>2.4% said Face-to-Face+E-mail+Web/Video conferencing (e.g., Skype, Zoom) (4 out of 158)<br>1.2% said Face-to-Face+E-mail+Social networking sites (2 out of 158)<br>0.6% said Face-to-Face+E-mail+Texting+Social networking sites (1 out of 158)<br>2.4% said Face-to-Face+E-mail+Web/Video conferencing (e.g., Skype, Zoom)+Social networking sites (4 out of 158)<br>12.6% said Face-to-Face+E-mail+Telephone (20 out of 158)<br>1.8% said Face-to-Face+Telephone+E-mail+Social networking sites (3 out of 158)<br>1.2% said Face-to-Face+E-mail+Telephone+Web/Video conferencing (e.g., Skype, Zoom) (2 out of 158) |
| **Non-Binary or Did not disclose Gender (n=6)** | **Use of a single approach Only:(n=2 or 33.3%)**<br><br>16.7% said Face-to-Face in-person (1 out of 6)<br>0% said Telephone (0 out of 6)<br>0% said E-mail (0 out of 6)<br>16.7% said Web/Video conferencing (e.g., Skype, Zoom) (1 out of 6)<br>0% said Social networking sites (0 out of 6)<br><br>**Use More than one approach:(n=4 or 66.7%)**<br><br>33.3% said Face-to-Face+E-mail (2 out of 6)<br>16.7% said Face-to-Face+E-mail+Social networking sites (1 out of 6)<br>16.7% said Face-to-Face+E-mail+Web/Video conferencing (e.g., Skype, Zoom) (1 out of 6) |

**Table S10. Do you find your interactions with your Mentor constructive? by gender**
Respondents (Faculty Mentees) rating the constructive quality of their interactions with their Mentor.
Data analysis for this table excludes survey respondents with "0" number of mentors.

| Theme | Quality of Mentor interactions with Mentee |
|---|---|
| **Total (n=365)** | 78.1% said Yes (285 out of 365)<br>3.6% said No (13 out of 365)<br>18.3% said Sometimes (67 out of 365) |

| | |
|---|---|
| **Women Mentee (n=199)** | 74.9% said Yes (149 out of 199)<br>3.5% said No (7 out of 199)<br>21.6% said Sometimes (43 out of 199) |
| **Men Mentee (n=161)** | 82% said Yes (132 out of 161)<br>3.7% said No (6 out of 161)<br>14.3% said Sometimes (23 out of 161) |
| **Non-binary or did not disclose gender (n=5)** | 80% said Yes (4 out of 5)<br>0% said No (0 out of 5)<br>20% said Sometimes (1 out of 5) |

**Table S10a. Do you find your interactions with your Mentor constructive? by continent** Respondents (Faculty Mentees) rating the constructive quality of their interactions with their Mentor. Data analysis for this table excludes survey respondents with "0" number of mentors.

| Theme | Quality of Mentor interactions with Mentee |
|---|---|
| **North America (n=147)** | 81.6% said Yes (120 out of 147)<br>0.7% said No (1 out of 147)<br>17.7% said Sometimes (26 out of 147) |
| **Europe (n=116)** | 76.7% said Yes (89 out of 116)<br>6.9% said No (8 out of 116)<br>16.4% said Sometimes (19 out of 116) |
| **Latin America+Asia+Africa+Oceania (n=96)** | 75% said Yes (72 out of 96)<br>3.1% said No (3 out of 96)<br>21.9% said Sometimes (21 out of 96) |

**Table S10b. Do you find your interactions with your Mentor constructive? Analysis by mentorship initiation mode,** Data analysis for this table excludes survey respondents with "0" number of mentors.

| Theme | Quality of Mentor interactions with Mentee |
|---|---|
| **Mentor was assigned to faculty mentee (n=106)** | 68.9% said Yes (73 out of 106)<br>6.6% said No (7 out of 106)<br>24.5% said Sometimes (26 out of 106) |
| **Faculty mentee chose mentor voluntarily (n=259)** | 81.9% said Yes (212 out of 259)<br>2.3% said No (6 out of 259)<br>15.8% said Sometimes (41 out of 259) |

| **Table S10c. Do you find your interactions with your Mentor constructive? by gender and method of choosing mentor,** Data analysis for this table excludes survey respondents with "0" number of mentors. | |
|---|---|
| **Theme** | **Quality of Mentor interactions with Mentee** |
| **Women Mentee**<br>**Chose mentor(s) voluntarily (n=139)** | 78.4% said Yes (109 out of 139)<br>2% said No (3 out of 139)<br>19.4% said Sometimes (27 out of 139) |
| **Women Mentee**<br>**Were assigned to mentor(s) (n=60)** | 66.7% said Yes (40 out of 60)<br>6.7% said No (4 out of 60)<br>26.6% said Sometimes (16 out of 60) |
| **Men Mentee**<br>**Chose mentor(s) voluntarily (n=114)** | 86.8% said Yes (99 out of 114)<br>2.6% said No (3 out of 114)<br>10.5% said Sometimes (12 out of 114) |
| **Men Mentee**<br>**Were assigned to mentor(s) (n=46)** | 71.7% said Yes (33 out of 46)<br>6.5% said No (3 out of 46)<br>21.7% said Sometimes (10 out of 46) |
| **Non-binary or did not disclose gender**<br>**Chose mentor(s) voluntarily (n=6)** | 66.7% said Yes (4 out of 6)<br>0% said No (0 out of 6)<br>33.3% said Sometimes (2 out of 6) |
| **Non-binary or did not disclose gender**<br>**Were assigned to mentor(s) (n=0)** | 0% said Yes (0 out of 0)<br>0% said No (0 out of 0)<br>0% said Sometimes (0 out of 0) |

| **Table S10d. Do you find your interactions with your Mentor constructive? Analysis by gender and method of choosing mentor,** Data analysis for this table excludes survey respondents with "0" number of mentors. | |
|---|---|
| **Theme** | **Quality of Mentor interactions with Mentee** |
| **North American Mentee**<br>**Chose mentor(s) voluntarily (n=98)** | 83.7% said Yes (82 out of 98)<br>1% said No (1 out of 98)<br>15.3% said Sometimes (52 out of 98) |
| **North American Mentee**<br>**Were assigned to mentor(s) (n=49)** | 77.6% said Yes (38 out of 49)<br>0% said No (0 out of 49)<br>22.4% said Sometimes (11 out of 49) |
| **European Mentee**<br>**Chose mentor(s) voluntarily (n=79)** | 86.8% said Yes (68 out of 79)<br>2.6% said No (2 out of 79)<br>11.4% said Sometimes (9 out of 79) |
| **European Mentee**<br>**Were assigned to mentor(s) (n=37)** | 56.8% said Yes (21 out of 37)<br>16.2% said No (6 out of 37)<br>27% said Sometimes (10 out of 37) |
| **Latin America+Asia+Africa+Oceania** | 76.6% said Yes (59 out of 77) |

| | |
|---|---|
| **Chose mentor(s) voluntarily (n=77)** | 2.6% said No (2 out of 77)<br>20.8% said Sometimes (16 out of of 77) |
| **Latin America+Asia+Africa+Oceania Were assigned to mentor(s) (n=19)** | 68.4% said Yes (13 out of 19)<br>5.3% said No (1 out of 19)<br>26.3% said Sometimes (5 out of 19) |

**Table S11. Method of original mentee-mentor matching**
Method of original Mentee (early- and mid-career faculty)-Mentor (senior faculty colleagues) matching, Data analysis for this table excludes survey respondents with "0" number of mentors.

| Theme | Original Matching |
|---|---|
| **Total (n=381)** | 70.7% said Voluntary (I chose Mentor) (258 out of 365)<br>29.3% said I was assigned to my Mentor (107 out of 365) |
| **Women (n=199)** | 69.8% said Voluntary (I chose Mentor) (139 out of 199)<br>30.2% said I was assigned to my Mentor (60 out of 199) |
| **Men (n=160)** | 71.3% said Voluntary (I chose Mentor) (114 out of 160)<br>28.7% said I was assigned to my Mentor (46 out of 160) |
| **Non-binary or did not disclose gender (n=6)** | 83.3% said Voluntary (I chose Mentor) (5 out of 6)<br>16.7% said I was assigned to my Mentor (1 out of 6) |

**Table S12. Number of mentors that Faculty Mentee had**
All percentages are calculated out of the total number of respondents to this question n=457. The results show that 17% of female faculty had no mentors compared to 21.9% of male faculty. Data analysis for this table includes survey respondents with "0" number of mentors.

| Theme | Number of mentors |
|---|---|
| **Total (n=456)** | 20% had 0 mentors (91 out of 456)<br>27.2% had 1 mentor (124 out of 456)<br>23.7% had 2 mentors (108 out of 456)<br>14% had 3 mentors (64 out of 456)<br>4.6% had 4 mentors (21 out of 456)<br>3.3% had 5 mentors (15 out of 456)<br>7.2% had >5 mentors (33 out of 456)<br>Average=2<br>Median=1 |
| **Women (n=240)** | 17.1% had 0 mentors (41 out of 240)<br>25.4% had 1 mentor (61 out of 240)<br>27.1% had 2 mentors (65 out of 240)<br>13.3% had 3 mentors (32 out of 240)<br>4.6% had 4 mentors (11 out of 240)<br>2.9% had 5 mentors (7 out of 240) |

| | |
|---|---|
| | 9.6% had >5 mentors (23 out of 240)<br>Average=2<br>Median=2 |
| **Men (n=207)** | 22.7% had 0 mentors (47 out of 207)<br>29.5% had 1 mentor (61 out of 207)<br>19.3% had 2 mentors (40 out of 207)<br>15% had 3 mentors (31 out of 207)<br>4.8% had 4 mentors (10 out of 207)<br>3.9% had 5 mentors (8 out of 207)<br>4.8% had >5 mentors (10 out of 207)<br>Average=2<br>Median=1 |
| **Non-binary or preferred not to disclose gender (n=9)** | 30% had 0 mentors (3 out of 9)<br>20% had 1 mentor (2 out of 9)<br>30% had 2 mentors (3 out of 9)<br>10% had 3 mentors (1 out of 9)<br>0% had 4 mentors (0 out of 9)<br>0% had 5 mentors (0 out of 9)<br>0% had >5 mentors (0 out of 9)<br>Average=1<br>Median=1 |
| **Did not respond to this survey Question** | 0.22% (1 out of 457) |

**Table S12a. Number of mentors that Faculty Mentee had by country/region**
All percentages are calculated out of the total number of respondents to this question n=449. Data analysis for this table includes survey respondents with "0" number of mentors.

| Theme | Number of mentors |
|---|---|
| **Total (n=449)** | 18.9% had 0 mentors (85 out of 449)<br>26.3% had 1 mentor (118 out of 449)<br>22.9% had 2 mentors (103 out of 449)<br>14.3% had 3 mentors (64 out of 449)<br>4.7% had 4 mentors (21 out of 449)<br>3.3% had 5 mentors (15 out of 449)<br>7.3% had >5 mentors (33 out of 449)<br>Average= 1<br>Median= 2 |
| **United States (n=156)** | 10.9% had 0 mentors (17 out of 156)<br>13.5% had 1 mentor (21 out of 156)<br>21.1% had 2 mentors (33 out of 156)<br>23.1% had 3 mentors (36 out of 156)<br>10.9% had 4 mentors (17 out of 156)<br>6.4% had 5 mentors (10 out of 156)<br>14.1% had >5 mentors (22 out of 156) |

| | |
|---|---|
| | Average= 2.9<br>Median= 3 |
| **United Kingdom (n=72)** | 19.4% had 0 mentors (14 out of 72)<br>31.9% had 1 mentor (23 out of 72)<br>27.8% had 2 mentors (20 out of 72)<br>11.1% had 3 mentors (8 out of 72)<br>0% had 4 mentors (0 out of 72)<br>4.2% had 5 mentors (3 out of 72)<br>5.5% had >5 mentors (4 out of 72)<br>Average= 1.8<br>Median= 1 |
| **North America (n=165)** | 11.4% had 0 mentors (19 out of 167)<br>13.2% had 1 mentor (22 out of 167)<br>22.1% had 2 mentors (37 out of 167)<br>22.8% had 3 mentors (38 out of 167)<br>10.8% had 4 mentors (18 out of 167)<br>6.6% had 5 mentors (11 out of 167)<br>13.2% had >5 mentors (22 out of 167)<br>Average= 2.8<br>Median= 3 |
| **Europe (n=158)** | 26.6% had 0 mentors (42 out of 158)<br>34.8% had 1 mentor (55 out of 158)<br>22.7% had 2 mentors (36 out of 158)<br>7.6% had 3 mentors (12 out of 158)<br>1.9% had 4 mentors (3 out of 158)<br>2.5% had 5 mentors (4 out of 158)<br>3.8% had >5 mentors (6 out of 158)<br>Average= 1.4<br>Median= 1 |
| **Africa+Asia+Oceania+Latin America (n=124)** | 22.5% had 0 mentors (28 out of 124)<br>36.2% had 1 mentor (45 out of 124)<br>21.8% had 2 mentors (27 out of 124)<br>10.5% had 3 mentors (13 out of 124)<br>1.6% had 4 mentors (2 out of 124)<br>2.4% had 5 mentors (3 out of 124)<br>4.8% had >5 mentors (6 out of 124)<br>Average= 1.5<br>Median= 1 |
| **Did not specify their country of research (n=10)** | 30% had 0 mentors (3 out of 10)<br>40% had 1 mentor (4 out of 10)<br>30% had 2 mentors (3 out of 10)<br>0% had 3 mentors (0 out of 10)<br>0% had 4 mentors (0 out of 10)<br>0% had 5 mentors (0 out of 10)<br>0% had >5 mentors (0 out of 10)<br>Average= 1<br>Median= 1 |

| **Table S13. Number of years mentee has worked with mentor(s)**<br>All percentages are calculated out of the total number of respondents to this question n=365. Data analysis for this table excludes survey respondents with "0" number of mentors. | |
|---|---|
| **Theme** | **Number of years of interactions** |
| **Total (n=365)** | 17.7% Less than one year (65 out of 365)<br>24.6% 1-2 years (90 out of 365)<br>21.3% 3-4 years (78 out of 365)<br>36.2% 5 or more years (132 out of 365) |
| **Women (n=199)** | 23.1% Less than one year (46 out of 199)<br>23.1% 1-2 years (46 out of 199)<br>19.6% 3-4 years (39 out of 199)<br>34.2% 5 or more years (68 out of 199) |
| **Men (n=160)** | 11.9% Less than one year (19 out of 160)<br>25.6% 1-2 years (41 out of 160)<br>23.8% 3-4 years (38 out of 160)<br>38.8% 5 or more years (62 out of 160) |
| **Non-binary or did not disclose gender (n=6)** | 0% Less than one year (0 out of 6)<br>50% 1-2 years (3 out of 6)<br>16.7% 3-4 years (1 out of 6)<br>33.3% 5 or more years (2 out of 6) |
| **Did not respond to this survey Question** | 20.1% (92 out of 457) |

| **Table S14. Respondent Age distribution**<br>Data analysis for this table includes survey respondents with "0" number of mentors. | |
|---|---|
| **Theme** | **Respondent Age** |
| **Total (n=428)** | Average = 39 years<br>Median = 38 years |
| **Women (n=226)** | Average = 38.9 years<br>Median =38 years |
| **Men (n=196)** | Average = 39.2 years<br>Median =39 years |
| **Minimum** | 28 years |
| **Maximum** | 65 years |
| **Non-binary or did not disclose gender(n=3)** | Average = 36 years<br>Median = 36 years |
| **Preferred not to say (n=3)** | 0.7% (3 out of 457) |
| **Did not respond to this survey Question** | 6.3% (29 out of 457) |

| **Table S15. Respondents (Faculty Mentees) rating of the quality of the match between mentee and mentor,** Data analysis for this table excludes survey respondents with "0" number of mentors. | |
|---|---|
| **Theme** | **Quality of the match between Mentee and Mentor** |
| **Total (n=364)** | 33.5% said Excellent (122 out of 364)<br>39% said Very Good (142 out of 364)<br>16.2% said Good (59 out of 364)<br>8% said Fair (29 out of 364)<br>3.3% said Poor (12 out of 364) |
| **Women Mentee (n=199)** | 34.2% said Excellent (68 out of 199)<br>36.7% said Very Good (73 out of 199)<br>15.6% said Good (31 out of 199)<br>11.1% said Fair (22 out of 199)<br>2.5% said Poor (5 out of 199) |
| **Men Mentee (n=159)** | 32.1% said Excellent (51 out of 159)<br>42.1% said Very Good (67 out of 159)<br>17.6% said Good (28 bout of 159)<br>4.4% said Fair (7 out of 159)<br>3.8% said Poor (6 out of 159) |
| **Non-binary or did not disclose gender (n=6)** | 50% said Excellent (3 out of 6)<br>33.3% said Very Good (2 out of 6)<br>0% said Good (0 out of 6)<br>0% said Fair (0 out of 6)<br>16.7% said Poor (1 out of 6) |

| **Table S15a. Respondents (Faculty Mentees) rating of the quality of the match between mentee and mentor by mentorship initiation mode,** Data analysis for this table excludes survey respondents with "0" number of mentors. | |
|---|---|
| **Theme** | **Quality of the match between Mentee and Mentor** |
| **Mentor was assigned to faculty Mentee (n=106)** | 23.6% said Excellent (25 out of 106)<br>35.8% said Very Good (38 out of 106)<br>19.8% said Good (21 out of 106)<br>12.3% said Fair (13 out of 106)<br>8.5% said Poor (9 out of 106) |
| **Faculty mentee chose mentor voluntarily (n=258)** | 37.5% said Excellent (97 out of 258)<br>40.3% said Very Good (104 out of 258)<br>14.7% said Good (38 out of 258)<br>6.2% said Fair (16 out of 258)<br>1.2% said Poor (3 out of 258) |

| **Table S15b. Respondents (Faculty Mentees) rating of the quality of the match between mentee and mentor by country/region,** Data analysis for this table excludes survey respondents with "0" number of mentors. Analysis performed for North America versus Europe versus all other continents (aggregate data on Asia+Africa+Oceania+Latin America) representing three comparable sample sizes. | |
|---|---|
| **Theme** | **Quality of the match between Mentee and Mentor** |
| **United States (n=134)** | 37.3% said Excellent (50 out of 134)<br>35.8% said Very Good (48 out of 134)<br>20.1% said Good (27 out of 134)<br>7.5% said Fair (10 out of 134)<br>1.5% said Poor (2 out of 134) |
| **United Kingdom (n=57)** | 28% said Excellent (16 out of 57)<br>33.3% said Very Good (19 out of 57)<br>21% said Good (12 out of 57)<br>12.3% said Fair (7 out of 57)<br>5.3% said Poor (3 out of 57) |
| **North America (n=144)** | 37.5% said Excellent (54 out of 144)<br>34% said Very Good (49 out of 144)<br>20.1% said Good (29 out of 144)<br>6.9% said Fair (10 out of 144)<br>1.4% said Poor (2 out of 144) |
| **Europe (n=117)** | 31.6% said Excellent (37 out of 117)<br>38.5% said Very Good (45 out of 117)<br>17.1% said Good (20 out of 117)<br>8.5% said Fair (10 out of 117)<br>4.3% said Poor (5 out of 117) |
| **Latin America+Africa+Asia+Oceania (n=97)** | 30.1% said Excellent (30 out of 97)<br>46.4% said Very Good (45 out of 97)<br>10.3% said Good (10 out of 97)<br>9.3% said Fair (9 out of 97)<br>3.1% said Poor (3 out of 97) |

| **Table S15c. Respondents (Faculty Mentees) rating of the quality of the match between mentee and mentor, Analysis by mentorship initiation mode and gender,** Data analysis for this table excludes survey respondents with "0" number of mentors. | |
|---|---|
| **Theme** | **Quality of Mentor interactions with Mentee** |
| **Women<br>Chose mentor(s) voluntarily (n=139)** | 37.4% said Excellent (52 out of 139)<br>38.8% said Very Good (54 out of 139)<br>14.4% said Good (20 out of 139)<br>8.6% said Fair (12 out of 139)<br>0.7% said Poor (1 out of 139) |
| **Women<br>Were assigned to mentor(s) (n=60)** | 26.7% said Excellent (16 out of 60)<br>31.7% said Very Good (19 out of 60) |

| | |
|---|---|
| | 18.3% said Good (11 out of 60)<br>16.7% said Fair (10 out of 60)<br>6.7% said Poor (4 out of 60) |
| **Men**<br>**Chose mentor(s) voluntarily (n=114)** | 36.8% said Excellent (42 out of 114)<br>43% said Very Good (49 out of 114)<br>15.8% said Good (18 out of 114)<br>3.5% said Fair (4 out of 114)<br>0.9% said Poor (1 out of 114) |
| **Men**<br>**Were assigned to mentor(s) (n=45)** | 20% said Excellent (9 out of 45)<br>40% said Very Good (18 out of 45)<br>22.2% said Good (10 out of 45)<br>6.7% said Fair (3 out of 45)<br>11.1% said Poor (5 out of 45) |
| **Non-binary or did not disclose gender**<br>**Chose mentor(s) voluntarily (n=1)** | 0% said Excellent (0 out of 1)<br>100% said Very Good (1 out of )1<br>0% said Good (0 out of 1)<br>0% said Fair (0 out of 1)<br>0% said Poor (0 out of 1) |
| **Non-binary or did not disclose gender**<br>**Were assigned to mentor(s) (n=5)** | 60% said Excellent (3 out of 5)<br>20% said Very Good (1 out of )5<br>0% said Good (0 out of 5)<br>0% said Fair (0 out of 5)<br>20% said Poor (1 out of 5) |

**Table S15d. Respondents (Faculty Mentees) rating of the quality of the match between mentor and mentee, Analysis by mentorship initiation mode and geographic location,** Data analysis for this table excludes survey respondents with "0" number of mentors.

| Theme | Quality of Mentor interactions with Mentee |
|---|---|
| **North American Mentee**<br>**Chose mentor(s) voluntarily (n=97)** | 42.3% said Excellent (41 out of 97)<br>30.9% said Very Good (30 out of 97)<br>20.6% said Good (20 out of 97)<br>6.2% said Fair (6 out of 97)<br>0% said Poor (0 out of 97) |
| **North American Mentee**<br>**Were assigned to mentor(s) (n=49)** | 28.6% said Excellent (14 out of 49)<br>40.8% said Very Good (20 out of 49)<br>18.4% said Good (9 out of 49)<br>8.2% said Fair (4 out of 49)<br>4.1% said Poor (2 out of 49) |
| **European Mentee**<br>**Chose mentor(s) voluntarily (n=79)** | 39.2% said Excellent (31 out of 79)<br>43% said Very Good (34 out of 79)<br>12.7% said Good (10 out of 79)<br>5.1% said Fair (4 out of 79)<br>0% said Poor (0 out of 79) |

| | |
|---|---|
| **European Mentee**<br>**Were assigned to mentor(s) (n=37)** | 16.2% said Excellent (6 out of 37)<br>29.7% said Very Good (11 out of 37)<br>24.3% said Good (9 out of 37)<br>16.2% said Fair (6 out of 37)<br>13.5% said Poor (5 out of 37) |
| **Latin America+Africa+Asia+Oceania**<br>**Chose mentor(s) voluntarily (n=77)** | 32.5% said Excellent (25 out of 77)<br>48.1% said Very Good (37 out of 77)<br>10.4% said Good (8 out of 77)<br>7.8% said Fair (6 out of 77)<br>1.3% said Poor (1 out of 77) |
| **Latin America+Africa+Asia+Oceania**<br>**Were assigned to mentor(s) (n=19)** | 26.3% said Excellent (5 out of 19)<br>36.8% said Very Good (7 out of 19)<br>10.5% said Good (2 out of 19)<br>15.8% said Fair (3 out of 19)<br>10.5% said Poor (2 out of 19) |

**Table S16. Do you maintain contact with your former mentors? by gender**
Respondents (Faculty Mentees) maintaining contact with former mentor(s). Data analysis for this table includes survey respondents with "0" number of mentors.

| Theme | Mentee relationship with former Mentors |
|---|---|
| **Total (n=385)** | 78.4% said Yes (302 out of 385)<br>21.5% said No (83 out of 385) |
| **Women Mentee (n=209)** | 76.1% said Yes (159 out of 209)<br>23.9% said No (50 out of 209) |
| **Men Mentee (n=169)** | 82.8% said Yes (140 out of 169)<br>17.1% said No (29 out of 169) |
| **Non-binary or did not disclose gender (n=7)** | 42.8% said Yes (3 out of 7)<br>57.1% said No (4 out of 7) |
| **Did not respond to this survey Question** | 15.7% (72 out of 457) |

**Table S16a. Do you maintain contact with your former mentors? by continent**
Respondents (Faculty Mentees) maintaining contact with former mentor(s). Data analysis for this table includes survey respondents with "0" number of mentors.

| Theme | Mentee relationship with former Mentors |
|---|---|
| **North America (n=153)** | 84.3% said Yes (129 out of 153)<br>15.7% said No (24 out of 153) |
| **Europe (n=126)** | 69% said Yes (87 out of 126)<br>31% said No (39 out of 126) |
| **Latin America+Africa+Oceania+Asia (n=110)** | 79.1% said Yes (87 out of 110) |

| | |
|---|---|
| | 20.9% said No (23 out of 110) |
| **Did not respond to this survey Question (n=68)** | 15.1% (68 out of 450) |

| **Table S17. Is your Mentor physically accessible? (work in the same campus or same building, so that you have unplanned meetings or encounters), Analysis by gender,** Data analysis for this table excludes survey respondents with "0" number of mentors. | |
|---|---|
| **Theme** | **Mentor accessible to mentee** |
| **Total (n=364)** | 77.5% said Yes (282 out of 364)<br>22.5% said No (82 out of 364) |
| **Women Mentee (n=199)** | 73.4% said Yes (146 out of 199)<br>26.6% said No (53 out of 199) |
| **Men Mentee (n=159)** | 83% said Yes (132 out of 159)<br>17% said No (27 out of 159) |
| **Non-binary or did not disclose gender (n=6)** | 66.7% said Yes (4 out of 6)<br>33.3% said No (2 out of 6) |

| **Table S17a. Mentor Location same department or institution as faculty mentee Responses included “Yes” or “No” only. Analysis by gender,** Data analysis for this table excludes survey respondents with "0" number of mentors. | |
|---|---|
| **Theme** | **Mentor location** |
| **Total (n=365)** | 74.8% said Yes (283 out of 365)<br>25.1% said No (82 out of 365) |
| **Women (n=199)** | 75.2% said Yes (151 out of 199)<br>24.8% said No (48 out of 199) |
| **Men (n=160)** | 76.2% said Yes (129 out of 160)<br>23.8% said No (31 out of 160) |
| **Non-binary or did not disclose gender (n=6)** | 27.5% said Yes (3 out of 6)<br>62.5% said No (3 out of 6) |

| **Table S17b. Mentor accessibility by mode of mentorship initiation: Is your Mentor physically accessible? (work in the same campus or same building) by mentorship initiation mode,** Data analysis for this table excludes survey respondents with "0" number of mentors. | |
|---|---|
| **Theme** | **Mentor accessible to mentee** |
| **Total (n=364)** | 77.5% said Yes (282 out of 364) |

| | |
|---|---|
| | 22.5% said No (82 out of 364) |
| **Mentees who chose mentor voluntarily (n=258)** | 73.6% said Yes (190 out of 258)<br>26.4% said No (68 out of 258) |
| **Mentees who were assigned to mentor (n=106)** | 86.8% said Yes (92 out of 106)<br>13.2% said No (14 out of 106) |

**Table S18. Respondent Citizenship status in country of research (country of faculty appointment),** Data analysis for this table includes survey respondents with "0" number of mentors.

| Theme | Citizen |
|---|---|
| **Total (n=461)** | 72.3% said Yes (333 out of 461)<br>27.7% said No (128 out of 461) |
| **Women (n=247)** | 74.5% said Yes (184 out of 247)<br>25.5% said No (63 out of 247) |
| **Men (n=210)** | 70% said Yes (147 out of 210)<br>30% said No (63 out of 210) |
| **Non-binary and did not respond to this survey Question (n=6)** | 71.4% said Yes (5 out of 7)<br>28.6% said No (2 out of 7) |

**Table S19. Does the institution you are hired at as an independent investigator provide a faculty mentoring program to mentor you on your own career? by gender**
Data analysis for this table includes survey respondents with "0" number of mentors.

| Theme | Institutional Faculty Mentoring program |
|---|---|
| **Total (n=410)** | 48.8% said Yes (200 out of 410)<br>51.2% said No (210 out of 410) |
| **Women Mentee (n=224)** | 50.4% said Yes (113 out of 224)<br>49.6% said No (111 out of 224) |
| **Men Mentee (n=179)** | 46.9% said Yes (84 out of 179)<br>53.1% said No (95 out of 179) |
| **Non-binary or did not disclose gender (n=7)** | 42.9% said Yes (3 out of 7)<br>57.1% said No (4 out of 7) |
| **Did not respond to this survey Question** | 10.3% (47 out of 457) |

**Table S19a. Does the institution you are hired at as an independent investigator provide a faculty mentoring program to mentor you on your own career? by continent**

| Data analysis for this table includes survey respondents with "0" number of mentors. | |
|---|---|
| **Theme** | **Institutional Faculty Mentoring program** |
| **North America (n=157)** | 69.4% said Yes (109 out of 157)<br>30.6% said No (48 out of 157) |
| **Europe (n=136)** | 46.3% said Yes (63 out of 136)<br>53.7% said No (73 out of 136) |
| **Latin America+Africa+Asia+Oceania (n=112)** | 20.5% said Yes (23 out of 112)<br>79.5% said No (89 out of 112) |
| **Did not respond to this survey Question** | 11.5% (52 out of 450) |

| **Table S20. Faculty Mentee participation in any form of formal (run by institutions) or informal peer-mentorship schemes, analysis by gender.** Data analysis for this table includes survey respondents with "0" number of mentors. | |
|---|---|
| **Theme** | **Availability of formal Mentors to Mentees** |
| **Total (n=406)** | 42.1% said Yes (171 out of 406)<br>57.9% said No (235 out of 406) |
| **Women Mentee(n=223)** | 48.4% said Yes (108 out of 223)<br>51.6% said No (115 out of 223) |
| **Men Mentee(n=176)** | 35.2% said Yes (62 out of 176)<br>64.8% said No (114 out of 176) |
| **Non-binary or did not disclose gender (n=7)** | 14.3% said Yes (1 out of 7)<br>85.7% said No (6 out of 7) |
| **Did not respond to this survey Question** | 11.1% (51 out of 457) |

| **Table S20a. Faculty Mentee participation in any form of formal (run by institutions) or informal peer-mentorship programs, analysis by continent,** Data analysis for this table includes survey respondents with "0" number of mentors. | |
|---|---|
| **Theme** | **Availability of formal Mentors to Mentees** |
| **North America (n=157)** | 47.1% said Yes (74 out of 157)<br>52.9% said No (83 out of 157) |
| **Europe (n=135)** | 47.4% said Yes (64 out of 135)<br>52.6% said No (71 out of 135) |
| **Latin America+Asia+Africa+Oceania (n=111)** | 27% said Yes (30 out of 111)<br>73% said No (81 out of 111) |

| | |
|---|---|
| **Did not respond to this survey Question** | 12% (54 out of 450) |

| **Table S21. Mentor gender same as mentee or not**<br>Responses included "Yes" or "No" only. Data analysis for this table excludes survey respondents with "0" number of mentors. | |
|---|---|
| **Theme** | **Mentor gender same as mentee** |
| **Total (n=364)** | 53.6% said Yes (195 out of 364)<br>46.4% said No (169 out of 364) |
| **Women Mentee(n=199)** | 40.7% said Yes (81 out of 199)<br>59.3% said No (118 out of 199) |
| **Men Mentee(n=159)** | 71.1% said Yes (113 out of 159)<br>28.9% said No (46 out of 159) |
| **Non-binary or did not disclose gender (n=6)** | 16.7% said Yes (1 out of 6)<br>83.3% said No (5 out of 6) |
| **Did not respond to this survey Question** | 20.3% (93 out of 457) |

| **Table S22. Do you feel happy & satisfied about your current research and position as an independent investigator? Analysis by gender,** Data analysis for this table includes survey respondents with "0" number of mentors. | |
|---|---|
| **Theme** | **Mentee satisfaction with current research program** |
| **Total (n=368)** | 73.6% said Yes (271 out of 368)<br>26.4% said No (97 out of 368) |
| **Women Mentee (n=194)** | 67% said Yes (130 out of 194)<br>33% said No (64 out of 194) |
| **Men Mentee (n=169)** | 81.7% said Yes (138 out of 169)<br>18.3% said No (31 out of 169) |
| **Non-binary or did not disclose gender (n=5)** | 60% said Yes (3 out of 5)<br>40% said No (2 out of 5) |
| **Did not respond to this survey Question** | 19.5% (89 out of 457) |

| **Table S22a. Do you feel happy & satisfied about your current research and position as an independent investigator? Analysis by mentorship initiation mode,** Data analysis for this table includes survey respondents with "0" number of mentors. | |
|---|---|
| **Theme** | **Mentee satisfaction with current research program** |
| **Total (n=327)** | 78.3% said Yes (256 out of 327) |

| | |
|---|---|
| | 21.7% said No (71 out of 327) |
| **Mentee chose mentor voluntarily (n=233)** | 78.5% said Yes (183 out of 233)<br>21.5% said No (50 out of 233) |
| **Mentee was assigned a mentor (n=94)** | 77.7% said Yes (73 out of 94)<br>22.3% said No (21 out of 94) |

**Table S22b. Do you feel happy & satisfied about your current research and position as an independent investigator? Analysis by geographic region (continent),** Data analysis for this table includes survey respondents with "0" number of mentors.

| **Theme** | **Mentee satisfaction with current research program** |
|---|---|
| **North America (n=137)** | 78.8% said Yes (108 out of 137)<br>21.2% said No (29 out of 137) |
| **Europe (n=117)** | 71.8% said Yes (84 out of 117)<br>28.2% said No (33 out of 117) |
| **Latin America+Oceania+Africa+Asia (n=106)** | 69.8% said Yes (74 out of 106)<br>30.2% said No (32 out of 106) |

**Table S23. Do you feel optimistic about the future? Analysis by gender,** Data analysis for this table includes survey respondents with "0" number of mentors.

| **Theme** | **Mentee optimism about the future** |
|---|---|
| **Total (n=367)** | 78.7% said Yes (289 out of 367)<br>21.3% said No (78 out of 367) |
| **Women Mentee (n=195)** | 72.8% said Yes (142 out of 195)<br>27.2% said No (53 out of 195) |
| **Men Mentee (n=167)** | 86.2% said Yes (144 out of 167)<br>13.8% said No (23 out of 167) |
| **Non-binary or did not disclose gender (n=5)** | 60% said Yes (3 out of 5)<br>40% said No (2 out of 5) |
| **Did not respond to this survey Question** | 19.7% (90 out of 457) |

**Table S23a. Do you feel optimistic about the future?**
**Analysis by mentorship initiation mode,** Data analysis for this table includes survey respondents with "0" number of mentors.

| **Theme** | **Mentee Optimism about the future** |
|---|---|

| | |
|---|---|
| **Total (n=326)** | 81.6% said Yes (266 out of 326)<br>18.4% said No (60 out of 326) |
| **Mentee chose mentor voluntarily (n=232)** | 81.5% said Yes (189 out of 232)<br>18.5% said No (43 out of 232) |
| **Mentee was assigned a mentor (n=94)** | 81.9% said Yes (77 out of 94)<br>18.1% said No (17 out of 94) |

**Table S23b. Do you feel optimistic about the future?**
**Analysis by geographic region (continent),** Data analysis for this table includes survey respondents with "0" number of mentors.

| Theme | Mentee Optimism about the future |
|---|---|
| **North America (n=136)** | 88.2% said Yes (120 out of 136)<br>11.8% said No (16 out of 136) |
| **Europe (n=117)** | 71.8% said Yes (84 out of 117)<br>28.2% said No (33 out of 117) |
| **Latin America+Oceania+Africa+Asia (n=106)** | 74.5% said Yes (79 out of 106)<br>25.5% said No (27 out of 106) |

**Table S24. Mentor values a working environment/colleagues with a diverse background?**
On a Likert Scale (1 lowest-5 highest), Data analysis for this table excludes survey respondents with "0" number of mentors.

| Theme | Mentor values colleagues from diverse backgrounds |
|---|---|
| **Total (n=355)** | 2.2% rated 1 (lowest score) (8 out of 355)<br>3.4% rated 2 (12 out of 355)<br>15.8% rated 3 (56 out of 355)<br>29.8% rated 4 (106 out of 355)<br>48.7% rated 5 (highest score) (173 out of 355) |
| **Women Mentee (n=198)** | 3.3% rated 1 (lowest score) (6 out of 198)<br>3.8% rated 2 (5 out of 198)<br>15.8% rated 3 (30 out of 198)<br>26.3% rated 4 (55 out of 198)<br>50.7% rated 5 (highest score) (102 out of 198) |
| **Men Mentee (n=153)** | 1.2% rated 1 (lowest score) (2 out of 153)<br>4.3% rated 2 (7 out of 153)<br>15.9% rated 3 (25 out of 153)<br>35.6% rated 4 (51 out of 153)<br>42.9% rated 5 (highest score) (68 out of 153) |
| **Non-binary or did not disclose gender** | 0% rated 1 (lowest score) (0 out of 4) |

| | |
|---|---|
| **(n=4)** | 0% rated 2 (0 out of 4)<br>25% rated 3 (1 out of 4)<br>0% rated 4 (0 out of 4)<br>75% rated 5 (highest score) (3 out of 4) |

| **Table S25. Mentor established a relationship based on trust with you**<br>On a Likert Scale (1 lowest-5 highest), Data analysis for this table excludes survey respondents with "0" number of mentors. | |
|---|---|
| **Theme** | **Mentor established trusting relationship with Mentee** |
| **Total (n=357)** | 3.2% rated 1 (lowest score) (8 out of 357)<br>4.5% rated 2 (13 out of 357)<br>10% rated 3 (36 out of 357)<br>29.6% rated 4 (109 out of 357)<br>52.8% rated 5 (highest score) (191 out of 357) |
| **Women Mentee(n=196)** | 2.9% rated 1 (lowest score) (3 out of 196)<br>6.3% rated 2 (10 out of 196)<br>9.2% rated 3 (19 out of 196)<br>29.5% rated 4 (60 out of 196)<br>52.2% rated 5 (highest score) (104 out of 196) |
| **Men Mentee(n=156)** | 3% rated 1 (lowest score) (5 out of 156)<br>2.4% rated 2 (3 out of 156)<br>10.2% rated 3 (16 out of 156)<br>30.1% rated 4 (48 out of 156)<br>53.6% rated 5 (highest score) (84 out of 156) |
| **Non-binary or did not disclose gender (n=5)** | 0% rated 1 (lowest score) (0 out of 5)<br>0% rated 2 (0 out of 5)<br>20% rated 3 (1 out of 5)<br>20% rated 4 (1 out of 5)<br>60% rated 5 (highest score) (3 out of 5) |

| **Table S26. Mentor acknowledges your professional contributions.**<br>On a Likert Scale (1 lowest-5 highest), Data analysis for this table excludes survey respondents with "0" number of mentors. | |
|---|---|
| **Theme** | **Mentor acknowledging mentee contributions** |
| **Total (n=359)** | 2.8% rated 1 (lowest score) (10 out of 359)<br>4.7% rated 2 (17 out of 359)<br>11.1% rated 3 (40 out of 359)<br>29.2% rated 4 (105 out of 359)<br>52.1% rated 5 (highest score) (187 out of 359) |
| **Women Mentee (n=198)** | 3% rated 1 (lowest score) (6 out of 198)<br>6.1% rated 2 (12 out of 198) |

| | |
|---|---|
| | 10.6% rated 3 (21 out of 198)<br>29.3% rated 4 (58 out of 198)<br>51% rated 5 (highest score) (101 out of 198) |
| **Men Mentee (n=155)** | 1.9% rated 1 (lowest score) (3 out of 155)<br>3.2% rated 2 (5 out of 155)<br>12.3% rated 3 (19 out of 155)<br>30.3% rated 4 (47 out of 155)<br>52.3% rated 5 (highest score) (81 out of 155) |
| **Non-binary or did not disclose gender (n=6)** | 16.7% rated 1 (lowest score) (1 out of 6)<br>0% rated 2 (0 out of 6)<br>0% rated 3 (0 out of 6)<br>0% rated 4 (0 out of 6)<br>83.3% rated 5 (highest score) (5 out of 6) |

**Table S27. Mentor works effectively with mentee whose personal background is different from their own (age, race, gender, class, region, culture, religion, family composition etc.)**
On a Likert Scale (1 lowest-5 highest), Data analysis for this table excludes survey respondents with "0" number of mentors.

| Theme | Mentor works effectively with mentee of different backgrounds |
|---|---|
| **Total (n=336)** | 3.3% rated 1 (lowest score) (11 out of 336)<br>3.9% rated 2 (13 out of 336)<br>27.4% rated 3 (92 out of 336)<br>30.3% rated 4 (102 out of 336)<br>35.1% rated 5 (highest score) (118 out of 336) |
| **Women Mentee (n=185)** | 4.9% rated 1 (lowest score) (9 out of 185)<br>3.2% rated 2 (6 out of 185)<br>24.3% rated 3 (45 out of 185)<br>31.9% rated 4 (59 out of 185)<br>35.7% rated 5 (highest score) (66 out of 185) |
| **Men Mentee (n=147)** | 0.68% rated 1 (lowest score) (1 out of 147)<br>4.8% rated 2 (7 out of 147)<br>30.6% rated 3 (45 out of 147)<br>29.3% rated 4 (43 out of 147)<br>34.7% rated 5 (highest score) (51 out of 147) |
| **Non-binary or did not disclose gender (n=4)** | 25% rated 1 (lowest score) (1 out of 4)<br>0% rated 2 (0 out of 4)<br>50% rated 3 (2 out of 4)<br>0% rated 4 (0 out of 4)<br>25% rated 5 (highest score) (1 out of 4) |

| Table S28. Mentor provides you with constructive feedback & positive affirmation<br>On a Likert Scale (1 lowest-5 highest), Data analysis for this table excludes survey respondents with "0" number of mentors. | |
|---|---|
| **Theme** | **Mentor provides constructive feedback to Mentee** |
| **Total (n=360)** | 5% rated 1 (lowest score) (14 out of 360)<br>5.7% rated 2 (17 out of 360)<br>9.4% rated 3 (35 out of 360)<br>30.4% rated 4 (114 out of 360)<br>49.5% rated 5 (highest score) (180 out of 360) |
| **Women Mentee (n=199)** | 7.6% rated 1 (lowest score) (12 out of 199)<br>5.2% rated 2 (8 out of 199)<br>8.1% rated 3 (17 out of 199)<br>25.7% rated 4 (54 out of 199)<br>53.3% rated 5 (highest score) (108 out of 199) |
| **Men Mentee (n=156)** | 0.6% rated 1 (lowest score) (1 out of 156)<br>6.6% rated 2 (9 out of 156)<br>11.4% rated 3 (18 out of 156)<br>36.7% rated 4 (59 out of 156)<br>44.6% rated 5 (highest score) (69 out of 156) |
| **Non-binary or did not disclose gender (n=5)** | 33.3% rated 1 (lowest score) (1 out of 5)<br>0% rated 2 (0 out of 5)<br>0% rated 3 (0 out of 5)<br>16.7% rated 4 (1 out of 5)<br>50% rated 5 (highest score) (3 out of 5) |

| Table S29. Mentor ensures a working environment free from discrimination and harassment<br>On a Likert Scale (1 lowest-5 highest), Data analysis for this table excludes survey respondents with "0" number of mentors. | |
|---|---|
| **Theme** | **Mentor ensures safe working environment for Mentee** |
| **Total (n=354)** | 4.8% rated 1 (lowest score) (17 out of 354)<br>5.4% rated 2 (19 out of 354)<br>18.6% rated 3 (66 out of 354)<br>22.3% rated 4 (79 out of 354)<br>48.9% rated 5 (highest score) (173 out of 354) |
| **Women Mentee (n=195)** | 5.1% rated 1 (lowest score) (10 out of 195)<br>5.1% rated 2 (10 out of 195)<br>20% rated 3 (39 out of 195)<br>21.5% rated 4 (42 out of 195)<br>48.2% rated 5 (highest score) (94 out of 195) |
| **Men Mentee (n=154)** | 4.5% rated 1 (lowest score) (7 out of 154)<br>5.8% rated 2 (9 out of 154)<br>16.2% rated 3 (25 out of 154)<br>24% rated 4 (37 out of 154) |

| | |
|---|---|
| | 49.3% rated 5 (highest score) (76 out of 154) |
| **Non-binary or did not disclose gender (n=5)** | 0% rated 1 (lowest score) (0 out of 5)<br>0% rated 2 (0 out of 5)<br>40% rated 3 (2 out of 5)<br>0% rated 4 (0 out of 5)<br>60% rated 5 (highest score) (3 out of 5) |

| **Table S30. Mentor uses active listening (Meaning Identifying and accommodating different communication styles and employing strategies to improve communication with you)** On a Likert Scale (1 lowest-5 highest), Data analysis for this table excludes survey respondents with "0" number of mentors. | |
|---|---|
| **Theme** | **Mentor use of active listening with Mentee** |
| **Total (n=355)** | 3.9% rated 1 (lowest score) (14 out of 355)<br>8.2% rated 2 (29 out of 355)<br>23.4% rated 3 (83 out of 355)<br>36.9% rated 4 (131 out of 355)<br>27.6% rated 5 (highest score) (98 out of 355) |
| **Women Mentee (n=195)** | 4.6% rated 1 (lowest score) (9 out of 195)<br>8.2% rated 2 (16 out of 195)<br>21% rated 3 (41 out of 195)<br>36.4% rated 4 (71 out of 195)<br>29.7% rated 5 (highest score) (58 out of 195) |
| **Men Mentee (n=155)** | 3.2% rated 1 (lowest score) (5 out of 155)<br>8.4% rated 2 (13 out of 155)<br>26.4% rated 3 (41 out of 155)<br>37.4% rated 4 (58 out of 155)<br>24.5% rated 5 (highest score) (38 out of 155) |
| **Non-binary or did not disclose gender (n=5)** | 0% rated 1 (lowest score) (0 out of 5)<br>0% rated 2 (0 out of 5)<br>20% rated 3 (1 out of 5)<br>40% rated 4 (2 out of 5)<br>40% rated 5 (highest score) (2 out of 5) |

| **Table S31. Overall, to what extent do you feel that your current mentor is meeting your expectations?** On a Likert Scale (1 lowest-5 highest), Data analysis for this table excludes survey respondents with "0" number of mentors. | |
|---|---|
| **Theme** | **Mentee overall satisfaction with mentor** |
| **Total (n=362)** | 5% rated 1 (lowest score) (18 out of 362)<br>6.1% rated 2 (22 out of 362)<br>18.7% rated 3 (68 out of 362)<br>41.4% rated 4 (150 out of 362)<br>28.7% rated 5 (highest score) (104 out of 362) |

| | |
|---|---|
| **Women Mentee (n=199)** | 8.6% rated 1 (lowest score) (12 out of 199)<br>6.2% rated 2 (12 out of 199)<br>19.6% rated 3 (39 out of 199)<br>34.4% rated 4 (71 out of 199)<br>31.1% rated 5 (highest score) (65 out of 199) |
| **Men Mentee (n=157)** | 3.8% rated 1 (lowest score) (6 out of 157)<br>5.7% rated 2 (9 out of 157)<br>18.5% rated 3 (29 out of 157)<br>49% rated 4 (77 out of 157)<br>22.9% rated 5 (highest score) (36 out of 157) |
| **Non-binary or did not disclose gender (n=6)** | 0% rated 1 (lowest score) (0 out of 6)<br>16.6% rated 2 (1 out of 6)<br>0% rated 3 (0 out of 6)<br>33.3% rated 4 (2 out of 6)<br>50% rated 5 (highest score) (3 out of 6) |

**Table S32. Mentor takes into account the biases and prejudices she/he brings to your mentor/mentee relationship?** On a Likert Scale (1 lowest-5 highest), Data analysis for this table excludes survey respondents with "0" number of mentors.

| Theme | Mentor taking into account implicit biases |
|---|---|
| **Total (n=348)** | 6% rated 1 (lowest score) (21 out of 348)<br>8.4% rated 2 (29 out of 348)<br>35.6% rated 3 (124 out of 348)<br>28.2% rated 4 (98 out of 348)<br>21.8% rated 5 (highest score) (76 out of 348) |
| **Women Mentee (n=193)** | 7.8% rated 1 (lowest score) (15 out of 193)<br>7.8% rated 2 (15 out of 193)<br>32.1% rated 3 (62 out of 193)<br>28% rated 4 (54 out of 193)<br>24.3% rated 5 (highest score) (47 out of 193) |
| **Men Mentee (n=150)** | 3.4% rated 1 (lowest score) (5 out of 150)<br>9.3% rated 2 (14 out of 150)<br>39.4% rated 3 (59 out of 150)<br>28.6% rated 4 (43 out of 150)<br>19.3% rated 5 (highest score) (29 out of 150) |
| **Non-binary or did not disclose gender (n=5)** | 20% rated 1 (lowest score) (1 out of 5)<br>0% rated 2 (0 out of 5)<br>60% rated 3 (3 out of 5)<br>20% rated 4 (1 out of 5)<br>0% rated 5 (highest score) (0 out of 5) |

| **Table S33. Mentor helps you develop strategies to meet career goals.** On a Likert Scale (1 lowest-5 highest), Data analysis for this table excludes survey respondents with "0" number of mentors. | |
|---|---|
| **Theme** | **Mentor meeting mentee career goals** |
| **Total (n=360)** | 7.8% rated 1 (lowest score) (28 out of 360)<br>7.5% rated 2 (27 out of 360)<br>17.5% rated 3 (63 out of 360)<br>30.5% rated 4 (110 out of 360)<br>36.7% rated 5 (highest score) (132 out of 360) |
| **Women Mentee (n=199)** | 9.1% rated 1 (lowest score) (18 out of 199)<br>8% rated 2 (16 out of 199)<br>18.1% rated 3 (36 out of 199)<br>28.1% rated 4 (56 out of 199)<br>36.7% rated 5 (highest score) (73 out of 199) |
| **Men Mentee (n=155)** | 5.8% rated 1 (lowest score) (9 out of 155)<br>7.1% rated 2 (11 out of 155)<br>17.4% rated 3 (27 out of 155)<br>32.9% rated 4 (51 out of 155)<br>36.8% rated 5 (highest score) (57 out of 155) |
| **Non-binary or did not disclose gender (n=6)** | 16.7% rated 1 (lowest score) (1 out of 6)<br>0% rated 2 (0 out of 6)<br>0% rated 3 (0 out of 6)<br>50% rated 4 (3 out of 6)<br>33.3% rated 5 (highest score) (2 out of6) |

| **Table S34. Mentor helps you acquire resources (e.g., grants, instrumentation, collaborations etc.),** On a Likert Scale (1 lowest-5 highest), Data analysis for this table excludes survey respondents with "0" number of mentors. | |
|---|---|
| **Theme** | **Mentor's assistance in acquiring resources** |
| **Total (n=361)** | 7.5% rated 1 (lowest score) (27 out of 361)<br>10.8% rated 2 (39 out of 361)<br>13% rated 3 (47 out of 361)<br>27.1% rated 4 (98 out of 361)<br>41.6% rated 5 (highest score) (150 out of 361) |
| **Women Mentee (n=199)** | 10% rated 1 (lowest score) (20 out of 199)<br>8.5% rated 2 (17 out of 199)<br>12.6% rated 3 (25 out of 199)<br>25.6% rated 4 (51 out of 199)<br>43.2% rated 5 (highest score) (86 out of 199) |
| **Men Mentee (n=156)** | 4.5% rated 1 (lowest score) (7 out of 156)<br>14.1% rated 2 (22 out of 156)<br>13.5% rated 3 (21 out of 156)<br>28.2% rated 4 (44 out of 156)<br>39.7% rated 5 (highest score) (62 out of 156) |

| | |
|---|---|
| **Non-binary or did not disclose gender (n=6)** | 0% rated 1 (lowest score) (0 out of 6)<br>0% rated 2 (0 out of 6)<br>16.7% rated 3 (1 out of 6)<br>50% rated 4 (3 out of 6)<br>33.3% rated 5 (highest score) (2 out of 6) |

**Table S35. Mentor helps you develop strategies to better mentor your own mentees (undergraduate/graduate/postdoctoral trainees)?** On a Likert Scale (1 lowest-5 highest), Data analysis for this table excludes survey respondents with "0" number of mentors.

| Theme | Mentor helps mentee with strategies to mentor their own lab more effectively |
|---|---|
| **Total (n=351)** | 11.4% rated 1 (lowest score) (40 out of 351)<br>13.1% rated 2 (46 out of 351)<br>29.1% rated 3 (102 out of 351)<br>27.6% rated 4 (97 out of 351)<br>18.8% rated 5 (highest score) (66 out of 351) |
| **Women Mentee (n=192)** | 13.5% rated 1 (lowest score) (26 out of 192)<br>14.6% rated 2 (28 out of 192)<br>27.1% rated 3 (52 out of 192)<br>26.6% rated 4 (51 out of 192)<br>18.2% rated 5 (highest score) (35 out of 192) |
| **Men Mentee (n=153)** | 8.5% rated 1 (lowest score) (13 out of 153)<br>11.8% rated 2 (18 out of 153)<br>31.4% rated 3 (48 out of 153)<br>28.8% rated 4 (44 out of 153)<br>19.5% rated 5 (highest score) (30 out of 153) |
| **Non-binary or did not disclose gender (n=6)** | 16.7% rated 1 (lowest score) (1 out of 6)<br>0% rated 2 (0 out of 6)<br>33.3% rated 3 (2 out of 6)<br>33.3% rated 4 (2 out of 6)<br>16.7% rated 5 (highest score) (1 out of 6) |

**Table S36. Mentor introduce you to speakers and other professors at meetings and seminars and scientific society gatherings?** On a Likert Scale (1 lowest-5 highest) by gender, Data analysis for this table excludes survey respondents with "0" number of mentors.

| Theme | Mentor promoting mentee by expanding mentee professional network |
|---|---|
| **Total (n=354)** | 13.8% rated 1 (lowest score) (49 out of 354)<br>13% rated 2 (46 out of 354)<br>22.0% rated 3 (78 out of 354)<br>24.9% rated 4 (88 out of 354)<br>26.3% rated 5 (highest score) (93 out of 354) |

| | |
|---|---|
| **Women Mentee (n=194)** | 16.5% rated 1 (lowest score) (32 out of 194)<br>17% rated 2 (33 out of 194)<br>17% rated 3 (33 out of 194)<br>23.2% rated 4 (45 out of 194)<br>26.3% rated 5 (highest score) (51 out of 194) |
| **Men Mentee (n=154)** | 10.4% rated 1 (lowest score) (16 out of 154)<br>8.3% rated 2 (13 out of 154)<br>27.9% rated 3 (43 out of 154)<br>27.3% rated 4 (42 out of 154)<br>26.0% rated 5 (highest score) (40 out of 154) |
| **Non-binary or did not disclose gender (n=6)** | 16.7% rated 1 (lowest score) (1 out of 6)<br>0% rated 2 (0 out of 6)<br>33.3% rated 3 (2 out of 6)<br>16.7% rated 4 (1 out of 6)<br>33.3% rated 5 (highest score) (2 out of 6) |
| **Did not respond to this survey Question** | 17.7% (81 out of 457) |

| **Table S37. Mentor suggests ways to you to balance work with your personal life**<br>On a Likert Scale (1 lowest-5 highest), Data analysis for this table excludes survey respondents with "0" number of mentors. | |
|---|---|
| **Theme** | **Mentor suggests ways mentee can balance work-life** |
| **Total (n=353)** | 15% rated 1 (lowest score) (53 out of 353)<br>18.4% rated 2 (65 out of 353)<br>34% rated 3 (120 out of 353)<br>14.4% rated 4 (51 out of 353)<br>18.1% rated 5 (highest score) (64 out of 353) |
| **Women Mentee (n=196)** | 16.3% rated 1 (lowest score) (32 out of 196)<br>19.9% rated 2 (39 out of 196)<br>31.1% rated 3 (61 out of 196)<br>12.2% rated 4 (24 out of 196)<br>20.4% rated 5 (highest score) (40 out of 196) |
| **Men Mentee (n=152)** | 13.1% rated 1 (lowest score) (20 out of 152)<br>16.5% rated 2 (25 out of 152)<br>37.5% rated 3 (57 out of 152)<br>17.8% rated 4 (27 out of 152)<br>15.1% rated 5 (highest score) (23 out of 152) |
| **Non-binary or did not disclose gender (n=5)** | 20% rated 1 (lowest score) (1 out of 5)<br>20% rated 2 (1 out of 5)<br>40% rated 3 (2 out of 5)<br>0% rated 4 (0 out of 5)<br>20% rated 5 (highest score) (1 out of 5) |

| Table S38. Please rate your feeling of happiness and satisfaction with your current research and position as an independent investigator. On a Likert Scale (1 lowest-5 highest), Data analysis for this table includes survey respondents with "0" number of mentors. | |
|---|---|
| **Theme** | **Rating of Mentee satisfaction with research program** |
| **Total (n=370)** | 5.9% rated 1 (lowest score) (22 out of 370)<br>9.7% rated 2 (36 out of 370)<br>18.9% rated 3 (70 out of 370)<br>43.8% rated 4 (162 out of 370)<br>21.6% rated 5 (highest score) (80 out of 370) |
| **Women Mentee (n=195)** | 7.7% rated 1 (lowest score) (15 out of 195)<br>12.3% rated 2 (24 out of 195)<br>20.5% rated 3 (40 out of 195)<br>40.5% rated 4 (79 out of 195)<br>19% rated 5 (highest score) (37 out of 195) |
| **Men Mentee (n=170)** | 3.5% rated 1 (lowest score) (6 out of 170)<br>6.5% rated 2 (11 out of 170)<br>17.6% rated 3 (30 out of 170)<br>48.2% rated 4 (82 out of 170)<br>24.1% rated 5 (highest score) (41 out of 170) |
| **Non-binary or did not disclose gender (n=5)** | 20% rated 1 (lowest score) (1 out of 5)<br>20% rated 2 (1 out of 5)<br>0% rated 3 (0 out of 5)<br>20% rated 4 (1 out of 5)<br>40% rated 5 (highest score) (2 out of 5) |
| **Did not respond to this survey Question** | 19% (87 out of 457) |

| Table S38a. Please rate your feeling of happiness and satisfaction with your current research and position as an independent investigator. On a Likert Scale (1 lowest-5 highest), Data analysis for this table includes survey respondents with "0" number of mentors. | |
|---|---|
| **Theme** | **Rating of Mentee satisfaction with research program** |
| **North America (n=136)** | 5.1% rated 1 (lowest score) (7 out of 136)<br>7.4% rated 2 (10 out of 136)<br>13.2% rated 3 (18 out of 136)<br>45.6% rated 4 (62 out of 136)<br>28.7% rated 5 (highest score) (39 out of 136) |
| **Europe (n=125)** | 7.2% rated 1 (lowest score) (9 out of 125)<br>14.4% rated 2 (18 out of 125)<br>20% rated 3 (25 out of 125)<br>40.8% rated 4 (51 out of 125)<br>17.6% rated 5 (highest score) (22 out of 125) |
| **Latin America+Oceania+Africa+Asia (n=109)** | 6.4% rated 1 (lowest score) (7 out of 109)<br>10.1% rated 2 (11 out of 109) |

| | |
|---|---|
| | 24.8% rated 3 (27 out of 109)<br>44% rated 4 (48 out of 109)<br>14.7% rated 5 (highest score) (16 out of 109) |
| **Did not respond to this survey Question** | 19.3% (87 out of 450) |

| **Table S38b. Please rate your feeling of happiness and satisfaction with your current research and position as an independent investigator.** On a Likert Scale (1 lowest-5 highest), Data analysis for this table includes survey respondents with "0" number of mentors. | |
|---|---|
| **Theme** | **Rating of Mentee satisfaction with research program** |
| **Mentor was assigned to faculty mentee (n=104)** | 4.8% rated 1 (lowest score) (5 out of 104)<br>8.7% rated 2 (9 out of 104)<br>20.2% rated 3 (21 out of 104)<br>50% rated 4 (52 out of 104)<br>16.3% rated 5 (highest score) (17 out of 104) |
| **Faculty mentee chose mentor voluntarily (n=249)** | 4.2% rated 1 (lowest score) (10 out of 249)<br>8.0% rated 2 (20 out of 249)<br>19.3% rated 3 (48 out of 249)<br>43% rated 4 (107 out of 249)<br>25.7% rated 5 (highest score) (64 out of 249) |
| **Did not respond to this survey Question** | 10.2% (39 out of 383) |

| **Table S39. Please rate your optimism about your future career as an independent investigator.** On a Likert Scale (1 lowest-5 highest), analysis by gender, Data analysis for this table includes survey respondents with "0" number of mentors. | |
|---|---|
| **Theme** | **Rating of Mentee optimism about the future** |
| **Total (n=371)** | 4% rated 1 (lowest score) (15 out of 371)<br>12.7% rated 2 (47 out of 371)<br>17.5% rated 3 (65 out of 371)<br>42.9% rated 4 (159 out of 371)<br>22.9% rated 5 (highest score) (85 out of 371) |
| **Women Mentee (n=196)** | 5.1% rated 1 (lowest score) (10 out of 196)<br>14.8% rated 2 (29 out of 196)<br>22.4% rated 3 (44 out of 196)<br>38.8% rated 4 (76 out of 196)<br>18.9% rated 5 (highest score) (37 out of 196) |
| **Men Mentee (n=170)** | 2.4% rated 1 (lowest score) (4 out of 170)<br>10.6% rated 2 (18 out of 170)<br>11.2% rated 3 (19 out of 170)<br>48.8% rated 4 (83 out of 170)<br>27.1% rated 5 (highest score) (46 out of 170) |

| | |
|---|---|
| **Non-binary or did not disclose gender (n=5)** | 20% rated 1 (lowest score) (1 out of 5)<br>0% rated 2 (0 out of 5)<br>40% rated 3 (2 out of 5)<br>0% rated 4 (0 out of 5)<br>40% rated 5 (highest score) (2 out of 5) |
| **Did not respond to this survey Question** | 18.8% (86 out of 457) |

| **Table S39a. Please rate your optimism about the future as an independent investigator.** On a Likert Scale (1 lowest-5 highest), analysis by continent, Data analysis for this table includes survey respondents with "0" number of mentors. | |
|---|---|
| **Theme** | **Rating of Mentee optimism about the future** |
| **North America (n=138)** | 2.2% rated 1 (lowest score) (3 out of 138)<br>5.8% rated 2 (8 out of 138)<br>15.9% rated 3 (22 out of 138)<br>44.2% rated 4 (61 out of 138)<br>31.9% rated 5 (highest score) (44 out of 138) |
| **Europe (n=121)** | 4.1% rated 1 (lowest score) (5 out of 121)<br>19.8% rated 2 (24 out of 121)<br>19.8% rated 3 (24 out of 121)<br>38.8% rated 4 (47 out of 121)<br>17.4% rated 5 (highest score) (21 out of 121) |
| **Latin America+Africa+Asia+Oceania (n=107)** | 4.7% rated 1 (lowest score) (5 out of 107)<br>13.1% rated 2 (14 out of 107)<br>18.7% rated 3 (20 out of 107)<br>45.8% rated 4 (49 out of 107)<br>17.7% rated 5 (highest score) (19 out of 107) |
| **Did not respond to this survey Question** | 20.2% (91 out of 450) |

| **Table S39b. Please rate your optimism about the future as an independent investigator.** On a Likert Scale (1 lowest-5 highest), Data analysis for this table includes survey respondents with "0" number of mentors. | |
|---|---|
| **Theme** | **Rating of Mentee satisfaction with research program** |
| **Mentor was assigned to faculty mentee (n=104)** | 3.8% rated 1 (lowest score) (4 out of 104)<br>13.5% rated 2 (14 out of 104)<br>17.3% rated 3 (18 out of 104)<br>44.2% rated 4 (46 out of 104)<br>21.2% rated 5 (highest score) (22 out of 104) |
| **Faculty mentee chose mentor voluntarily (n=248)** | 2% rated 1 (lowest score) (5 out of 248)<br>10.5% rated 2 (26 out of 248)<br>17.7% rated 3 (44 out of 248)<br>45.2% rated 4 (112 out of 248) |

| | |
|---|---|
| | 24.6% rated 5 (highest score) (61 out of 248) |
| **Did not respond to this survey Question (n=41)** | 10.7% (41 out of 383) |

**Table S40a. corresponding to Figure S3a**
**Ordinary one-way ANOVA results, multiple comparisons, mentorship quality (poor-to-excellent rating of quality of match between faculty mentee and their mentor) for same versus different mentor-mentee gender.** Data analysis for this table excludes survey respondents with "0" number of mentors.

| | |
|---|---|
| **Number of families** | 1 |
| **Number of comparisons per family** | 6 |
| **Alpha** | 0.05 |

| **Holm-Šídák's multiple comparisons test** | Mean Diff. | 95.00% CI of diff. | Below threshold? | Summary | Adjusted P Value | |
|---|---|---|---|---|---|---|
| **Assigned - same vs. Assigned - diff** | -0.1532 | -0.6311 to 0.3247 | No | ns | 0.9519 | A-B |
| **Assigned - same vs. Voluntary - same** | -0.5724 | -1.011 to -0.1334 | Yes | ** | 0.0038 | A-C |
| **Assigned - same vs. Voluntary - different** | -0.4113 | -0.8937 to 0.07106 | No | ns | 0.1393 | A-D |
| **Assigned - diff vs. Voluntary - different** | -0.2581 | -0.6891 to 0.1729 | No | ns | 0.5164 | B-D |
| **Assigned - diff vs. Voluntary - same** | -0.4192 | -0.8010 to -0.03738 | Yes | * | 0.0232 | B-C |
| **Voluntary - same vs. Voluntary - different** | 0.1611 | -0.2264 to 0.5485 | No | ns | 0.8513 | C-D |

| **Test details** | Mean 1 | Mean 2 | Mean Diff. | SE of diff. | n1 | n2 | t | DF |
|---|---|---|---|---|---|---|---|---|
| **Assigned - same vs. Assigned - diff** | 3.517 | 3.67 | -0.1532 | 0.1807 | 58 | 88 | 0.8481 | 371 |
| **Assigned - same vs. Voluntary - same** | 3.517 | 4.09 | -0.5724 | 0.166 | 58 | 145 | 3.449 | 371 |
| **Assigned - same vs. Voluntary - different** | 3.517 | 3.929 | -0.4113 | 0.1824 | 58 | 84 | 2.256 | 371 |
| **Assigned - diff vs. Voluntary - different** | 3.67 | 3.929 | -0.2581 | 0.1629 | 88 | 84 | 1.584 | 371 |

| | | | | | | | | |
|---|---|---|---|---|---|---|---|---|
| **Assigned - diff vs. Voluntary - same** | 3.67 | 4.09 | -0.4192 | 0.1443 | 88 | 145 | 2.904 | 371 |
| **Voluntary - same vs. Voluntary - different** | 4.09 | 3.929 | 0.1611 | 0.1465 | 145 | 84 | 1.1 | 371 |

| **Table S40b. corresponding to Figure S3b**<br>**Ordinary one-way ANOVA results, multiple comparisons, mentorship quality (poor-to-excellent rating of quality of match between faculty mentee and their mentor) for Men versus Women, Assigned versus Voluntarily selected mentor(s).**<br>Data analysis for this table excludes survey respondents with "0" number of mentors. | | | | | | | | |
|---|---|---|---|---|---|---|---|---|
| **Number of families** | 1 | | | | | | | |
| **Number of comparisons per family** | 6 | | | | | | | |
| **Alpha** | 0.05 | | | | | | | |
| **Tukey's multiple comparisons test** | Mean Diff. | 95.00% CI of diff. | Below threshold? | Summary | Adjusted P Value | | | |
| **M - assigned vs. M - chose** | -0.5621 | -1.040 to -0.08458 | Yes | * | 0.0135 | A-B | | |
| **M - assigned vs. F - assigned** | 0.1573 | -0.3742 to 0.6888 | No | ns | 0.8708 | A-C | | |
| **M - assigned vs. F - chose** | -0.503 | -0.9718 to -0.03413 | Yes | * | 0.03 | A-D | | |
| **M - chose vs. F - assigned** | 0.7193 | 0.2991 to 1.140 | Yes | **** | <0.0001 | B-C | | |
| **M - chose vs. F - chose** | 0.05909 | -0.2785 to 0.3967 | No | ns | 0.9693 | B-D | | |
| **F - assigned vs. F - chose** | -0.6602 | -1.071 to -0.2498 | Yes | *** | 0.0002 | C-D | | |
| **Test details** | Mean 1 | Mean 2 | Mean Diff. | SE of diff. | n1 | n2 | t | DF |
| **M - assigned vs. M - chose** | 3.511 | 4.073 | -0.5621 | 0.185 | 45 | 123 | 4.296 | 371 |
| **M - assigned vs. F - assigned** | 3.511 | 3.354 | 0.1573 | 0.2059 | 45 | 65 | 1.08 | 371 |
| **M - assigned vs. F - chose** | 3.511 | 4.014 | -0.503 | 0.1817 | 45 | 142 | 3.915 | 371 |
| **M - chose vs. F - assigned** | 4.073 | 3.354 | 0.7193 | 0.1629 | 123 | 65 | 6.247 | 371 |
| **M - chose vs. F - chose** | 4.073 | 4.014 | 0.05909 | 0.1308 | 123 | 142 | 0.6388 | 371 |
| **F - assigned vs. F - chose** | 3.354 | 4.014 | -0.6602 | 0.159 | 65 | 142 | 5.871 | 371 |

| Table S40c. corresponding to Figure S3c<br>**Ordinary one-way ANOVA results, mentorship quality (poor-to-excellent rating of quality of match between faculty mentee and their mentor) by number of mentors.**<br>Data analysis for this table excludes survey respondents with "0" number of mentors. | | | | | | | | |
|---|---|---|---|---|---|---|---|---|
| **Number of families** | 1 | | | | | | | |
| **Number of comparisons per family** | 15 | | | | | | | |
| **Alpha** | 0.05 | | | | | | | |
| **Tukey's multiple comparisons test** | **Mean Diff.** | **95.00% CI of diff.** | **Below threshold ?** | **Summary** | **Adjusted P Value** | | | |
| **1 vs. 2** | -0.1829 | -0.5763 to 0.2106 | No | ns | 0.7672 | A-B | | |
| **1 vs. 3** | -0.2427 | -0.6996 to 0.2142 | No | ns | 0.6502 | A-C | | |
| **1 vs. 4** | -0.4488 | -1.147 to 0.2494 | No | ns | 0.4402 | A-D | | |
| **1 vs. 5** | -0.525 | -1.333 to 0.2834 | No | ns | 0.4282 | A-E | | |
| **1 vs. 5+** | -0.5083 | -1.096 to 0.07896 | No | ns | 0.1328 | A-F | | |
| **2 vs. 3** | -0.05985 | -0.5271 to 0.4074 | No | ns | 0.9991 | B-C | | |
| **2 vs. 4** | -0.2659 | -0.9710 to 0.4391 | No | ns | 0.8888 | B-D | | |
| **2 vs. 5** | -0.3421 | -1.156 to 0.4722 | No | ns | 0.8348 | B-E | | |
| **2 vs. 5+** | -0.3255 | -0.9209 to 0.2699 | No | ns | 0.6213 | B-F | | |
| **3 vs. 4** | -0.2061 | -0.9484 to 0.5362 | No | ns | 0.9682 | C-D | | |
| **3 vs. 5** | -0.2823 | -1.129 to 0.5645 | No | ns | 0.9315 | C-E | | |
| **3 vs. 5+** | -0.2656 | -0.9047 to 0.3735 | No | ns | 0.8411 | C-F | | |
| **4 vs. 5** | -0.07619 | -1.074 to 0.9217 | No | ns | >0.9999 | D-E | | |
| **4 vs. 5+** | -0.05952 | -0.8885 to 0.7695 | No | ns | >0.9999 | D-F | | |
| **5 vs. 5+** | 0.01667 | -0.9070 to 0.9404 | No | ns | >0.9999 | E-F | | |
| **Test details** | **Mean 1** | **Mean 2** | **Mean Diff.** | **SE of diff.** | **n1** | **n2** | **t** | **DF** |
| **1 vs. 2** | 3.742 | 3.925 | -0.1829 | 0.1373 | 120 | 106 | 1.883 | 352 |
| **1 vs. 3** | 3.742 | 3.984 | -0.2427 | 0.1595 | 120 | 64 | 2.153 | 352 |
| **1 vs. 4** | 3.742 | 4.19 | -0.4488 | 0.2437 | 120 | 21 | 2.605 | 352 |
| **1 vs. 5** | 3.742 | 4.267 | -0.525 | 0.2821 | 120 | 15 | 2.632 | 352 |
| **1 vs. 5+** | 3.742 | 4.25 | -0.5083 | 0.205 | 120 | 32 | 3.508 | 352 |
| **2 vs. 3** | 3.925 | 3.984 | -0.05985 | 0.1631 | 106 | 64 | 0.519 | 352 |
| **2 vs. 4** | 3.925 | 4.19 | -0.2659 | 0.2461 | 106 | 21 | 1.529 | 352 |

| | | | | | | | | |
|---|---|---|---|---|---|---|---|---|
| **2 vs. 5** | 3.925 | 4.267 | -0.3421 | 0.2842 | 106 | 15 | 1.703 | 352 |
| **2 vs. 5+** | 3.925 | 4.25 | -0.3255 | 0.2078 | 106 | 32 | 2.215 | 352 |
| **3 vs. 4** | 3.984 | 4.19 | -0.2061 | 0.2591 | 64 | 21 | 1.125 | 352 |
| **3 vs. 5** | 3.984 | 4.267 | -0.2823 | 0.2955 | 64 | 15 | 1.351 | 352 |
| **3 vs. 5+** | 3.984 | 4.25 | -0.2656 | 0.223 | 64 | 32 | 1.684 | 352 |
| **4 vs. 5** | 4.19 | 4.267 | -0.07619 | 0.3483 | 21 | 15 | 0.3094 | 352 |
| **4 vs. 5+** | 4.19 | 4.25 | -0.05952 | 0.2893 | 21 | 32 | 0.291 | 352 |
| **5 vs. 5+** | 4.267 | 4.25 | 0.01667 | 0.3224 | 15 | 32 | 0.07312 | 352 |

**Table S40d. corresponding to Figure S3d**
**Ordinary one-way ANOVA results, Multiple Comparisons, Number of mentors reported by mentees in North America, Europe, or other continents.**
Data analysis for this table includes survey respondents with "0" number of mentors.

| | | | | | | | | |
|---|---|---|---|---|---|---|---|---|
| **Number of families** | 1 | | | | | | | |
| **Number of comparisons per family** | 3 | | | | | | | |
| **Alpha** | 0.05 | | | | | | | |
| **Tukey's multiple comparisons test** | **Mean Diff.** | **95.00% CI of diff.** | **Below threshold?** | **Summary** | **Adjusted P Value** | | | |
| **North America vs. Europe** | 1.183 | 0.7656 to 1.601 | Yes | **** | <0.0001 | A-B | | |
| **North America vs. Column C** | 1.093 | 0.6466 to 1.540 | Yes | **** | <0.0001 | A-C | | |
| **Europe vs. Column C** | -0.08988 | -0.5589 to 0.3791 | No | ns | 0.894 | B-C | | |
| **Test details** | **Mean 1** | **Mean 2** | **Mean Diff.** | **SE of diff.** | **n1** | **n2** | **q** | **DF** |
| **North America vs. Europe** | 3.115 | 1.932 | 1.183 | 0.1775 | 148 | 117 | 9.43 | 355 |
| **North America vs. Column C** | 3.115 | 2.022 | 1.093 | 0.1898 | 148 | 93 | 8.146 | 355 |
| **Europe vs. Column C** | 1.932 | 2.022 | -0.08988 | 0.1993 | 117 | 93 | 0.6379 | 355 |

**Table S40e. corresponding to Figure S3e**
**Ordinary one-way ANOVA results, Multiple Comparisons, mentorship quality (poor-to-excellent rating of quality of match between faculty mentee and their mentor) by mentees in North America, Europe, or other continents.** Data analysis for this table excludes survey respondents with "0" number of mentors.

| | |
|---|---|
| **Number of families** | 1 |
| **Number of comparisons per family** | 3 |
| **Alpha** | 0.05 |

| | | | | | | | | |
|---|---|---|---|---|---|---|---|---|
| **Tukey's multiple comparisons test** | **Mean Diff.** | **95.00% CI of diff.** | **Below threshold?** | **Summary** | **Adjusted P Value** | | | |
| **North America vs. Europe** | 0.125 | -0.1775 to 0.4276 | No | ns | 0.5946 | A-B | | |
| **North America vs. Column C** | 0.02274 | -0.3009 to 0.3463 | No | ns | 0.985 | A-C | | |
| **Europe vs. Column C** | -0.1023 | -0.4420 to 0.2375 | No | ns | 0.7586 | B-C | | |
| **Test details** | **Mean 1** | **Mean 2** | **Mean Diff.** | **SE of diff.** | **n1** | **n2** | **q** | **DF** |
| **North America vs. Europe** | 3.98 | 3.855 | 0.125 | 0.1285 | 148 | 117 | 1.376 | 355 |
| **North America vs. Column C** | 3.98 | 3.957 | 0.02274 | 0.1375 | 148 | 93 | 0.2339 | 355 |
| **Europe vs. Column C** | 3.855 | 3.957 | -0.1023 | 0.1444 | 117 | 93 | 1.002 | 355 |

| **Table S41a. corresponding to Figure S4a**<br>**Two-tailed Mann-Whitney test results**, **Meeting frequency by mentorship initiation mode: Assigned mentors (column A) versus Voluntarily selected mentors (column B).** Data analysis for this table excludes survey respondents with "0" number of mentors. | |
|---|---|
| **Mann-Whitney test** | |
| P value | 0.9213 |
| Exact or approximate P value? | Exact |
| P value summary | ns |
| Significantly different (P < 0.05)? | No |
| One- or two-tailed P value? | Two-tailed |
| Sum of ranks in column A, B | 4100, 10607 |
| Mann-Whitney U | 2924 |
| Difference between medians | |
| Median of column A | 3.000, n=48 |
| Median of column B | 3.000, n=123 |
| Difference: Actual | 0 |
| Difference: Hodges-Lehmann | 0 |

| **Table S41b. corresponding to Figure S4b**<br>**Two-tailed Mann-Whitney test results**, **Constructiveness of mentorship interactions by mentorship initiation mode: Assigned mentors (column A) versus Voluntarily selected mentors (column B).** Data analysis for this table excludes survey respondents with "0" number of mentors. | |
|---|---|
| **Mann-Whitney test** | |
| P value | 0.004 |
| Exact or approximate P value? | Approximate |
| P value summary | ** |
| Significantly different (P < 0.05)? | Yes |
| One- or two-tailed P value? | Two-tailed |
| Sum of ranks in column A, B | 17464, 48966 |
| Mann-Whitney U | 11793 |

| | |
|---|---|
| Difference between medians | |
| Median of column A | 1.000, n=106 |
| Median of column B | 1.000, n=258 |
| Difference: Actual | 0 |
| Difference: Hodges-Lehmann | 0 |

| **Table S41c. corresponding to Figure S4c** **Two-tailed Mann-Whitney test results**, **Perceived quality of mentorship match (poor-to-excellent) by mentorship initiation mode: Assigned mentors (column A) versus Voluntarily selected mentors (column B).** Data analysis for this table excludes survey respondents with "0" number of mentors. | |
|---|---|
| **Mann-Whitney test** | |
| P value | 0.0001 |
| Exact or approximate P value? | Approximate |
| P value summary | *** |
| Significantly different (P < 0.05)? | Yes |
| One- or two-tailed P value? | Two-tailed |
| Sum of ranks in column A, B | 16005, 50425 |
| Mann-Whitney U | 10334 |
| Difference between medians | |
| Median of column A | 4.000, n=106 |
| Median of column B | 4.000, n=258 |
| Difference: Actual | 0 |
| Difference: Hodges-Lehmann | 0 |

| **Table S41d. corresponding to Figure S4d** **Two-tailed Mann-Whitney test results**, **Number of mentors by faculty mentee gender: Women (column A) versus Men (column B).** Data analysis for this table excludes survey respondents with "0" number of mentors. | |
|---|---|
| **Mann-Whitney test** | |
| P value | 0.0687 |
| Exact or approximate P value? | Approximate |
| P value summary | ns |
| Significantly different (P < 0.05)? | No |
| One- or two-tailed P value? | Two-tailed |
| Sum of ranks in column A, B | 56184, 43944 |
| Mann-Whitney U | 22416 |
| Difference between medians | |
| Median of column A | 2.000, n=240 |
| Median of column B | 1.000, n=207 |
| Difference: Actual | -1 |
| Difference: Hodges-Lehmann | 0 |

| **Table S41e. corresponding to Figure S4e** **Two-tailed Mann-Whitney test results**, **Mentee-mentor meeting frequency by faculty mentee gender: Women (column A) versus Men (column B).** Data analysis for this table excludes survey respondents with "0" number of mentors. | |
|---|---|
| **Mann-Whitney test** | |
| P value | 0.4505 |
| Exact or approximate P value? | Exact |
| P value summary | ns |

| | |
|---|---|
| Significantly different (P < 0.05)? | No |
| One- or two-tailed P value? | Two-tailed |
| Sum of ranks in column A, B | 7119, 7246 |
| Mann-Whitney U | 3330 |
| Difference between medians | |
| Median of column A | 3.000, n=81 |
| Median of column B | 3.000, n=88 |
| Difference: Actual | 0 |
| Difference: Hodges-Lehmann | 0 |

| **Table S41f. corresponding to Figure S4f**<br>**Two-tailed Mann-Whitney test results, Constructiveness of mentorship interactions by faculty mentee gender: Women (column A) versus Men (column B).**<br>Data analysis for this table excludes survey respondents with "0" number of mentors. | |
|---|---|
| **Mann-Whitney test** | |
| P value | 0.1303 |
| Exact or approximate P value? | Approximate |
| P value summary | ns |
| Significantly different (P < 0.05)? | No |
| One- or two-tailed P value? | Two-tailed |
| Sum of ranks in column A, B | 34754, 29866 |
| Mann-Whitney U | 14854 |
| Difference between medians | |
| Median of column A | 1.000, n=199 |
| Median of column B | 1.000, n=160 |
| Difference: Actual | 0 |
| Difference: Hodges-Lehmann | 0 |

| **Table S41g. corresponding to Figure S4g**<br>**Two-tailed Mann-Whitney test results, Quality of mentor-mentee match (poor-to-excellent) by faculty mentee gender: Women (column A) versus Men (column B).**<br>Data analysis for this table excludes survey respondents with "0" number of mentors. | |
|---|---|
| **Mann-Whitney test** | |
| P value | 0.7926 |
| Exact or approximate P value? | Approximate |
| P value summary | ns |
| Significantly different (P < 0.05)? | No |
| One- or two-tailed P value? | Two-tailed |
| Sum of ranks in column A, B | 35478, 28784 |
| Mann-Whitney U | 15578 |
| Difference between medians | |
| Median of column A | 4.000, n=199 |
| Median of column B | 4.000, n=159 |
| Difference: Actual | 0 |
| Difference: Hodges-Lehmann | 0 |

| Table S42a. corresponding to Figure S5a<br>**Two-tailed Mann-Whitney test results, Mentor introduces faculty mentee to colleagues at events? by mentee gender: Women (Column A) versus Men (Column B) mentee.**<br>Data analysis for this table excludes survey respondents with "0" number of mentors. | |
|---|---|
| **Mann-Whitney Test** | |
| P value | 0.154 |
| Exact or approximate P value? | Approximate |
| P value summary | ns |
| Significantly different (P < 0.05)? | No |
| One- or two-tailed P value? | Two-tailed |
| Sum of ranks in column A, B | 32557, 28170 |
| Mann-Whitney U | 13642 |
| **Difference between medians** | |
| Median of column A | 3.000, n=194 |
| Median of column B | 4.000, n=154 |
| Difference: Actual | 1 |
| Difference: Hodges-Lehmann | 0 |

| Table S42b. corresponding to Figure S5b<br>**Two-tailed Mann-Whitney test results, Mentor assists faculty mentee with strategies to mentor their own lab? by gender: Women (column A) versus Men (column B) mentee.**<br>Data analysis for this table excludes survey respondents with "0" number of mentors. | |
|---|---|
| **Mann-Whitney Test** | |
| P value | 0.2353 |
| Exact or approximate P value? | Approximate |
| P value summary | ns |
| Significantly different (P < 0.05)? | No |
| One- or two-tailed P value? | Two-tailed |
| Sum of ranks in column A, B | 32154, 27531 |
| Mann-Whitney U | 13626 |
| **Difference between medians** | |
| Median of column A | 3.000, n=192 |
| Median of column B | 3.000, n=153 |
| Difference: Actual | 0 |
| Difference: Hodges-Lehmann | 0 |

| Table S42c. corresponding to Figure S5c<br>**Two-tailed Mann-Whitney test results, Mentor suggests strategies on balancing work-life? by gender: Women (column A) versus Men (column B) mentee.** Data analysis for this table excludes survey respondents with "0" number of mentors. | |
|---|---|
| **Mann-Whitney Test** | |
| P value | 0.7474 |
| Exact or approximate P value? | Approximate |
| P value summary | ns |
| Significantly different (P < 0.05)? | No |
| One- or two-tailed P value? | Two-tailed |
| Sum of ranks in column A, B | 33911, 26816 |
| Mann-Whitney U | 14605 |
| **Difference between medians** | |
| Median of column A | 3.000, n=196 |
| Median of column B | 3.000, n=152 |

| | |
|---|---|
| Difference: Actual | 0 |
| Difference: Hodges-Lehmann | 0 |

**Table S43a. corresponding to Figure S6a**
**Turkey's multiple comparison test results. Difference in mentees in in North America, Europe, or other continents keeping in touch with former mentor(s).**
Data analysis for this table includes survey respondents with "0" number of mentors.

| | | | | | | | | |
|---|---|---|---|---|---|---|---|---|
| Number of families | 1 | | | | | | | |
| Number of comparisons per family | 3 | | | | | | | |
| Alpha | 0.05 | | | | | | | |
| Tukey's multiple comparisons test | Mean Diff. | 95.00% CI of diff. | Below threshold? | Summary | Adjusted P Value | Mean Diff. | | |
| Europe vs. North America | -0.1854 | -0.3021 to -0.06866 | Yes | *** | 0.0006 | A-B | | |
| Europe vs. All others | -0.09196 | -0.2229 to 0.03894 | No | ns | 0.2248 | A-C | | |
| North America vs. All others | 0.09343 | -0.03053 to 0.2174 | No | ns | 0.1798 | B-C | | |
| Test details | Mean 1 | Mean 2 | Mean Diff. | SE of diff. | n1 | n2 | q | DF |
| Europe vs. North America | 0.693 | 0.8784 | -0.1854 | 0.0496 | 114 | 148 | 5.286 | 352 |
| Europe vs. All others | 0.693 | 0.7849 | -0.09196 | 0.05562 | 114 | 93 | 2.338 | 352 |
| North America vs. All others | 0.8784 | 0.7849 | 0.09343 | 0.05267 | 148 | 93 | 2.509 | 352 |

**Table S43b. corresponding to Figure S6b**
**Turkey's multiple comparison test results. Difference in mentees in North America, Europe, or other continents in having formal mentorship programs at their institution.**
Data analysis for this table includes survey respondents with "0" number of mentors.

| | | | | | | | | |
|---|---|---|---|---|---|---|---|---|
| Number of families | 1 | | | | | | | |
| Number of comparisons per family | 3 | | | | | | | |
| Alpha | 0.05 | | | | | | | |
| Tukey's multiple comparisons test | Mean Diff. | 95.00% CI of diff. | Below threshold? | Summary | Adjusted P Value | Mean Diff. | | |
| Europe vs. North America | -0.2076 | -0.3420 to -0.07321 | Yes | *** | 0.0009 | A-B | | |
| Europe vs. All others | 0.2807 | 0.1296 to 0.4317 | Yes | **** | <0.0001 | A-C | | |
| North America vs. All others | 0.4883 | 0.3449 to 0.6317 | Yes | **** | <0.0001 | B-C | | |
| Test details | Mean 1 | Mean 2 | Mean Diff. | SE of diff. | n1 | n2 | q | DF |

| | | | | | | | | |
|---|---|---|---|---|---|---|---|---|
| Europe vs. North America | 0.5172 | 0.7248 | -0.2076 | 0.0571 | 116 | 149 | 5.142 | 355 |
| Europe vs. All others | 0.5172 | 0.2366 | 0.2807 | 0.06418 | 116 | 93 | 6.185 | 355 |
| North America vs. All others | 0.7248 | 0.2366 | 0.4883 | 0.06094 | 149 | 93 | 11.33 | 355 |

| **Table S43c. corresponding to Figure S6c** **Turkey's multiple comparison test results. Difference in mentees in North America, Europe, or other continents participating in any formal or informal mentorship schemes.** Data analysis for this table includes survey respondents with "0" number of mentors. | | | | | | | | |
|---|---|---|---|---|---|---|---|---|
| Number of families | 1 | | | | | | | |
| Number of comparisons per family | 3 | | | | | | | |
| Alpha | 0.05 | | | | | | | |
| Tukey's multiple comparisons test | Mean Diff. | 95.00% CI of diff. | Below threshold? | Summary | Adjusted P Value | Mean Diff. | | |
| Europe vs. North America | 0.0192 2 | -0.1227 to 0.1612 | No | ns | 0.9456 | A-B | | |
| Europe vs. All others | 0.224 | 0.06387 to 0.3841 | Yes | ** | 0.0031 | A-C | | |
| North America vs. All others | 0.2048 | 0.05241 to 0.3571 | Yes | ** | 0.0048 | B-C | | |
| Test details | Mean 1 | Mean 2 | Mean Diff. | SE of diff. | n1 | n2 | q | DF |
| Europe vs. North America | 0.4957 | 0.4765 | 0.01922 | 0.06031 | 117 | 149 | 0.450 6 | 355 |
| Europe vs. All others | 0.4957 | 0.2717 | 0.224 | 0.06803 | 117 | 92 | 4.656 | 355 |
| North America vs. All others | 0.4765 | 0.2717 | 0.2048 | 0.06474 | 149 | 92 | 4.473 | 355 |

| **Table S44a. corresponding to Figure S7a** **Turkey's multiple comparison test results. Number of mentors by mentees in North America, Europe, or other continents.** Data analysis for this table includes survey respondents with "0" number of mentors. | | | | | | | | |
|---|---|---|---|---|---|---|---|---|
| Number of families | 1 | | | | | | | |
| Number of comparisons per family | 3 | | | | | | | |
| Alpha | 0.05 | | | | | | | |
| Tukey's multiple comparisons test | Mean Diff. | 95.00% CI of diff. | Below threshold? | Summary | Adjusted P Value | Mean Diff. | | |
| Europe vs. North America | -1.183 | -1.601 to -0.7656 | Yes | **** | <0.0001 | A-B | | |
| Europe vs. All others | - 0.0898 8 | -0.5589 to 0.3791 | No | ns | 0.894 | A-C | | |
| North America vs. All others | 1.093 | 0.6466 to 1.540 | Yes | **** | <0.0001 | B-C | | |
| Test details | Mean 1 | Mean 2 | Mean Diff. | SE of diff. | n1 | n2 | q | DF |

| | | | | | | | | |
|---|---|---|---|---|---|---|---|---|
| Europe vs. North America | 1.932 | 3.115 | -1.183 | 0.1775 | 117 | 148 | 9.43 | 355 |
| Europe vs. All others | 1.932 | 2.022 | -0.08988 | 0.1993 | 117 | 93 | 0.637 9 | 355 |
| North America vs. All others | 3.115 | 2.022 | 1.093 | 0.1898 | 148 | 93 | 8.146 | 355 |

| **Table S44b. corresponding to Figure S7b**<br>**Turkey's multiple comparison test results. Mentor-mentee meeting frequency by mentees in North America, Europe, or other continents.**<br>Data analysis for this table includes survey respondents with "0" number of mentors. | | | | | | | | |
|---|---|---|---|---|---|---|---|---|
| Number of families | 1 | | | | | | | |
| Number of comparisons per family | 3 | | | | | | | |
| Alpha | 0.05 | | | | | | | |
| Tukey's multiple comparisons test | Mean Diff. | 95.00% CI of diff. | Below threshold? | Summary | Adjusted P Value | Mean Diff. | | |
| Europe vs. North America | 0.0035 53 | -0.6281 to 0.6352 | No | ns | >0.9999 | A-B | | |
| Europe vs. All others | -0.591 | -1.264 to 0.08160 | No | ns | 0.0975 | A-C | | |
| North America vs. All others | - 0.5946 | -1.188 to - 0.001435 | Yes | * | 0.0493 | B-C | | |
| Test details | Mean 1 | Mean 2 | Mean Diff. | SE of diff. | n1 | n2 | q | DF |
| Europe vs. North America | 2.767 | 2.764 | 0.003553 | 0.2671 | 43 | 72 | 0.018 81 | 165 |
| Europe vs. All others | 2.767 | 3.358 | -0.591 | 0.2844 | 43 | 53 | 2.939 | 165 |
| North America vs. All others | 2.764 | 3.358 | -0.5946 | 0.2508 | 72 | 53 | 3.353 | 165 |

| **Table S44c. corresponding to Figure S7c**<br>**Turkey's multiple comparison test results. Constructiveness of mentorship interactions by mentees in North America, Europe, or other continents.**<br>Data analysis for this table includes survey respondents with "0" number of mentors. | | | | | | |
|---|---|---|---|---|---|---|
| Number of families | 1 | | | | | |
| Number of comparisons per family | 3 | | | | | |
| Alpha | 0.05 | | | | | |
| Tukey's multiple comparisons test | Mean Diff. | 95.00% CI of diff. | Below threshold? | Summary | Adjusted P Value | Mean Diff. |
| Europe vs. North America | - 0.0528 4 | -0.1267 to 0.02107 | No | ns | 0.2133 | A-B |
| Europe vs. All others | - 0.0233 | -0.1056 to 0.05898 | No | ns | 0.7831 | A-C |
| North America vs. All others | 0.0295 4 | -0.04820 to 0.1073 | No | ns | 0.6443 | B-C |

| Test details | Mean 1 | Mean 2 | Mean Diff. | SE of diff. | n1 | n2 | q | DF |
|---|---|---|---|---|---|---|---|---|
| Europe vs. North America | 0.8465 | 0.8993 | -0.05284 | 0.0314 | 114 | 149 | 2.379 | 356 |
| Europe vs. All others | 0.8465 | 0.8698 | -0.0233 | 0.03496 | 114 | 96 | 0.942 5 | 356 |
| North America vs. All others | 0.8993 | 0.8698 | 0.02954 | 0.03303 | 149 | 96 | 1.265 | 356 |

**Table S44d. corresponding to Figure S7d**
**Turkey's multiple comparison test results. Quality of mentor-mentee match (poor-to-excellent) by mentees in North America, Europe, or other continents.**
Data analysis for this table includes survey respondents with "0" number of mentors.

| | | | | | | | | |
|---|---|---|---|---|---|---|---|---|
| Number of families | 1 | | | | | | | |
| Number of comparisons per family | 3 | | | | | | | |
| Alpha | 0.05 | | | | | | | |
| Tukey's multiple comparisons test | Mean Diff. | 95.00% CI of diff. | Below threshold? | Summary | Adjusted P Value | Mean Diff. | | |
| Europe vs. North America | - 0.1464 | -0.4509 to 0.1581 | No | ns | 0.4953 | A-B | | |
| Europe vs. All others | - 0.1458 | -0.4844 to 0.1927 | No | ns | 0.5686 | A-C | | |
| North America vs. All others | 0.0005 63 | -0.3197 to 0.3208 | No | ns | >0.9999 | B-C | | |
| Test details | Mean 1 | Mean 2 | Mean Diff. | SE of diff. | n1 | n2 | q | DF |
| Europe vs. North America | 3.833 | 3.98 | -0.1464 | 0.1294 | 114 | 148 | 1.6 | 355 |
| Europe vs. All others | 3.833 | 3.979 | -0.1458 | 0.1438 | 114 | 96 | 1.434 | 355 |
| North America vs. All others | 3.98 | 3.979 | 0.000563 | 0.1361 | 148 | 96 | 0.005 852 | 355 |

**Table S45a. corresponding to Figure S8a**
**Two-tailed Mann-Whitney test results, Women (column A) versus Men (column B) mentee on their academic satisfaction.**
Data analysis for this table excludes survey respondents with "0" number of mentors.

| | |
|---|---|
| **Mann-Whitney Test** | |
| P value | 0.035 |
| Exact or approximate P value? | Approximate |
| P value summary | * |
| Significantly different (P < 0.05)? | Yes |
| One- or two-tailed P value? | Two-tailed |
| Sum of ranks in column A, B | 26047, 26279 |
| Mann-Whitney U | 11341 |
| **Difference between medians** | |
| Median of column A | 4.000, n=171 |
| Median of column B | 4.000, n=152 |
| Difference: Actual | 0 |
| Difference: Hodges-Lehmann | 0 |

| Table S45b. corresponding to Figure S8b<br>**Two-tailed Mann-Whitney test results, Women (column A) versus Men (column B) mentee on their career optimism.**<br>Data analysis for this table excludes survey respondents with "0" number of mentors. | |
|---|---|
| **Mann-Whitney Test** | |
| P value | 0.0036 |
| Exact or approximate P value? | Approximate |
| P value summary | ** |
| Significantly different (P < 0.05)? | Yes |
| One- or two-tailed P value? | Two-tailed |
| Sum of ranks in column A, B | 25652, 26998 |
| Mann-Whitney U | 10774 |
| **Difference between medians** | |
| Median of column A | 4.000, n=172 |
| Median of column B | 4.000, n=152 |
| Difference: Actual | 0 |
| Difference: Hodges-Lehmann | 0 |

| Table S45c. corresponding to Figure S8c<br>**Two-tailed Mann-Whitney test results, Mentees with Assigned (column A) versus Voluntarily chosen (column B) mentor on their academic satisfaction.**<br>Data analysis for this table excludes survey respondents with "0" number of mentors. | |
|---|---|
| **Mann-Whitney Test** | |
| P value | 0.1469 |
| Exact or approximate P value? | Exact |
| P value summary | ns |
| Significantly different (P < 0.05)? | No |
| One- or two-tailed P value? | Two-tailed |
| Sum of ranks in column A, B | 14320, 38982 |
| Mann-Whitney U | 9855 |
| **Difference between medians** | |
| Median of column A | 4.000, n=94 |
| Median of column B | 4.000, n=232 |
| Difference: Actual | 0 |
| Difference: Hodges-Lehmann | 0 |

| Table S45d. corresponding to Figure S8d<br>**Two-tailed Mann-Whitney test results, Mentees with Assigned (column A) versus Voluntarily chosen (column B) mentor on their career optimism.**<br>Data analysis for this table excludes survey respondents with "0" number of mentors. | |
|---|---|
| **Mann-Whitney Test** | |
| P value | 0.501 |
| Exact or approximate P value? | Exact |
| P value summary | ns |
| Significantly different (P < 0.05)? | No |
| One- or two-tailed P value? | Two-tailed |
| Sum of ranks in column A, B | 14925, 38703 |
| Mann-Whitney U | 10460 |
| **Difference between medians** | |
| Median of column A | 4.000, n=94 |

| | |
|---|---|
| Median of column B | 4.000, n=233 |
| Difference: Actual | 0 |
| Difference: Hodges-Lehmann | 0 |

**Table S45e. corresponding to Figure S8e**
**Turkey's multiple comparison test results. Differences in mentees by region (North America, Europe, or other continents) on their academic satisfaction.** Data analysis for this table includes survey respondents with "0" number of mentors.

| | | | | | | | | |
|---|---|---|---|---|---|---|---|---|
| Number of families | 1 | | | | | | | |
| Number of comparisons per family | 3 | | | | | | | |
| Alpha | 0.05 | | | | | | | |
| Tukey's multiple comparisons test | Mean Diff. | 95.00% CI of diff. | Below threshold? | Summary | Adjusted P Value | Mean Diff. | | |
| Europe vs. North America | -0.2053 | -0.5148 to 0.1042 | No | ns | 0.2637 | A-B | | |
| Europe vs. All others | 0.1393 | -0.1974 to 0.4760 | No | ns | 0.5939 | A-C | | |
| North America vs. All others | 0.3446 | 0.02566 to 0.6635 | Yes | * | 0.0306 | B-C | | |
| Test details | Mean 1 | Mean 2 | Mean Diff. | SE of diff. | n1 | n2 | q | DF |
| Europe vs. North America | 3.733 | 3.938 | -0.2053 | 0.1314 | 101 | 129 | 2.209 | 318 |
| Europe vs. All others | 3.733 | 3.593 | 0.1393 | 0.143 | 101 | 91 | 1.377 | 318 |
| North America vs. All others | 3.938 | 3.593 | 0.3446 | 0.1354 | 129 | 91 | 3.598 | 318 |

**Table S45e. corresponding to Figure S8e**
**Ordinary one-way ANOVA, Differences in mentees by region (North America, Europe, or other continents) on their academic satisfaction.** Data analysis for this table includes survey respondents with "0" number of mentors.

| | |
|---|---|
| **ANOVA summary** | |
| F | 3.376 |
| P value | 0.0354 |
| P value summary | * |
| Significant diff. among means (P < 0.05)? | Yes |
| R squared | 0.02079 |
| **Brown-Forsythe test** | |
| F (DFn, DFd) | 0.1490 (2, 318) |
| P value | 0.8616 |
| P value summary | ns |
| Are SDs significantly different (P < 0.05)? | No |

| Bartlett's test | | | | | |
|---|---|---|---|---|---|
| Bartlett's statistic (corrected) | 0.4749 | | | | |
| P value | 0.7886 | | | | |
| P value summary | ns | | | | |
| Are SDs significantly different (P < 0.05)? | No | | | | |
| **ANOVA table** | **SS** | **DF** | **MS** | **F (DFn, DFd)** | **P value** |
| Treatment (between columns) | 6.608 | 2 | 3.304 | F (2, 318) = 3.376 | P=0.0354 |
| Residual (within columns) | 311.2 | 318 | 0.9787 | | |
| Total | 317.9 | 320 | | | |
| **Data summary** | | | | | |
| Number of treatments (columns) | 3 | | | | |
| Number of values (total) | 321 | | | | |

**Table S45f. corresponding to Figure S8f**
**Turkey's multiple comparison test results. Differences in mentees by region (North America, Europe, or other continents) on their career optimism.** Data analysis for this table includes survey respondents with "0" number of mentors.

| | | | | | | | | |
|---|---|---|---|---|---|---|---|---|
| Number of families | 1 | | | | | | | |
| Number of comparisons per family | 3 | | | | | | | |
| Alpha | 0.05 | | | | | | | |
| Tukey's multiple comparisons test | Mean Diff. | 95.00% CI of diff. | Below threshold? | Summary | Adjusted P Value | Mean Diff. | | |
| Europe vs. North America | -0.4774 | -0.7781 to -0.1766 | Yes | *** | 0.0006 | A-B | | |
| Europe vs. All others | -0.02024 | -0.3480 to 0.3075 | No | ns | 0.9884 | A-C | | |
| North America vs. All others | 0.4571 | 0.1472 to 0.7671 | Yes | ** | 0.0017 | B-C | | |
| Test details | Mean 1 | Mean 2 | Mean Diff. | SE of diff. | n1 | n2 | q | DF |
| Europe vs. North America | 3.584 | 4.062 | -0.4774 | 0.1277 | 101 | 130 | 5.286 | 319 |
| Europe vs. All others | 3.584 | 3.604 | -0.02024 | 0.1392 | 101 | 91 | 0.2056 | 319 |
| North America vs. All others | 4.062 | 3.604 | 0.4571 | 0.1316 | 130 | 91 | 4.912 | 319 |

**Table S45f. corresponding to Figure S8f**
**Ordinary one-way ANOVA, Differences in mentees by region (North America, Europe, or other continents) on their career optimism.** Data analysis for this table includes survey respondents with "0" number of mentors.

| **ANOVA summary** | |
|---|---|
| F | 9.157 |
| P value | 0.0001 |
| P value summary | *** |

| | | | | | |
|---|---|---|---|---|---|
| Significant diff. among means (P < 0.05)? | Yes | | | | |
| R squared | 0.05429 | | | | |
| **Brown-Forsythe test** | | | | | |
| F (DFn, DFd) | 2.415 (2, 319) | | | | |
| P value | 0.091 | | | | |
| P value summary | ns | | | | |
| Are SDs significantly different (P < 0.05)? | No | | | | |
| **Bartlett's test** | | | | | |
| Bartlett's statistic (corrected) | 8.687 | | | | |
| P value | 0.013 | | | | |
| P value summary | * | | | | |
| Are SDs significantly different (P < 0.05)? | Yes | | | | |
| **ANOVA table** | **SS** | **DF** | **MS** | **F (DFn, DFd)** | **P value** |
| Treatment (between columns) | 16.98 | 2 | 8.491 | F (2, 319) = 9.157 | P=0.0001 |
| Residual (within columns) | 295.8 | 319 | 0.9273 | | |
| Total | 312.8 | 321 | | | |
| **Data summary** | | | | | |
| Number of treatments (columns) | 3 | | | | |
| Number of values (total) | 322 | | | | |

**Table S46. What do you like about your mentoring experiences?** Data analysis for this table includes survey respondents with "0" number of mentors.

| theme | sub-theme | comment |
|---|---|---|
| Diversity (more than one mentor, mentors for diverse purposes) span a range of areas of expertise | | -My mentors span a range of areas of expertise, institution, and lived experience.<br>-By having several, I always have someone I can ask a question to.<br>-There are many potential sources for advice.<br>-Get the best of each mentor at academic and personal level. Each mentor I had was important at professional and/or personal level.<br>-They taught different aspects of a scientist job from several point of view. |
| | Mentors inside & outside own institution | -Having mentors inside and outside my institution to provide guidance on a variety of issues related to my lab, the tenure process, my career progression and establishing myself in the field. |

| | | |
|---|---|---|
| Wish I had a mentor | | -I have found them very very useful, in the past I had one through a formal scheme and one I used myself which developed naturally. At present i have none, but would like one. |
| Like Nothing about my mentor | | -My mentor was assigned by the department and has not proved to be very helpful when considering the career. I get a feeling that he is himself struggling with the university process. So far, I have not received any sort of advice for research or career development even when I approached him.<br><br>-I hate the mentorship I receive. It is a mentorship relationship more suitable for exploitation.<br><br>-Not very much. We meet sparingly and he doesn't provide much specific advice on how to be successful.<br><br>-I have not had any substantive mentoring. I have experienced bullying.<br><br>-Actually, I do not like my mentor. My mentor is somebody who was supposed to mentor me through the process of setting up the lab, but turned out to be a someone who tries to take advantage of my funding and my junior position. He was also a trigger for some conflicts between my group and his group. Mentor's way of 'leading' the research team is very dictatorial. And I can't accept that he makes my people feel like they are less than they are. I had to separate myself and my groups from this toxic person. At the moment we are quite alone but much happier. |
| Psychosocial support | Empathy | -Having a sympathetic ear,<br><br>-I feel understood and relieved. |
| | Wholehearted (genuine/sincere) Support & concern | -Support, the transition to Independence is hard and having mentors to ease my way is keeping me sane<br><br>-Mentor has my back.<br><br>-A support system |
| | Cultural sensitivity | -Mentor's cultural sensitivity |
| | Understanding of mentee vulnerability | -Mentor's understanding of mentee vulnerability |
| | Values mentee worth & success/shows respect | -Mentor's valuation of mentee worth & success/shows respect |
| | Encouragement | -Make me feel I can do this. |

| | | |
|---|---|---|
| | Positive affirmation/reassurance | -Self-confidence<br><br>-Someone who tells you you're doing well.<br><br>-Gives me more confidence that I'm on the right track. |
| | Strong belief in mentee potential | -Mentor’s strong belief in mentee potential |
| | Care for well-being | -They genuinely care about me as a person. |
| | Care for mentee success | -Mentor’s devotion<br><br>-My mentor seems invested in my success<br><br>-Mentor puts my career development and progress ahead of his personal interests |
| | Care for progress | -Mentor’s care for progress |
| Positive personal descriptions of Mentor | Open mind | -Mentor’s open mind |
| | Calmness | -Mentor’s calmness |
| | Approachable | -Approachability of my mentor |
| | Active listening | -Active listening |
| | Generous with personal time | These people are generous with their time and gain no personal benefit from the experience. |
| | Profound Trust | -Mentor’s profound trust |
| | Friendliness | - Mentor’s friendliness |
| | Patience | - Mentor’s patience |
| | Availability/Accessibility/reliability/open door policy | -I can go to the when I have questions<br><br>-Meetings as needed<br><br>-I know who I can go to for help<br><br>-I feel able to regularly consult my mentor<br><br>-If I need advice, they are there to help guide me<br><br>-It's nice to know that you can ask questions of someone more senior without feeling like you're bothering them or taking up too much of their time. |

| | | |
|---|---|---|
| | Flexibility | -Mentor's flexibility with time |
| | Honesty & Confidentiality | -The possibility to speak frankly to a colleague.<br><br>-Having raw conversations about our jobs.<br><br>-Honest opinion about my progress and my future career<br><br>-Honest advice<br><br>-the possibility to talk directly and discuss and learn from he<br><br>-I can rely on both for unvarnished, truthful advice.<br><br>-People to ask questions to that won't judge me for asking them! |
| | Camaraderie/Collegiality | -They are very nice.<br><br>-They make me feel safe and welcome; that I belong here and people I trust and admire believe in me and are committed to my success. |
| | Responsiveness | -My mentors are all very responsive and genuine.<br><br>-Easy place to go to try and get answers |
| | Someone to aspire to/ a role model | -My mentors lead by example, which inspires me to follow. |
| | Vision | -Mentor's vision |
| | Quality of Advice | -Quality of Advice |
| | Active contribution to learning process | -Active contribution to learning process |
| Practical help | Help me solve problems | -Mentor help me solve problems |
| | Practical Help | -Mentor help with practical issues |
| | Focused on problem solving for mentee needs | -My needs are considered the most important and conversations are tailored to what I find most useful. |
| Promoting Professional Development | Career (development) Guidance/Advice | -In the past, I have appreciated my mentors' advice about which conferences to attend, which journals to submit to, plus general moral support and encouragement.<br><br>-Solid advice on how to keep my job |

| | | |
|---|---|---|
| | | -My mentors help me to see how to navigate my career track to my goals.<br><br>-Advice and guidance about having a career in science.<br><br>-That ECRs want to get advice about their career trajectories and also their personal lives.<br><br>-The possibility of work in science<br><br>-How to navigate through my career |
| | Help with career development | -Mentor help with career development |
| | Can discuss concerns & plans for future/strategy | -I get good unbiased advice on career moves.<br><br>-Discussion |
| | Open/transparent & Constructive & rigorous Feedback | -Open/honest communication<br><br>-Getting feedback to make sure I'm on track for a successful tenure case.<br><br>-The tenure process, my career progression and establishing myself in the field |
| | Inform of opportunities | -Forward me opportunities whenever they hear about them.<br><br>-helps me learn about new opportunities for advancement. |
| | Alert mentee on opportunities: funding training, awards | - Mentor alert mentee on opportunities: funding training, awards |
| | Helps with Networking/Professional connections | - Mentor helps with Networking/Professional connections |
| | Can gauge and monitor progress | - Can gauge and monitor progress |
| Advice about personal challenges | Advice about challenges difficult decisions & stressful situations | -They help me navigate in issues that was I never prepared for.<br><br>-They have helped me through several difficult situations.<br><br>-Mentor help solves issues<br><br>-Sharing the challenges and getting advice |

| | | |
|---|---|---|
| | | -They give me feedback to help me work through issues I'm facing.<br><br>-About the difficulties I face, more in a perspective fashion.<br><br>-Having a support system and knowing that I can get some input on complex matters when I feel lost. |
| | Advice from experienced senior colleagues | -Senior folks able to advice on problems they are experienced with; they are willing to listen to me venting about stuff. |
| | Advice on unexpected problems | - Advice on unexpected problems |
| | Work-life balance advice | -Facing reality of work-life challenges<br><br>-Advice/ guidance in different aspects of life, career, strategy |
| | Advice on Personal issues | -Having someone to discuss all different types of professional issues with. |
| | Willing to share Wisdom & Experience | -It is nice to talk to someone who understands what you are going through.<br><br>-Others who can anticipate things I need to think about that I'm not aware of (i.e., who can flag "unknown unknowns")<br><br>-Sharing experience<br><br>-Drawing on the wisdoms of others.<br><br>-Telling you things they tried that failed is reassuring somehow.<br><br>-Also hearing about how some struggling with various, often different aspects, is interesting. Helps to realize the heterogeneity in faculty members and more importantly that not everyone is the same and so will not conform to the exact same route to chair/success.<br><br>-Finding career direction from an experienced and successful senior |
| | Help with gender related issues | -Mentor helps with gender related issues. |
| Guidance and advice | Perspective | -Mentors give an additional perspective from an understanding of your research focus combined with their greater experience and knowledge to help guide you with measured direction. Helping at times |

| | | |
|---|---|---|
| | | to keep you on track and focused, and not to get distracted by other research metrics; like journal ranking and reference points.<br><br>-The different perspectives are always very illuminating, often showing that there isn't any 'one' right way to do anything.<br><br>-Perspective: a third party who can help you evaluate your standing and track record.<br><br>-Being offered perspectives based on information otherwise unavailable to me. |
| | Targeted Advice | -My mentors offer targeted advice when asked as well as general perspectives on common challenges. |
| | Informal Advice | -Informal Advice |
| | Help clarify my thinking | -Help clarify my thinking |
| | Help determine time-priorities/how to focus effort | -I value mentor's perspective, helps me prioritize ideas/activities and not waste energy in things that are not relevant (both merely academic and work relationships).<br><br>-Guidance on where and how to focus my efforts; advice on how to achieve what I want to achieve. |
| | Clear Guidance | -Someone who gives one direction and relief.<br><br>-Opportunity to receive valuable guidance.<br><br>-Guidance; openly discuss my failures and problems with my mentors. |
| | Thinking out of the box | -The outside of the box suggestions.<br><br>-Mentoring helped me to think outside the box and find blind spots. |
| Community and Institutional advice | Advice on how things work (navigating institutional bureaucracy, university administration, politics & sexism, funding & hiring decisions) | -Getting insight into the functioning of academic units and how to play the game more effectively without being destroyed by the never-ending demands.<br><br>-Individuals willing to help me to solve problems from navigating the institutional environment<br><br>-Getting different perspectives, |
| | Worldview & overview on academic careers, | -That they have "the whole picture" in mind (but sometimes they do not share it adequately, in a clear easily understood way) |

| | | |
|---|---|---|
| | sharing thoughts, reflecting | -Transfer of experience on how the scientific community work. |
| Funding | Grant Writing & Application Advice | -Looking over grant applications has been super helpful |
| | How Funding Agencies work | -Transfer of experience on how the funding agencies work. |
| Academic Advice | Advice on lab management & hiring trainees | -Managing lab group members<br><br>-Managing my own group<br><br>-Guidance on a variety of issues related to my lab |
| | Advice on setting up a lab | -Advice on setting up a lab |
| | Advice/Approach on mentoring trainees | -Helping navigate complex issues in mentoring<br><br>-Help on how to best mentor trainees.<br><br>-Mentor taught me (sometimes showing what I shouldn't do) the best way to contribute to the professional and personal growth of people that work with me |
| | Advice on writing papers/preparing manuscripts | -Advice on writing papers/preparing manuscripts |
| | Guidance on how to communicate my work better | -Guidance on how to best spin our research to gain attention. |
| | Help with how to collaborate | -Periodical evaluation of self-growth and opportunity to collaborate! |
| | Service advice | -Teaching/service advice |
| | Stimulate scientifically | -I like when they actually stimulate me to want to do my job better. |
| | Sharing Knowledge | -Knowledge transfer opportunities<br><br>-Learn from others<br><br>-Each of them fills a gap of knowledge I require |
| | Help on focus on short-term & long-term goals | -Helped me to give a clear focus to my research |
| | Advice based on their research experience | -Their expertise<br><br>-Advice on experiments |

| | | |
|---|---|---|
| | Guidance to elevate mentee work | -Guidance to elevate my work |
| | Focused Scientific support | -Advice is specific to the research task and is intuitive<br><br>-Research questions<br><br>-My mentors provide complementary advice that helps me to see how I can improve my science. |
| | Sounding board of ideas & directions/being able to brainstorm/bounce off ideas/offer new ideas | -to have someone to bounce plans off, sanity-check ideas, and to draw on their experience<br><br>-New ideas or ways to do the things<br><br>-The ability to test ideas in a low-risk setting. |
| Learn new skills | Transfer of Skills | - Transfer of Skills |
| | Learning new skills from mentor | Learning new skills from mentor |

**Table S47. Responses to "If you answered "No" or "Sometimes", what type of mentoring relationship were you looking for? What were you hoping to achieve with your mentor?** Data analysis for this table includes survey respondents with "0" number of mentors.

| Theme | sub-theme | comment |
|---|---|---|
| Tenured faculty also need mentors | | My mentor is the chair of my department. He's not a mentor in any official capacity but he's the only person I've really gotten advice from since I arrived. The junior faculty all have mentorship committees but apparently there is no such thing when folks arrive with tenure. |
| Promoting Professional Development | Focused Career Guidance/Advice | -Helping navigate career trajectory in terms of sharing opportunities.<br><br>-Provide advice about career options or lack thereof.<br><br>-Guidance, constructive criticism.<br><br>-Mentor would have given me some guidance to build up my career.<br><br>-Appropriate support and guidance.<br><br>-I want more structured support in developing my research profile - my previous roles (3 years' worth) have not provided that. My mentor didn't choose to mentor me (though she agreed without any problem - but it wasn't exactly her |

| | | |
|---|---|---|
| | | choice) and I'm not sure she feels fully confident in how to mentor (yet).<br><br>-I would have liked to have some strategic tips regarding the mid-term planning of my career, both from the point of view of grant applications and in deciding how to expand vs focus the research interests of my group.<br><br>-I was looking for guidance on how to navigate the pitfalls of starting my own project. At the very least feedback on my (funded) project would have been nice. Instead, my mentor expected me to manage my project and execute technical work for her project (in a different area). |
| | Advocate | -Also wished the mentor would act as an advocate.<br><br>-I was hoping that mentor looked after my interests as well, not just own |
| Relationship dynamics | Not a competitor | rather than seeing me as a competitor.<br><br>-I would have liked to feel that my "mentor" selflessly would promote my chances at success. It is sometimes true for small things. But often I feel he sees me as a sometime competitor for department resources and also undermines me on occasion, which is the opposite of supportive.<br><br>-Can’t be sure if he is giving opinions/directions that are in my best interest, as opposed to sometimes thinking more of the benefit of himself or his favorite successor. |
| | Regular meetings/progress checks | -He gives great advice but as I’ve learned more (closed the gap between us), and I’ve started asking more complicated questions, sometimes his advice has not been as good.<br>It’s a little too sporadic. It would be more helpful if it were more often. |
| | Collegiality/great relationship based on trust & respect | -Constructive relationship with compromise, time for discussion, without lost time. more velocity in the feedback.<br><br>-They were at the beginning, but lately we almost stopped talking to each other.<br><br>-A relationship in which the problems that arise at work are accepted, and we work to solve them instead of assuming that they should not exist and it is my fault. Let there be help and work valuation. One in which there is trust, in which it is recognized that the decisions were taken together. One in which rest time is respected. |
| Guidance and advice | Constructive<br>Positive<br>Honest/Sincere<br>Feedback & Communication | -Positive, with constructive and honest advice.<br><br>-I need more feedback. |

| | | |
|---|---|---|
| | | -I would like my mentor would be sincere with the expectations instead of telling me something that he knows it is not real.<br><br>-Honest, constructive feedback - I sometimes feel he is very vague and afraid to direct me in a certain way.<br><br>-A bit more of understanding, being more open and a more fluid communication. |
| Emotional support | More empathetic | -It would be great to have a mentor on site and someone I could talk to in person.<br><br>-To get more understanding...the mentor should understand what I'm going through.<br><br>-A person more empathic, more liberal, a person who do not think you are inoperative without a real reason, a person who gives you courage to became a scientist. |
| | Consistent Support | -Someone more consistently supportive |
| | Be Available/Care | -I effectively have no relationship with my mentor. I have met them once in 4 years.<br><br>-More interested in my career. Felt more like a research meeting than mentoring one.<br><br>-My mentor is super hands off which makes me feel like I can only go to him with important or pressing questions/matters. |
| | Values my research | -I was hoping for encouragement but we don't particularly get on - and I don't feel she understands or values my research. |
| | Encouragement/Empowerment/positive | -Encourages creativity and innovation rather than just more papers.<br><br>-An empowerment, they should ask what I want to learn and give constructive suggestion on how to achieve it.<br><br>- Got along with early career faculty |
| Scientific Advice | Professional Respect/values my opinions | -Though, he is a good person and that has made the interactions more relaxed. He does not make me feel that my concerns are unimportant.<br><br>-Cordial productive interactions instead of aggression<br><br>-To admire the person that will share and guide me through his critical thinking, rather than not respecting me as a human being neither a scientist. |

| | | |
|---|---|---|
| | | -We also tried mentoring on the project progress, but it was more like he was diminishing my ideas and making me feel like I don't deserve to be where I am.<br><br>-I was hoping to brainstorm together, instead of them forcing their ideas on me, even if they may not 100% agree with the direction I have in mind. |
| | Scientific Support | -I would expect more guidance related to scientific content |
| | Invested in my success | -Someone with enough time to provide contextual advice, not just standard aphorisms that apply to all.<br><br>-I expected that my mentor was more involved and compromised with my research projects and guided me in a closed way. I think sometimes you need some freedom but in other cases it's necessary to be supported. A fluent exchange of ideas is vital. |
| | Knowledgeable | I was hoping for more constructive guidance, but my original mentor is entirely clueless about modern careers.<br><br>-They don't need to be my area, but that is an advantage in terms of accessing networks and specific knowledge<br><br>-I hoped for some suggestions that I had not come up with yet<br><br>-I think there are sometimes questions she doesn't have the answer from.<br><br>-Understanding my specific expertise. Was hoping to be incorporated into projects.<br><br>-It would be useful sometimes to have someone closer to my own field. I am a biochemist and my mentor is an engineer/analytical chemist.<br><br>-mentorship in specific scientific approaches is somewhat lacking.<br><br>-I think we might be a bit too far apart in terms of research area/interest.<br><br>-Present (career) mentor does not work in the same field |
| Community and Institutional advice | Workplace advice | -I was hoping for someone that spoke to all necessary components to be successful in my department. Then they would ask enough questions to determine where I need the most help.<br><br>-I was hoping my mentor will help me to understand how the reality works in new home institution Working as a PI is not easy. And help to understand home institution is crucial. |

| | | |
|---|---|---|
| Advice about personal challenges | Advice on Work-life balance | -Work-life balance, less like a boss, someone who celebrates and encourages success and growth above and beyond publishing papers. |
| | Gender related issues | -I don't have anyone directly in my field, or who are women, who can give more specific mentoring on my field or women-specific questions I have. |
| Personal characteristics | Active listening | -Relationship where mentors listen a lot and ask inquisitive/thought-provoking questions<br><br>-Focused advice. |
| | Visionary/Role model | -I would like my institution to have a serious vision of science development. I feel that as scientists, we are totally abandoned and have zero support.<br><br>-To admire the person that will share and guide me. |

| **Table S48. Overall to what extent do you feel that your current mentor is meeting your expectations? Please explain your answer to the previous question.** | | |
|---|---|---|
| Lack of Mentoring | Mentoring is neglected | -Feel pretty much on my own. Mentoring in department is not negative, just benign neglect. |
| | No institutional mentors | -Being a distant relationship, I have not met my mentor face to face |
| | No institutional mentoring program | -Our university doesn't have a formal mentoring program so I had a very low bar as to what to expect from a mentor. |
| | No institutional mentors | -I have an assigned mentor at my university but they are not helpful or trustworthy. My mentors off campus are fantastic, but so far away that it is hard to get advice or feel supported sometimes |
| | No formal mentors | -No formal mentorship means no explicit expectations. Ongoing support from a more senior colleague is really valuable though.<br><br>-My mentor is an informal relationship; they were not assigned to me but have mentored me of their own accord. |
| | No mentor | -never had a mentor<br><br>-my mentor is not really mentoring me: he was assigned to be my mentor, but I have the impression that he does not understand what sort of support I need.<br><br>-currently I don’t have one actively.<br>-would have stop the mentoring if not |

| | | |
|---|---|---|
| | Need for formal/Institutional mentors | -this mentor was not assigned, it is purely informal<br><br>-My mentor was not assigned to me, but came from the same lab I did a postdoc in. He did his postdoc in the same lab 15 years earlier. We had that in common so when I came to my institution, he reached out to help me. He has looked out for me ever since.<br><br>-My mentor is not officially assigned but is really just someone who has volunteered to help with grant applications.<br><br>-Difficult. Not a formal mentor. My postdoc advisor has acted as my unofficial mentor and has helped much more than any mentor at my current institution. They have been a huge help, but admittedly someone one the same campus might have been more helpful at times.<br><br>-Wish it was more formal so that I didn't feel like I was bothering them. Hopefully then the multiple formal interactions would also lead to an informal relationship. |
| Need Multiple mentors | | -I receive excellent mentoring from multiple sources and the person I consider my primary mentor has been spectacular.<br><br>-The mentor I have in mind is also Department Chair. He provides the professional mentoring I need sometimes.<br><br>-She is fantastic, but there are some things that she is not the right person for. That's why I have multiple mentors.<br><br>-I answered the above questions really in regards to a variety of folks I have looked to as mentors. No one is perfect in any one thing. Sometimes you have to figure out what a person is good at to understand who would be the best person to ask for advice on certain things.<br><br>-I have several mentors from multiple programs/departments<br><br>-My actual mentor is way better than my previous ones when I worked as a postdoc<br>I have chosen a group of mentors, each for specific skills as the assigned senior faculty mentor wasn't very helpful. Each mentor does a great job at their task, but I don't expect one to advise effectively on all topics.<br><br>-I have many mentors - it's not one mentor - Not everyone has all qualities i am looking for - So i assembled a team for myself prior to moving to this independent job. I have equal number of men and women mentors, locally, previous institutions and also other institutions within the area. I approach each mentor depending on the question/problem i am facing. |

| | | |
|---|---|---|
| | | -My mentors are 2 senior males and a senior female who provide complementary information about how to advance my career. |
| Informal mentors are necessary | | -My mentor is great for strategic questions. However, I would prefer a mentor with the same gender as me and outside my line management hierarchy.<br><br>-I have a formal mentorship relationship that only makes sure I'm checking boxes for tenure. |
| General Description of mentorship | So-so mentor/barely acceptable mentorship | -My mentor has been good, not outstanding, and occasionally disappointing.<br><br>-Many things can be better<br><br>-It is very rare that it hardly makes a difference. Still better than nothing though.<br><br>-It is neutral because sometimes I feel positive and sometimes it is opposite<br><br>-Things are good in some areas and could be improved in others.<br><br>-In the middle because is not white or black, but a grey depending on the particular aspect.<br><br>-We are persons with good and bad things and if not trained this is reflected in your mentorship. Also, the context and the pressures funding/publishing matters a lot.<br><br>-She is fantastic, but there are some things that she is not the right person for. That's why I have multiple mentors.<br><br>-My mentor is also a fairly young hirer so has limited time/opportunity to help me with certain things but I'm very grateful for the guidance I do get.<br><br>-Discussions with my mentor are often helpful, but not always through what is said, but through learning to understand ”how things work” at my current institution.<br><br>-moderate<br><br>-He is very mediocre<br><br>-my mentor is a good mentor that need help to better create relationship with their people.<br><br>Relatively low effort interactions |

| | | |
|---|---|---|
| | Former mentors horrible | The best mentor I've had, previous mentors have been worse than dirt.<br><br>-Far more positive experience than with previous mentor.<br><br>-I had horrible former mentorship. I search literally around the world for a mentor with high soft abilities. My mentor always privileges well-being over productivity. |
| | Great Mentor | -I was incredibly lucky to have a fantastic grad advisor; our formal obligations to each other ended with my graduation but his support and mentorship did not. I am incredibly grateful and lucky to have him there every step of the way, and know I can always rely on him for advice and support. I'm getting teary-eyed a little thinking about it. |
| Advice on difficult & unexpected situation | | -Provides lots of advice for unique situations that come up that I haven't handled before. |
| Challenges me | | He challenges on my pre-conceptions personal and scientific. |
| Honest feedback | | He is not afraid to tell me when something can be done better. |

| | | |
|---|---|---|
| Careers (positive) | Promote ECR PIs work | -Mentor makes sure to highlight my success to the senior leadership. |
| | Provides sponsorship | -My mentor is also my sponsor and actively supports my career development through discussions about research/career strategy, by recognizing my contributions to research, and by introducing me to senior research leaders. Mentor also provide critical and constructive feedback for research grants/papers. |
| | Helps with networking | -Mentor connects me with people who can help, |
| | Helps with career development | -Suggests career development books, workshops.<br><br>-I expect advice on scientific issues and professional issues in general,<br><br>-Helping and guiding me throughout my career as an early researcher.<br><br>-provides me with ample opportunities to promote my career. |
| Careers (to be improved) | No active endorsement/sponsorship | Gives advice in very severe situations; not active encouragement or "sponsorship". |

| | | |
|---|---|---|
| | No focused career advice | He provided a safe environment, however, career wise there is not much guidance. |
| | No effective advice | -They are very good at pointing what I have achieved and what I need to achieve, but do not provide good constructive advice on how to achieve this, or help develop a career plan. |
| | Need help with career advice | -Not helping navigate career or professional development.<br><br>-Less direct interactions intended to help my career.<br><br>-Mentor told me we would work together to get our goals and this is not happening, I am working and learning by myself. |
| | No networking/speaking opportunities | -As my mentor isn't very senior there is limited opportunities for her to put me forward for speaking slot.<br><br>-I think he could do more to promote me in the international scientific scene. |
| Emotional support (positive) | Genuine Support | -Regardless of his personal biases, my mentor could guide me through and help me identify opportunities for my own work. |
| | Provides Encouragement | -Provides encouragement.<br><br>-My mentor has been key in my career since I was still a PhD student and has supported me with a constant attitude of "let's do it, everything is possible", and teaching me self-confidence. |
| | Values mentees | -He values my contributions and my qualities.<br><br>-my contributions were barely recognized in publications.<br><br>-At the personal level I feel safe and professionally I feel valued.<br><br>-I hope more empowerment not exploitation to do all their project without any acknowledgment |
| | Supports my research | -My mentor supports research |
| | Very supportive/helpful | -I chose my mentor based on his attributes in other ways and he is very supportive in connecting me with the clinical aspects related to the basic research I do.<br><br>-My mentors have tried to remove any barriers to my progress and have been fair in any advice/critiques.<br><br>-Without their support I would have already left academia.<br><br>-Feel good about my mentor's support |

| | | |
|---|---|---|
| | | -Always supportive; a cheerleader.<br><br>-Always on hand to lend advice and support my career.<br><br>-I think my mentor is very similar to me personality-wise, and they have gone above and beyond to help me recover from a really bad postdoctoral experience.<br><br>-We have differences but overall, he looks after me.<br><br>-Although my current mentor is from a different field of Expertise, he gives me the support I need to develop my career.<br><br>-Always willing to help/listen and gives great advice.<br><br>-He is very knowledgeable yet humble, he is so appreciative and works very hard to make academic life and work easy for me, which I did not expect.<br><br>-Will take time out of his busy schedule to give detailed feedback on grants, advise on career moves and strategies, with no formal recognition or benefit to him.<br><br>-Simply, my mentor provides the support I need.<br><br>-A supportive figure when needed. There have been times when she has mediated when events have transpired.<br><br>-I have low expectations, but this mentor is very encouraging and supportive.<br><br>-My mentor is always available to help me in any situation, big or small, professional or personal (like needing help getting childcare so I can work) to facilitate my research goals.<br><br>-My mentor has gone beyond was expected with help even outside of hours.<br><br>-Current mentor is very helpful with many aspects of my life and career.<br><br>-Although he works in a slightly different field, mentor is very available and empathic.<br><br>-I am a research assistant professor in one department (so I can apply as a PI for grants) and a postdoctoral fellow in another. Both departments are very supportive.<br><br>-My mentor is just excellent. Mentor has extensive experience, and has mentored many people. Mentor looks out for me, forwards me every opportunity that is out there, |

| | | |
|---|---|---|
| | | and is very helpful when I apply for any opportunity. Mentor tries their best to ensure I achieve my goals.<br><br>-I have found my experience totally supportive and having not previously had a "proper" mentor before I am extremely happy.<br><br>-I had great encouragement and support.<br>-Supports my ideas, gives me advice about career development and grant writing.<br><br>-This particular mentor is a mentor that I have had since grad school and has guided me through many aspects of my career development and job transitions. It took me years to realize how fortunate I was compared to others. |
| | Very satisfied with advice | -Because this is considered a loose relationship without any commitments, I am very satisfied with the supports and advices I have gotten so far.<br><br>-Overall great, I'm still not 100% forthcoming about some struggles.<br><br>-so far so good<br><br>-Mentor is excellent and compassionate and dedicate significant time to the relationship. |
| Emotional support (negative) | Lacks cultural sensitivity | -My mentor does not understand his privilege or how to navigate in a changing academic culture. He also does not understand how our field is changing with funding, methods, and systems thinking. |
| | No respect | -We disagreed on certain issues, and since then I do not feel the mutual respect is sufficient to have a good mentoring relationship.<br><br>-There is a basic lack of respect and communication. |
| | Not respectful | -Due to a lot of uncalled for aggression, there's been a lot of loss of time and productivity. Aggravated my depression and anxiety issues.<br><br>-We have distanced each other and only talk if necessary. Mentor is not satisfied with my work and with the fact that I had two children in the past 4 years. I disagree in how mentor led the project I'm working on.<br><br>-I have previously had mostly positive relationships with mentors, and so expected the same here. I am regularly helped by my current one when required, but feel that all mentors should be expected to do so. |

| | | |
|---|---|---|
| | Not Invested in mentee success | -My mentor does not care about my own projects or career. Instead, she took advantage of the forced relationship for me to conduct experiments for her own work.<br><br>-However, since he is retiring very soon his motivation and energy to apply for grants, write projects and publish is decreasing.<br><br>-My mentor does not seem to care about my success neither the project's that we share. |
| | Mentors need to be effective/mentor needs to be invested | -My five mentors run the gamut such that the average is a 3.<br><br>-Not much interest and not many skills in mentoring.<br><br>-I effectively have no relationship with my mentor having only met them once. |
| | No genuine support | -but I have the impression that he does not understand what sort of support I need.<br><br>-the support is poor<br><br>-My mentor can be very negative and often 'forgets' to include or support me at key moments.<br><br>-I expected our team leader to be a mentor for the younger group members. Mentor expects us to emerge on our own by not getting in our way (consciously).<br><br>-My mentor is a senior person in our dept; gives advice, but only when asked.<br><br>-My mentor is very good in giving guidelines for research excellence and to obtain research funds however, he fails to ensure a working environment free of harassment and he is sometimes biased. This directly affects my development as an independent scientist.<br><br>-I feel that he is present and helpful when I need it but doesn't go out of his way to provide guidance.<br><br>-He is great to talk to and use as a sounding board. He gives advice, but there often is less follow through than what he says he will do.<br><br>-My mentor makes good suggestions, but is generally hands off, which encourages my independence.<br><br>-Assists with specific questions, provides opinion when asked, but does not go above and beyond to help promote my career. Though I expect this as everyone is busy with their own research and career plans. |

| | | |
|---|---|---|
| | | -We have limited interaction, on very focused questions. It is not a general relationship.<br><br>-I know that they have my back, they make sure to highlight my success to the senior leadership. However, their support sometimes feels superficial. We all are very busy, and it takes lots of effort to listen to each other, and truly hear each other's needs. Overall, I am grateful they spend the time they do for mentoring me. But I am also aware of the shortcomings.<br><br>-My mentor can be great but sometimes excludes me from key organizations and has in the past kept me off grants and papers that I authored so there is a weird tension because we both work in the same field. |
| Personal descriptions of Mentor (positive) | Is available | -Mentor is available for me.<br><br>-Readily accessible and provides practical and pragmatic advice.<br><br>-Investment of time and efforts to improve. |
| | Mentor is generous | -My mentor is very helpful and willing to share. |
| | Mentor adapts communication style | -I like the fact that my mentor adapts to my personality and to my view of science. |
| | Mentor does not impose views/allows independence | -Mentor does not impose his own view (more experienced one, of course), but he lets me try and explore all the possibilities before giving me his opinion. He also underlines that his opinions are his own ones, which might be not suitable for me. |
| | Similar in personality | -I think my mentor is very similar to me personality-wise, and they have gone above and beyond to help me recover from a really bad postdoctoral experience.<br><br>-Same personal background as me. Also, is my current boss (not formal mentor program) but is similar. |
| Personal characteristics (negative) | Not available | -Sometimes difficult to get hold of mentor but when we do meet, they are excellent.<br><br>-We don't meet frequently enough to make progress on the issues I face more urgently<br><br>-Great relationship when meeting but availability not always good, initiative almost exclusively mine.<br><br>-Our interactions are sporadic but always helpful.<br><br>-His lack of time |

| | | |
|---|---|---|
| | | -My contact with my mentor is only by e-mail. thus, our interaction is restricted to advice for specific agendas and thus my be restricted in some ways.<br><br>-I am generally happy but would appreciate more availability.<br><br>-Mentor definitely sees the whole picture and is very precise on how to get to the final goal (a grant or getting the results published in the highest quality journals) but has a really poor mentoring relationship with me. Mentor only has meetings with me if I ask (beg) for them, though his desk is at the same distance from mine as the distance I walk to his. We can spend months without talking unless I ask to. |
| | No time | -He is excellent and also super busy so we don't usually have the time to go as in depth as possible.<br><br>-More could be done to engage regularly.<br><br>-lack of time<br><br>-I'm very appreciative, but she is super busy and our areas are not well aligned, so there might be a better mentor out there for me. |
| | Not to impose opinions | I often feel that my mentor tells me what they would do in a situation, which isn't something I'm always comfortable with.<br><br>-It’s sometimes unclear whether the relationship is of a mentor-mentee or a boss-employee<br><br>-Negative self-observed person who just wants to the boss.<br><br>-My mentor is self-aware, understands the constraints and challenges of the system and helps me think creatively on how to overcome these. He does not judge, nor impose ideas. |

| | | |
|---|---|---|
| Personal descriptions of Mentor (negative) | Lacks active listening | -There's a lot of potential, but also some gaps- differences in communication style, etc. that sometimes makes it difficult. But he does not actively seek out mentoring opportunities or provide guidance unasked for.<br><br>-Mentor is a very difficult person to talk to and does not listen. |
| | Mentor training is vital | This is one of my informal mentors. She is amazing. But she has also received training on mentoring. Training is key. And buy-in.<br><br>-Not many skills in mentoring |

| | | |
|---|---|---|
| Relationship (negative) | Mis-communications | -My mentor communicates with me a lot, however, there are a lot of misunderstandings and changes in opinion. |
| | Harmful | -Mentor is self-centered and prevents growth |
| Academic advice (positive) | Advice on managing lab | -Mentor can sometimes be very helpful with advice on managing staff or situations |
| | Help with facilitating collaborations | -Mentor has helped by facilitating collaborations. |
| | Offer new ideas | -Thus far she confirmed my ideas and suggestions, but nothing new or with her initiative |
| | Helps scientifically | -My current mentor helps me scientifically and will go out of his way to help me when I express a problem. |
| Academic advice (negative) | Mentors should help with building independent research project | -My experience is such that I am almost a self-mentored person my supervisor gave me space to develop.<br><br>-While my mentor cheers for me, and gives me lots of positive feedback, unfortunately they have an attitude that feels hierarchical ("You are too inexperienced, let me tell you what is best for you"). They sometimes fail to listen to my needs, and project their experience on me. |
| | Mentors should help with transitions | -My current mentor is interested in helping me overcome all the transition difficulties like helping me choose the best collaborations, finding resources to help getting time to focus on the research and really helping me finding my own projects to build my independent research. |
| | Mentors should help with finding collaborators | -I expected to be more accompanied in the academic and scientific field. I wished a major effort from he/she to exploit my academic potential, to stablish collaborations, to produce better scientific projects and reports. |
| | Need mentoring on how to mentor my own trainees | I need more effective communication and more training about how to become a mentor. |
| | Discuss aspects of being a PI | I have been assigned a mentor through a professional society, as such my mentor is strictly professional, and does not consider it part of the mentorship to discuss other aspects of being a PI. |
| | Need focused/concrete research advice | -I wish my mentor were able to provide me with more concrete advice about priorities in my lab and in my science.<br><br>-Mentor can’t offer research mentorship but mentor is very nice. |
| | Need mentor in the same field | -My mentor is great, however in a very different field (clinician, while I do basic science with a health focus) so I |

| | | |
|---|---|---|
| | | don't have a mentor I can discuss science (or lab-based science) with.<br><br>-Sometimes you are assigned a mentor who does not understand what your research is, and gives limited support.<br><br>-He supports me as best he can. Again, we have very little research area overlap so that can be limiting.<br><br>-I feel the experience would have been even better if my mentor worked in the same field as our lab (currently not really an option in our institution).<br><br>-I wish I had a committee of mentors that were closer to my field.<br><br>-My mentor is an ophthalmologist dedicated to the clinic and I am a researcher dedicated to neuroscience. I think he meet my expectations considering our differences, but I would love to have a mentor who could mentor me in my career as a researcher.<br><br>-My mentor isn't in the same field of study as me so all help is limited to how to advance in my career at my Institution. Unfortunately, I don't have a mentor that helps me advance in my field of study so that's up to me alone. |
| Funding | Helps with grant writing | -Grant revisions<br><br>-My mentor has been helpful in reviewing my grants, putting my name forward for a review, in short more of a champion, less advice. |
| | Mentor should help in finding grants | -Funding is a big obstacle, and there is no active position to promote young PIs to search for international grants. |
| Community and Institutional advice | Need to know how academia works today | -Not sure any of them recognize the differences faced in academia by different generations. |
| | Helpful in navigating institutional politics. | -So far, he has been useful in helping me navigate the politics at my school and university but less helpful in hiring and training personnel. |

**Table S49. Other Comments about the survey or your experience that was not addressed in the questions?** Data analysis for this table includes survey respondents with "0" number of mentors.

| Theme | sub-theme | Comment |
|---|---|---|
| No mentors | | -As I do not have a mentor. I was briefly involved in a group mentoring scheme but this was not successful or continued. I have not had the opportunity to have another mentor but wish it was possible, as I feel my progression and |

| | | |
|---|---|---|
| | | promotion chances have been limited by a lack of support and guidance from more experienced members of staff.<br><br>-Based on my PhD, postdoc and junior PI experience, a good mentor (or even an average mentor, who knows what mentoring means at all) is something like a unicorn. Only exists in fairytales.<br><br>-It's very uncommon to have a mentor in the Spanish system, which I think it's unfortunate. Specially an independent person acting as a mentor, not a former supervisor keeping you under his/her umbrella… |
| | Mid-career Faculty also need mentors | -I mentor lots or PhD students and postdocs and tenure-track fellows, informally and as part of a mentoring program, but I have never been offered any formal mentoring from my institute.<br><br>-Other than brief interactions (maybe 2-3 people, less than one hour/year for specific questions) my only mentors are the internet, twitter, and midcareer PI Slack.<br><br>-We don't have mentors ourselves and now have unrealistic expectations put on ourselves by our graduate students. So, in order to alleviate graduate student stress and protect their work life balance and mental health - I feel I have to do it all myself. |
| | Minorities Need mentors | -As a female, lack of mentorship and lack of academic support from senior researchers is the number one contributor to the leaky pipeline.<br><br>-Opportunities for women to transition from postdoc stage to independent investigator stage. |
| Mentorship team | | -Mentorship comes in many ways: on-site vs off-campus<br><br>-I benefit from supportive professors externally who have been encouraging, but internally it seems to be expected that you wouldn't benefit from this once you have a tenured/associate professor position, even though your funding opportunities are reduced compared with junior colleagues.<br><br>-Different kinds of mentors for different issues are important!<br><br>-I have different mentors for different needs. For instance, I have a separate mentor for career and work/life balance mentor.<br><br>- I get different things from different people as my mentors and have highly different experiences with all of them.<br><br>-I am lucky to have several mentors (both seniors and peers) and I get different needs met via different mentors. I chose to |

| | | |
|---|---|---|
| | | focus on one of my closest institutional mentors here, thinking that their shortcomings are more important to highlight. |
| | Provide informal & Formal Mentorship | -I think it should be informal, and diverse, with many different people, on a need- to -know basis.<br><br>-My institution has a formal mentor program that sucks. If I did not seek my own mentor out, I would not have been as happy. |
| Institutions need to offer Mentorship & training for Mentor | | -My institution does not have a mentor program, so I feel fortunate to have one, however I think should an essential part of every institution, but even more in the academia.<br><br>-My school is too small to have anyone who works in my specific field. I basically had to mentor myself in terms of my research trajectory. I just got tenure and during that entire time not a single tenured faculty member at my school ever asked me to be on a grant or paper with them. I had to do everything myself and I'm admittedly bitter about it. I am trying to be more supportive of the current junior faculty.<br><br>-I find it awkward that I can't change my mentor without offending a colleague (since my mentor is a current close colleague). I think it's impossible for academics to be expected to be good mentors without training.<br><br>-I have just sign ed up to a formal mentoring scheme so well see how that goes. |
| Mentoring Relationships | Former mentors are valuable | -I had a great mentorship experience during my PhD. Despite my current mentorship being unhelpful I still value and receive mentorship via email from my PhD supervisor.<br><br>-My mentor is my former PhD advisor. We kept our relationship going over 20 years. |
| | Negative mentoring experiences | -My PhD mentor was horrible. I learned how to choose a mentor based on that extremely negative experience. |
| | Professional Jealousy & Competition | -Lots of professional jealousy rather than generosity among peers, perhaps because of a situation of penury. Power and political games mess with mentee head and distract them from effective work...So I feel fully unsupported and there is no effective institutional effort to organize mentoring at any level, except for a good workshop I attended some years ago about mentoring predoctoral positions. Nothing like that for upper career levels and a lot of unsavory character traits emerge among peers. |
| | No real advice/Poor mentorship | -There is a lot of pressure on younger PIs to get grants/fellowships and papers, but often not the support to help you achieve this. Mentors and performance reviewers are very quick to tell you to get more 4* papers and money |

| | | |
|---|---|---|
| | | but not good at helping you achieve this. Everyone knows you need high quality papers and grants, so this information is redundant and useless. One of my mentors told me to drop all projects that were not going to make it into the top shelf journals, with that view I might as well give up now. Often new PIs are just handed keys to an office and are expected to thrive and pump out 4* papers and grants, which may or may not happen.<br><br>-I wish I had a mentor.<br><br>-Mentoring in Spain, as defined in this survey, is non-existent in my opinion. The system is not prepared for this kind of career guidance. There is no culture of mentoring, but of competition among all researchers (senior vs junior many times in the same call) for a meagre amount of money. "Jungle-type" of research culture. Ego-based relationships. No cooperation or mentoring in my experience<br><br>-I am not sure I have ever had a supporting mentorship. There has been no formal mentoring system previously and I have been “misled” by previous from my supervisors which have not been too encouraging or in the best interest for my career it theirs. I have gotten much more help and support to grow academically from peers, which is good but not as efficient maybe, not the same thing at least. It would be fun to know how many mentors people have had in the past and the fraction of those they have found good/supportive. |
| Emotional support | Mentors not invested | -I was given a mentor for the first three years after my PhD, but not any longer. The mentor changed every year and most of them showed zero interest in what I was doing. |
| | Need for Respect & Collegiality | -There’s no formal mentoring in my college mostly because there’s not much need - good hires with a low tenure bar and collegial environment where informal mentoring is readily available. Yeah, for real. It exists.<br><br>-hostile untrustworthy mentors than undermine junior faculty |
| Personal characteristics of mentor | Allow independence | -In that experience my mentor allowed for personal growth and encouraged new ideas. |
| | No Feedback | -Mentors never read anything I requested feedback on. |
| Mentors are vital for success | | -I value mentorship as one of the most important factors in long-term success. |
| Academic advice | Knowledgeable Mentors | -The qualification of the mentors<br><br>-Previous mentoring from a scientific society paired me with a retired prof who had really had his whole career in a different era and while he as supportive he was unable to provide advice on many of the thorny issues I faced in a more |

| | | |
|---|---|---|
| | | modern context (publish or perish; move a continent away from your partner for a job etc). |
| | No advice on managing lab & mentoring own mentees | -There is no real training on how to manage a group and no support when things go wrong. |
